# AMPK Regulates Phagophore-to-Autophagosome Maturation

**DOI:** 10.1101/2023.09.28.559981

**Authors:** Carlo Barnaba, David G. Broadbent, Gloria I. Perez, Jens C. Schmidt

## Abstract

Autophagy is an important metabolic pathway that can non-selectively recycle cellular material or lead to targeted degradation of protein aggregates or damaged organelles. Autophagosome formation starts with autophagy factors accumulating on lipid vesicles containing ATG9. These phagophores attach to donor membranes, expand via ATG2-mediated lipid transfer, capture cargo, and mature into autophagosomes, ultimately fusing with lysosomes for their degradation. Autophagy can be activated by nutrient stress, for example by a reduction in the cellular levels of amino acids. In contrast, how autophagy is regulated by low cellular ATP levels via the AMP-activated protein kinase (AMPK), an important therapeutic target, is less clear. Using live-cell imaging and an automated image analysis pipeline, we systematically dissect how nutrient starvation regulates autophagosome biogenesis. We demonstrate that glucose starvation downregulates autophagosome maturation by AMPK mediated inhibition of phagophores tethering to donor membranes. Our results clarify AMPK’s regulatory role in autophagy and highlight its potential as a therapeutic target to reduce autophagy.

## Introduction

Autophagy is a catabolic pathway activated under conditions of chemical and nutrient stress, allowing the recycling of cellular material to sustain metabolism. Depending on the target for degradation, autophagy can be categorized into non-selective and selective forms, with the latter targeting specific substrates, for instance damaged organelles, for breakdown ^1,2^. The cellular material targeted for degradation is enclosed within a *de novo* structure known as the autophagosome, which fuses with the lysosome to recycle its content ^3,4^. Autophagy serves two important functions; it is critical in degrading damaged organelles and protein aggregates, and it provides molecular building blocks under starvation and stress conditions. As a consequence, defects in autophagy are associated with neurodegenerative diseases and autophagy is activated in cancers to provide building blocks for their rapid proliferation ^1,5–8^.

The autophagosome is a vesicle enclosed by two lipid membranes, and its initiation and assembly are tightly controlled by autophagy genes (ATGs) ^1^. One model for autophagosome biogenesis proposes that ATG9 containing vesicles serve as the seed for recruitment of other autophagy factors ^9–11^. ATG9 is the only known transmembrane protein among the ATGs and has lipid scramblase activity, transferring phospho-lipids between the two lipid bilayer leaflets ^11,12^. Mobile ATG9 vesicles are specified to form autophagosomes by a phospho-lipid signaling cascade, enabling the recruitment of autophagy machinery required for autophagosome maturation ^9–11^. Key maturations factors include the scaffold protein WIPI2 and the ATG16L1-ATG12-ATG5 complex, which acts like a ubiquitin ligase conjugating phosphatidylethanolamine to ATG8-like proteins (LC3, GABARAP) ^1,13,14^. The expansion of the ATG9 vesicle seed into a fully formed autophagosome occurs upon its tethering to a lipid-membrane source by ATG2, a lipid transfer protein that transports lipids to the growing autophagosome structure ^12,15,16^.

Nutrient stress can induce autophagy via several mechanisms, depending on the type of starvation ^17,18^. Amino acids, especially leucine, glutamine, and arginine, are essential activators of the mammalian target of rapamycin complex 1 (mTORC1) ^19,20^. When these amino acids are absent or levels are low, mTORC1 dissociates from the lysosome which relieves its inhibition of the Unc-51-like autophagy activating kinase (ULK1/2) complex ^19^. Activation of the ULK1-kinase complex triggers the phospho-lipid signaling cascade by activating a PI3K-kinase complex, causing local accumulation of PI3P on ATG9 vesicles ^20,21^. Lack of growth factors also promotes autophagy by an mTORC1-dependent mechanism ^22^. In contrast, how the autophagy machinery responds to energy starvation, in particularly when triggered by glucose withdrawal, has been a subject of intense debate ^23–26^. In mammalian cells, the AMP-activated protein kinase (AMPK) complex senses the AMP/ATP ratio and activates catabolic pathways including fatty acids catabolism and potentially autophagy, concurrently suppressing biosynthetic pathways ^20^. AMPK directly phosphorylates ULK1, blocking its interaction with mTORC1 and thus activating its kinase activity towards downstream signaling phospholipid complexes ^27^, including the recently reported PIKfyve complex, which was suggested to upregulate autophagy ^23^. In contrast, other studies have shown that glucose starvation suppresses autophagic flux in both mammalian cells ^24^ and yeast ^25^. In addition, *Park et al.* recently proposed that AMPK activation results in ULK1 phosphorylation at Ser^556^ and Thr^660^, leading to ULK1 inactivation, an increase stability of the AMPK/ULK1 complex, and inhibition of autophagy ^26^. It is critical to resolve these conflicting results, given that AMPK has been extensively explored as a drug target to treat several chronic diseases ^28,29^. For instance, metformin and canagliflozin are FDA-approved drugs to treat type 2 diabetes, with ongoing preclinical trials exploring their potential as therapeutics for cardiovascular disease and several types of cancer ^30,31^. They indirectly target AMPK by inhibiting the mitochondrial respiratory chain and thus decreasing the cellular ATP levels ^31^. While direct AMPK activators like compound 991 and MK8722 exist, their use is currently primarily limited to in vitro or animal models ^31^.

To systematically analyze how amino acids, glucose, and growth factors regulate phagophore initiation and maturation into autophagosomes, we used our recently developed collection of cell lines expressing HaloTagged autophagy factors from their endogenous loci ^9^. This approach allows us to analyze autophagic flux by determining the initiation and maturation kinetics of autophagosomes in living cells, avoiding artifacts due to protein overexpression ^32,33^. In this study, we expand on our previous work by developing a high-throughput computational pipeline (K-FOCUS), which combines automated cell segmentation, with multi-color single-particle tracking to dissect the kinetics of autophagosome maturation at the single-cell level. K-FOCUS assesses protein co-localization by analyzing the co-diffusion of fluorescent signals over time, and outperforms traditional method of colocalization analysis, including object-based algorithms such as SODA ^34^, which are often limited to a single timepoint. Using this methodology, we investigate autophagosome biogenesis under various nutrient conditions, including glucose withdrawal, focusing on ATG13, ULK1, WIPI2, and ATG2A recruitment to phagophores/autophagosomes. Our results demonstrate that the rate of phagophore initiation is increased by amino acid or glucose starvation. However, the fraction of phagophores that mature to the point of LC3 accumulation is significantly reduced upon glucose starvation, suggesting that phagophore maturation is inhibited when glucose is absent. Importantly, direct activation of AMPK with a small molecule mimics this effect, and inhibition of AMPK increases the maturation of autophagosomes in the absence of glucose, suggesting that AMPK activation is sufficient to inhibit autophagosome maturation. Finally, we demonstrate that upon glucose starvation or AMPK activation, autophagy proteins accumulate on highly mobile structures, consistent with ATG9 vesicles that fail to be tethered to a donor membrane by ATG2. Collectively, these results support a model in which glucose starvation and AMPK activation inhibit autophagosome biogenesis by preventing the tethering of ATG9 vesicles to donor membranes.

## Results

### K-FOCUS: single-cell monitoring of autophagy dynamics

In fluorescence microscopy images of eukaryotic cells, autophagosomes appear as bright cytoplasmic puncta, also referred to as foci ^35^. Using time-lapse imaging, it is possible to monitor the rate of foci formation, their lifetime and mobility ^9^. These metrics provide valuable insights into the dynamics of autophagic flux and the molecular mechanisms underlying autophagosome biogenesis. To study autophagy in human cancer cells, we have developed a cell line panel in which the HaloTag is introduced at the endogenous loci of the autophagy factors ATG13, ULK1, WIPI2 and ATG2A ^9^. This panel of cell lines provides a sensitive tool to detect autophagosome formation. Notably, like many other biological processes ^36^, the number of autophagy factor foci formed displays substantial cell-to-cell variability ^37^. To dissect autophagosome formation at the single-cell level, we have created K-FOCUS, a computational pipeline to analyze autophagic flux at the single-cell level using live-cell imaging. K-FOCUS allows simultaneous analysis of multiple autophagy markers to examine autophagosome maturation, for example by analyzing an initiation factor and a downstream cargo-adaptor (P62) or ATG8-like protein (LC3, GABARAP).

K-FOCUS encompasses three key steps: single-cell segmentation, foci localization and tracking, and colocalization analysis (**Fig. 1**). For cell segmentation, we used CellPose ^38,39^, a deep-learning tool, which was trained using cells expressing GFP-LC3, and provides a reliable outline of the cells analyzed (**Fig. 1A-B**). This cell segmentation is then used for the analysis of all imaging channels. Cellular segmentation remained robust even under conditions of high cell confluency (**Fig. 1A**). The resulting single-cell regions of interest (ROIs) were imported into TrackIt, a GUI-based tool used for single-particle tracking ^40^. Among the available tracking software options, we selected TrackIt for its user-friendly GUI interface that enables high throughput optimization and analysis of multiple ROIs per image. Furthermore, TrackIt uses a wavelet analysis to detect single-particles ^41^, which outperforms other algorithms when analyzing the Brownian motions of vesicles, as well as particles switching from random to directed motion ^42^. Prior to tracking, manual inspection of the cellular segmentation was performed in TrackIt to correct any miss-segmented cells (**Fig. 1B**). Subsequently, tracking was carried out for all channels by optimizing the tracking parameters and exported as a batch file to be used as the input file for the colocalization analysis with K-FOCUS. Two parameters are particularly critical when tracking foci in TrackIt: the threshold factor and tracking radius. In TrackIt, foci were filtered using an intensity threshold factor to avoid the detection of false positive signal and optimized separately for each protein analyzed. After threshold filtering, the nearest neighbor algorithm will link spots that are nearest neighbor in two consecutive frames based on a maximum used-defined tracking radius. In our experiment, a tracking radius of 5 pixels (∼1 μm) was optimal for both autophagy factors and LC3. In both cases, visual inspection of tracking was carefully performed for each experiment.

**Figure 1.**
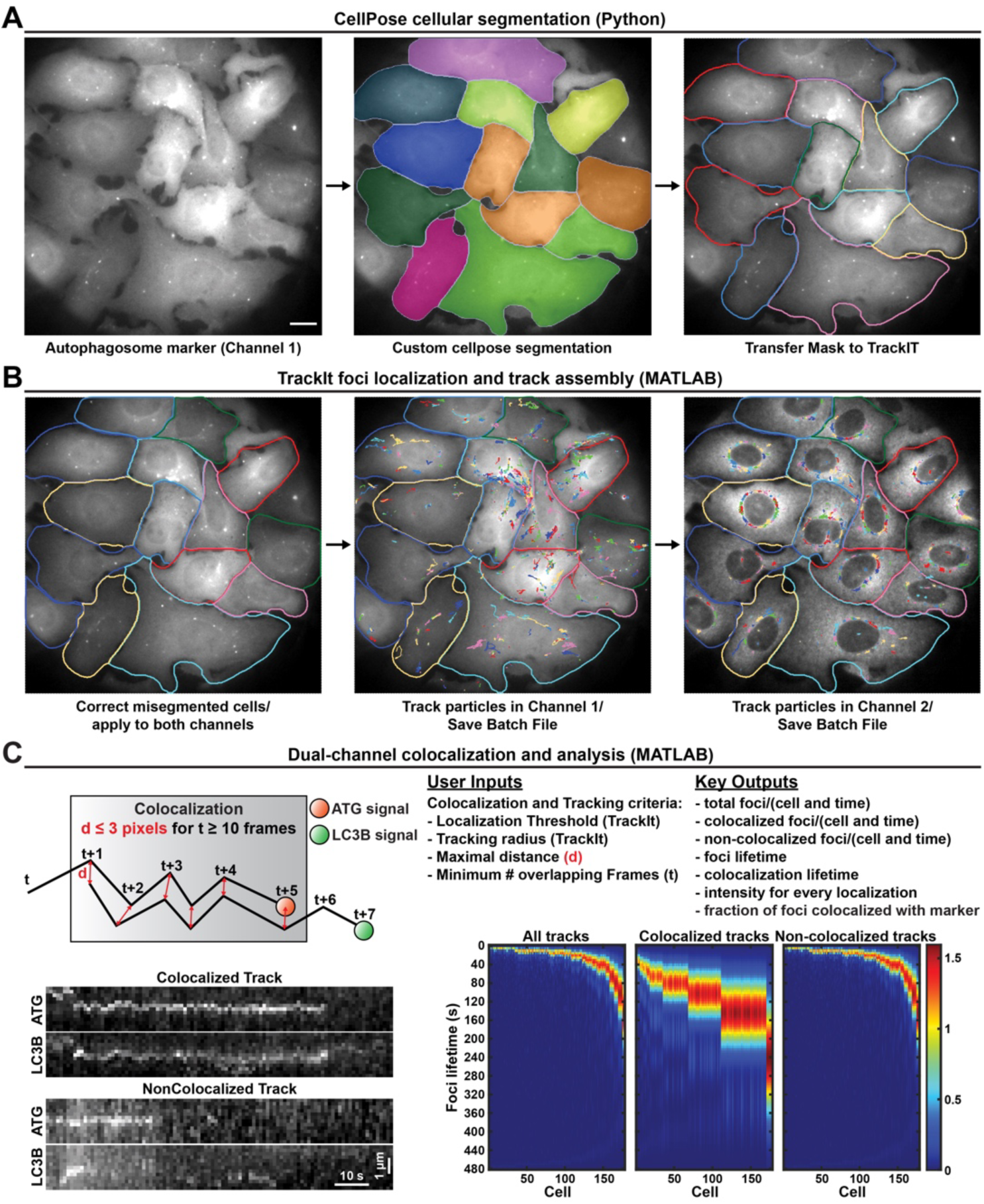
K-FOCUS: Live-cell high-throughput single-cell analysis of foci colocalization. (**A**) Illustration of the analysis pipeline for the Cellpose-TrackIt module, which incorporates CellPose segmentation into TrackIt (scale bar = 10 μm). **(B)** Workflow for manual ROI quality control and foci tracking using the TrackIt GUI. **(C)** A schematic outlining the colocalization criteria, including user-defined inputs, and key outputs. Additionally, it presents a visual representation of a colocalized and non-colocalized track, along with a kernel density plot of track length per cell for both colocalized and non-colocalized tracks.

K-FOCUS enables the processing of hundreds of cells, generating time-lapse data for thousands of autophagy foci (**Fig. 1C**). K-FOCUS determines the colocalization of foci in multiple fluorescence channels and calculates the frequency of foci occurrence, foci lifetime, and diffusion dynamics. Once particle tracks have been determined in the imaging channels, K-FOCUS carries out the colocalization analysis between channels. The precise colocalization criteria can be defined by the user. We typically consider two signals colocalized if their centroids are within 3 pixels (∼650 nm in our microscopy setup) of each other for at least 10 imaging frames (10 seconds), which is highly stringent. In the context of autophagosome biogenesis a critical output is fraction of autophagy factor foci that colocalizes with a distinct autophagosome marker to assess autophagosome maturation. K-FOCUS calculates the total number of foci for both channels, and the number and fraction of mature autophagosome foci (e.g., GFP-LC3) that colocalize with autophagy protein foci (e.g., Halo-ATG13). In addition, K-FOCUS determines foci lifetime, colocalization time, the time delay before colocalization occurs, step sizes of all tracks, background corrected fluorescence intensity for all foci localizations, among others. The K-FOCUS output is a MATLAB structure containing single-cell foci data that can be easily used for additional analyses. Overall, K-FOCUS provides detailed information defining the life cycle of autophagosomes at the single-cell level.

### K-FOCUS outperforms other approaches to determine foci colocalization

Several metrics have been traditionally used to assess the degree of colocalization between two fluorescent signals in fixed cells or single images of living cells. Threshold-based methods such as Pearson’s correlation coefficient and Manders’ overlap coefficient offer distinct advantages. These metrics are widely used and standardized, enabling comparisons across different experiments and facilitating a quantitative understanding of colocalization ^43^. Recently, the Statistical Object Distance Analysis (SODA) was developed as a statistical approach to map coupled objects within the cell, providing insights into molecular assemblies in high-resolution fluorescence imaging ^34^. To test how K-FOCUS compares with threshold and object-based methods, we calculated these metrics at single-cell level using two datasets that exemplify common fluorescence colocalization challenges in the autophagy field (**Fig. 2**). Cells expressing Halo-ATG9A from its endogenous locus, and LAMP1-mNeonGreen, a marker of lysosomes, were selected as the first test dataset. We recently demonstrated that Halo-ATG9A accumulates in lysosomal compartment over time ^9^.

**Figure 2.**
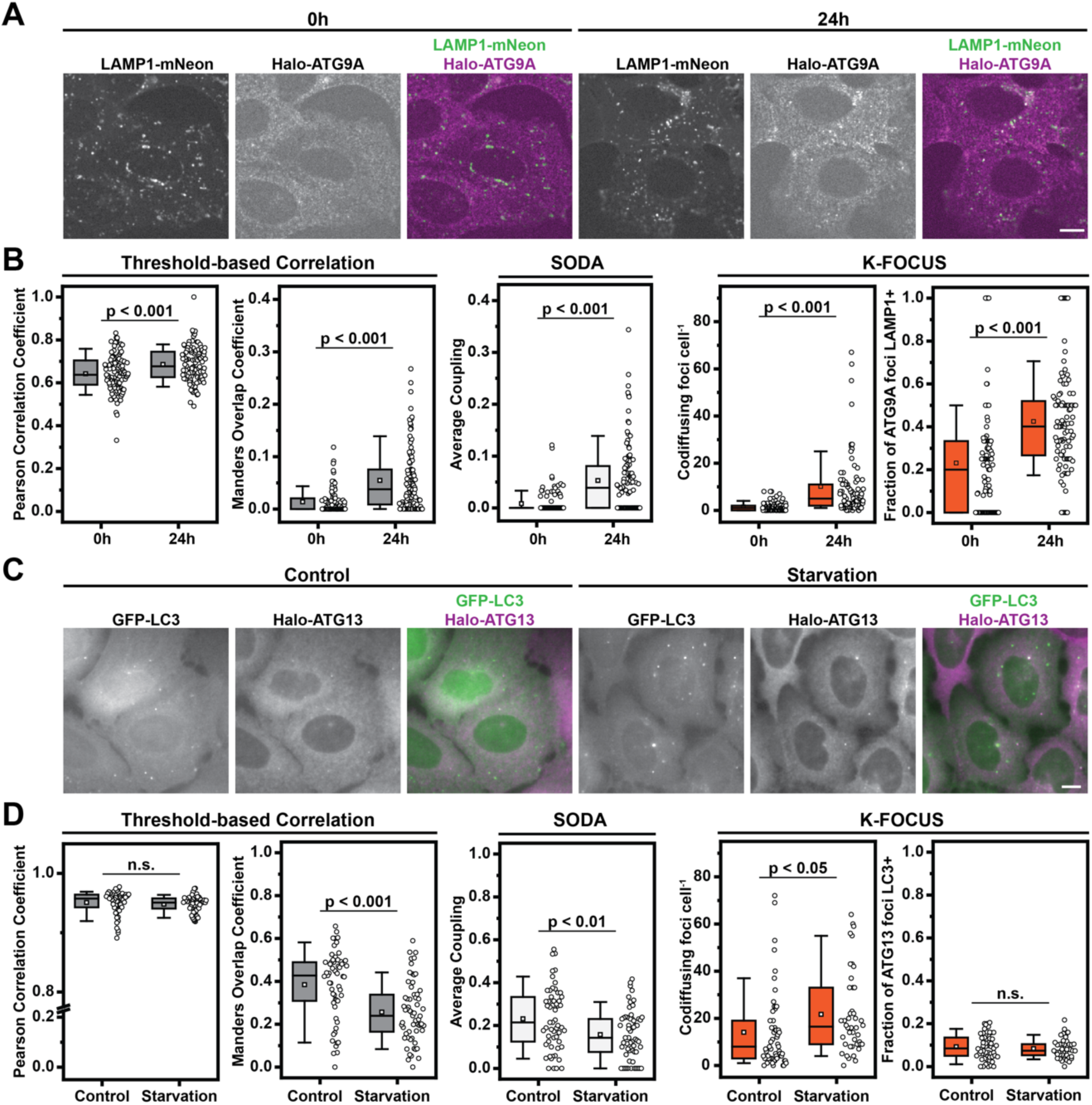
K-FOCUS is a robust object-based colocalization tool in live-cell imaging. (**A**) Example images demonstrating the accumulation of Halo-ATG9A in lysosomes, captured immediately after or 24 hours following labeling with JFX650 in U2OS cells (scale bar = 10 μm). **(B)** A comparison between different colocalization analysis methods for the data shown in (**A**), including Threshold-based Pearson and Manders coefficients, as well as wavelet spot detection-based SODA and K-FOCUS. The box indicates the interquartile range, the whiskers the 10-90% confidence interval, the square indicates the average, and the horizontal line is the median. **(C)** Example images of U2OS cells expressing GFP-LC3B and Halo-ATG13 under both control and EBSS conditions (scale bar = 5 μm). **(D)** A comparison between different colocalization analysis methods for the data shown in (**C**), including Threshold-based Pearson and Manders coefficients, as well as wavelet spot detection-based SODA and K-FOCUS analyses. The box indicates the interquartile range, the whiskers the 10-90% confidence interval, the square indicates the average, and the horizontal line is the median.

The use of HaloTag, which is resistant to lysosomal proteolysis ^44^, allows us to track the accumulation of Halo-ATG9A by pulse labeling with a HaloTag ligand conjugated with JFX650 and monitoring its co-localization with the lysosomal marker LAMP1-mNeonGreen over time (**Fig. 2A, Movie S1**). Immediately after labeling we expect to observe minimal colocalization and significant colocalization should be observed 24 hours after labeling. After CellPose cell segmentation, we analyzed a single frame per cell and performed colocalization analysis (**Fig. 2B**). As expected, both threshold-based methods and SODA showed a significant increase of Halo-ATG9A and Lamp1-mNeonGreen colocalization 24 hours after labeling. Rather than producing a coupling or colocalization coefficient, K-FOCUS provides a numerical value for total foci number per cell for each channel and a fraction of colocalized signals. Next, we generated a second dataset by imaging cells expressing Halo-ATG13 from its endogenous locus and GFP-LC3 by integration into the AAVS1 locus, which provides stable and homogeneous expression levels. Importantly, both fluorescence channels display substantial cytoplasmic background signals. Accumulation of GFP-LC3 serves as a marker for the transition from a phagophore to an autophagosome ^45^. In the context of autophagosome formation, Halo-ATG13 forms foci that detectably accumulate GFP-LC3 ∼40 seconds after their appearance ^9^. Autophagy occurs under nutrient-rich conditions (basal autophagy) and increases when autophagy is triggered by nutrient starvation (**Fig. 2C, Movie S2**). Despite our edited cell lines showing a strong response to starvation based on the LC3 biochemical assay ^9^, Pearson’s correlation coefficient did not detect an increased overlap between LC3 and ATG13 (**Fig. 2D**). Even more surprisingly, both Mander’s overlap coefficient and SODA analysis yielded less colocalization between the Halo-ATG13 and GFP-LC3 channels under starvation conditions compared to control cells (**Fig. 2D**). In contrast, K-FOCUS was able to detect a significant increase in the number of ATG13 foci that colocalized with LC3 signals under starvation conditions compared to control cells (**Fig. 2D**). Interestingly, the fraction of ATG13 foci that co-localized with LC3 did not change, suggesting that the total number of ATG13 foci has increased in starved cells (**Fig. 2D**), consistent with previous results ^9^.

In summary, these results demonstrate that K-FOCUS provides higher sensitivity and more detailed information than other fluorescence colocalization methods by providing numerical values for colocalization of foci overtime, offering a novel tool for the quantitative colocalization of fluorescence signals in live-cell microscopy data.

### Glucose depletion inhibits phagophore maturation

In response to low nutrient levels, autophagy is induced as a catabolic process to recycle cytosolic material, to provide the building blocks required to sustain cellular metabolism ^17^. Regulation of autophagy occurs by sensing the level of cytosolic nutrients. The mTOR complex inhibits autophagy in the presence of essential amino acids ^19,20^. In contrast, the AMP-activated protein kinase (AMPK) complex is a cellular energy sensor, activated by a high AMP-to-ATP ratio ^20^. To investigate how the removal of specific class of nutrients impacts the formation of autophagy factor foci, we performed live-cell imaging of human U2OS cancer cells expressing HaloTagged ATG13, ULK1, WIPI2 and ATG2A from their endogenous loci. The selection of these proteins allows us to characterize autophagosome biogenesis starting with protein kinase signaling (ATG13 and ULK1), to lipid transfer (ATG2A), and finally the LC3-conjugation machinery (WIPI2). In these HaloTagged cell lines, we stably expressed GFP-LC3 from the AAVS1 safe-harbor locus to mark mature autophagosomes ^9^.

We systematically analyzed autophagy foci formation in media lacking amino acids, glucose, amino acids and FBS, or all three components (**Fig. 3A, Fig. S1, Movie S3-S6**). Cells were imaged every second and movies were analyzed using K-FOCUS to determine three key metrics: foci formation rate (phagophore initiation rate), the fraction of autophagy foci that recruited LC3 (conversion ratio), and foci lifetimes. The foci formation rate was further stratified into autophagy factor foci that colocalized with GFP-LC3 signal over time (LC3+ foci) and those that do not (LC3-foci). We observed changes in both endogenous HaloTagged autophagy factor and GFP-LC3 foci formation dynamics when cells were subjected to different media conditions (**Fig. 3A, Fig. S1, Movie S3-S6).** Removal of amino acids was not sufficient to increase endogenous phagophore formation rate of the autophagy proteins (**Fig. 3B**, left panels), with the exception of Halo-ATG2A, which showed a slight increase from 6 to 11 foci per cell per minute. Removal of fetal bovine serum (FBS) in addition to amino acids increased the formation rate of Halo-ATG2 (from 6 to 15 foci per cell per minute) and Halo-ATG13 (19 to 38 foci per cell per minute) foci (**Fig. 3B**). The median foci formation rate per cell also increased for WIPI2 and ULK1 in the absence of amino acids and FBS but the difference was not statistically significant (**Fig. 3B**). When glucose was removed from the media, the foci formation rate was increased for Halo-ATG13 and Halo-ATG2A (**Fig. 3B**). In addition, we observed a significant increase in formation of WIPI2 foci when amino acids, FBS, and glucose were withheld (**Fig. 3B**). These observations suggest that nutrient starvation, in particular the removal of amino acids and FBS, or glucose can increase the autophagic foci formation rate for ATG13 and ATG2, but only has only marginal effects on the number of foci formed by ULK1 and WIPI2. It is important to note that ATG13 forms significantly more foci per cell compared to all other factors analyzed, consistent with previous results ^9^. This could be a consequence of ATG13 being one of the first factors recruited to phagophores, or that ATG13 also accumulates on structures not involved in autophagy.

**Figure 3.**
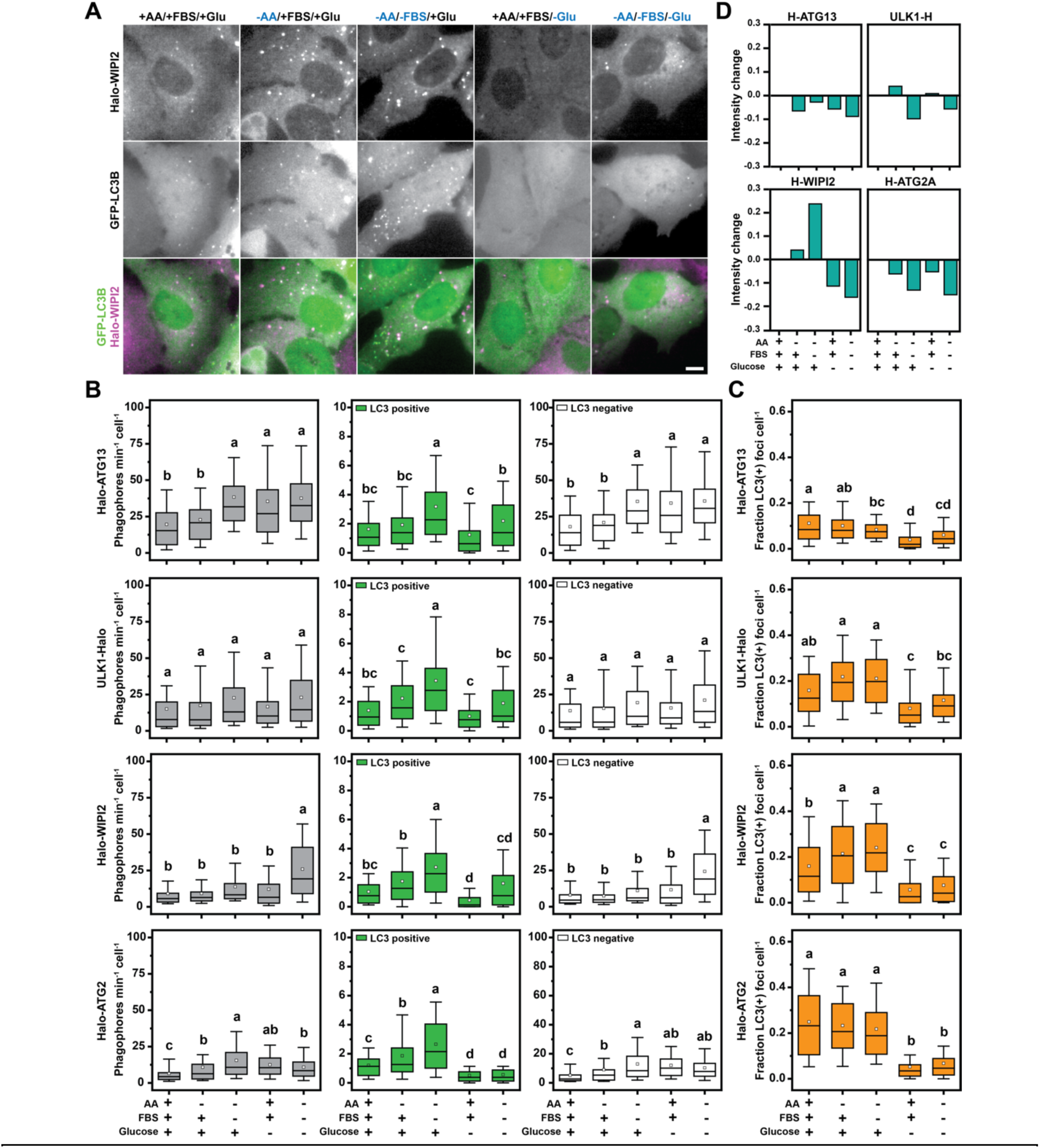
K-FOCUS analysis of the kinetics of autophagy factor foci formation under nutrient starvation. (**A**) Live-Cell images of Halo-WIPI2 and GFP-LC3B in control and starvation medium (scale bar = 5 μm). **(B)** Quantification of colocalization kinetics of single cells in (A) and (Fig. S1) using K-FOCUS including total, colocalized, and non-colocalized phagophore formation rates. **(C)** Quantified fractions of LC3+ foci/cell from imaging in (A) and Fig.S1. For (B) and (C) the box indicates the interquartile range, the whiskers the 10-90% confidence interval, the square indicates the average, and the horizontal line is the median. **(D)** Intensity quantification of autophagy proteins from (A) and Fig. S1, including Halo-ATG13, Halo-ATG2A, Halo-WIPI2, and ULK1-Halo.

The number of autophagy factor foci that accumulate LC3 (LC3+ foci) were significantly increased in the absence of amino acids or amino acids and FBS (**Fig. 3B**), consistent with the established role of mTOR signaling in autophagy induction ^19,46^. All proteins had comparable rates of LC3+ foci formation at maximal induction (∼3 LC3+ phagophores per cell per minute in –AA/-FBS/+Glucose media), confirming that autophagic flux is similar in all edited cell lines. The absence of glucose significantly decreased the rate at which LC3+ foci formed for all autophagy factors analyzed, suggesting that autophagosome maturation is inhibited in the absence of glucose (**Fig. 3B**). It is important to note that only a small fraction (∼10-20%) of the autophagy factor foci colocalizes with LC3 (**Fig. 3B**). For this reason, the formation rate of autophagy protein foci that do not colocalize with LC3 closely corresponds to the overall foci formation rate (**Fig. 3B**).

The conversion ratio, which represents the fraction of HaloTagged autophagy foci that detectably accumulate GFP-LC3 signal over the imaging time course was not affected by withdrawal of amino acids or amino acids and FBS for ATG13 and ATG2 but was increased for ULK1 and WIPI2 under the same conditions (**Fig. 3C**). This suggest that the increase in LC3 positive ATG13 and ATG2 foci is largely driven by an increase in autophagosome formation, while ULK1 and WIPI2 foci more efficiently mature into LC3 positive autophagosomes in the absence of amino acids or amino acids and FBS. This is consistent with ULK1 and WIPI2 controlling key autophagosome maturation steps, which are upregulated under amino acid starvation ^45,47^. In contrast, glucose withdrawal alone, or in the context of media also lacking amino acids and FBS significantly reduced the fraction of autophagy protein foci that accumulate LC3 for all proteins tested. This demonstrates that autophagosome maturation is strongly inhibited in the absence of glucose.

Next, we examined the lifetime of autophagy protein foci to determine how the different nutrients affect the kinetics of autophagosome biogenesis (**Fig. S2, Fig. S3A**). We first determined the lifetime of LC3+ and LC3-autophagy factor foci. LC3-foci had a lifespan of approximately 35 seconds for all proteins analyzed. In contrast, LC3+ foci autophagy factor foci lasted significantly longer, with lifetimes ranging from 75 to 90 seconds depending on the specific autophagy protein, consistent with our previous results ^9^. We only observed minimal changes in these lifetimes under the different nutrient conditions tested (**Fig. S2, Fig. S3A**).

In our imaging experiments, we can analyze four distinct steps of autophagosome biogenesis: the accumulation of autophagy proteins (ATG signal only), the appearance of LC3 (ATG and LC3 signal colocalization), the dissociation of autophagy factors (LC3 signal only), and the disappearance of LC3 signal due to fusion of the autophagosome with the lysosome ^9^. To define how these critical steps in the autophagosome live cycle are regulated by nutrient availability, we determined their duration under the different media conditions (**Fig. S3B**). The time from autophagy protein foci formation to GFP-LC3 signal appearance was ∼25 s for Halo-ATG13, ULK1-Halo, and Halo-ATG2A, while Halo-WIPI2 exhibited a longer delay of ∼50 s (**Fig. S3B**). Similar to overall autophagy protein foci lifetime, we observed only marginal changes in the pre-colocalization and LC3-colocalization lifetimes under different nutrient conditions (**Fig. S3B**).

To determine whether nutrient starvation impacts quantity of autophagy factors recruited to phagophores, we analyzed the intensity of autophagy foci in the different media conditions (**Fig. 3D**). For Halo-WIPI2, autophagy induction by withdrawal of amino acids and FBS led to a significant increase (∼20%) in foci intensity, suggesting that WIPI2 recruitment is increased under these conditions or the size of the autophagosome is increased (**Fig. 3D**). In contrast, removal of glucose resulted in a ∼10% decrease in WIPI2 foci intensity, compared to control conditions, suggesting that phagophores are smaller or less WIPI2 is recruited (**Fig. 3D**). For all other proteins tested, all starvation conditions led to slight decreases in foci intensity, suggesting that their recruitment to phagophores is largely unchanged (**Fig. 3D**).

In total these observations confirm that the rate of autophagosome formation is increased by amino acid and FBS withdrawal. In addition, our results demonstrate that glucose withdrawal significantly inhibits the maturation of autophagy factor foci into LC3 containing autophagosomes without dramatically changing the rate of phagophore initiation, leading to an overall downregulating of autophagosome biogenesis.

### Activation of AMPK downregulates autophagy

In cultured eukaryotic cells, glucose availability is directly linked to ATP levels primarily by its metabolism via glycolysis and the citric acid cycle ^48^. ATP levels are sensed by the AMP-activated protein kinase (AMPK), a protein kinase complex that is activated by a high AMP to ATP ratio ^49^. The regulation of autophagy by AMPK signaling is unclear with several studies suggesting AMPK activation promotes autophagy ^23^, while others suggest it has an inhibitory effect on autophagosome formation ^24,26^. To determine whether the reduction in autophagosome maturation, observed in the absence of glucose, is mediated by AMPK signaling, we used two small molecules to modulate AMPK activity. The AMPK agonist MK8722, a pan-AMPK allosteric activator, was used to stimulate AMPK in the presence of glucose ^50^. Conversely, the potent and selective AMPK inhibitor BAY3827, was used to reduce AMPK activity in the absence of glucose ^51^. Cells were treated with either the AMPK agonist or antagonist for 18 hours, followed by nutrient starvation (amino acid and FBS withdrawal, with glucose for the AMPK agonist, without glucose for the AMPK antagonist) and either continued treatment with the drug or its removal (**Fig. S4**). Importantly, we used a concentration of 1 µM MK8722 and BAY3827, which is substantially lower than the 10-50 µM used in other studies ^26,52–54^.

In cells cultured in glucose-containing media, the presence of the AMPK agonist led to a significant decrease in formation rate of LC3 positive autophagy factor foci in all four cell lines (**Fig. 4A, Fig. S4, Movie S7-S8**). Consequently, the fraction of autophagy factor foci that matured into LC3+ containing autophagosomes was reduced by AMPK stimulation in all HaloTagged cell lines, with the exception of ULK1-Halo (**Fig. 4A**). When AMPK was inhibited in cells cultured in the absence of glucose, we observed a significant increase in formation rate of LC3 positive autophagy factor foci across all cell lines, without affecting the formation of aborted autophagosomes (**Fig. 4B, Fig. S4, Movie S9-S10**). As a result, the fraction of LC3 positive foci was increased when AMPK was inhibited (**Fig. 4B**).

**Figure 4.**
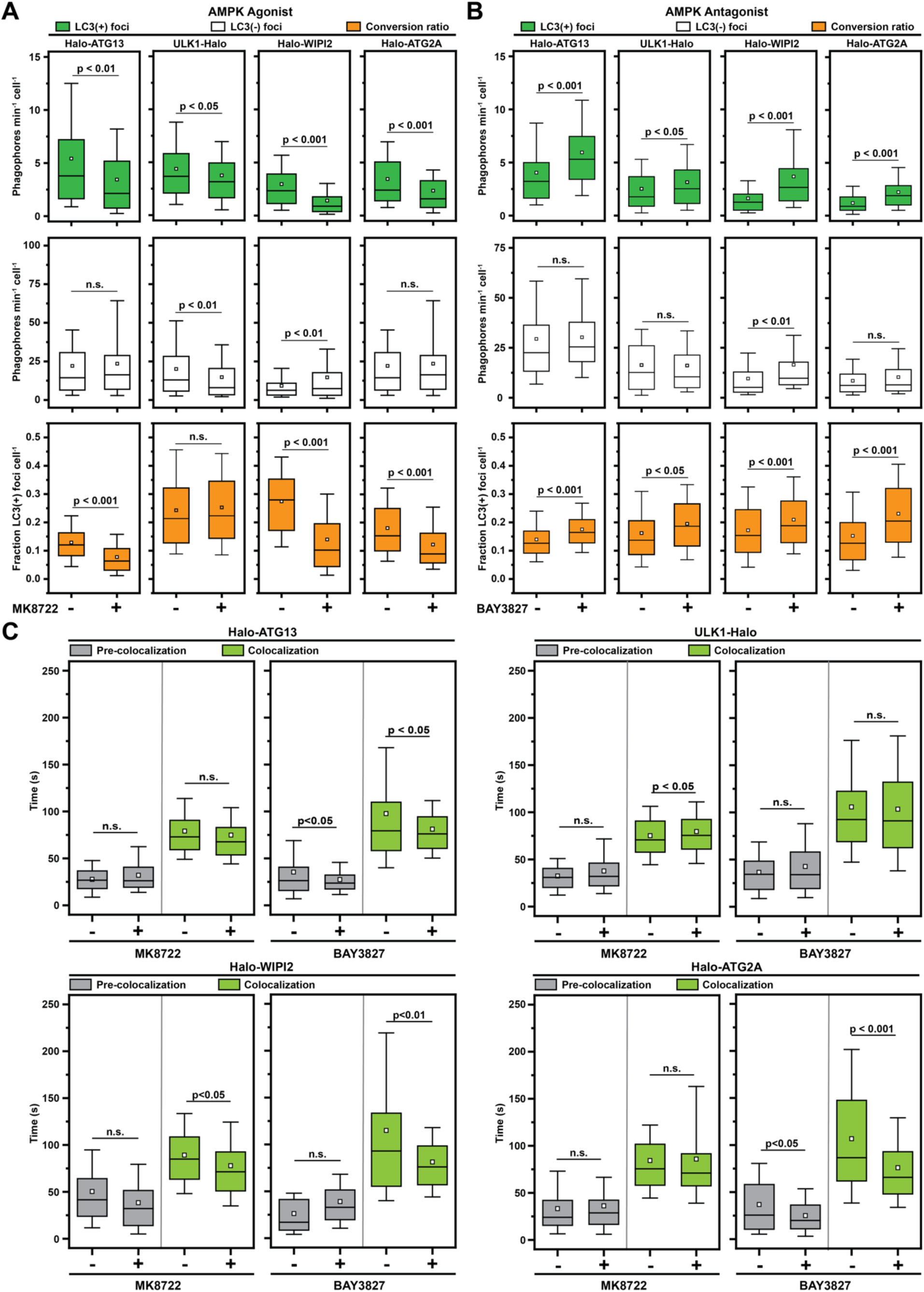
AMPK activation inhibits autophagosome maturation. (**A**) Formation rate and conversion ratio of LC3 positive and LC3 negative foci formed by Halo-ATG13, ULK1-Halo, Halo-WIPI2, and Halo-ATG2A in glucose containing media in the absence and presence of the AMPK agonist MK8722. **(B)** Formation rate and conversion ratio of LC3 positive and LC3 negative foci formed by Halo-ATG13, ULK1-Halo, Halo-WIPI2, and Halo-ATG2A in media containing no glucose in the absence and presence of the AMPK inhibitor BAY3827. (**C**) Lifetimes of foci formed by by Halo-ATG13, ULK1-Halo, Halo-WIPI2, and Halo-ATG2A before and during co-localization with GFP-LC3B in glucose containing media in the presence and absence of the AMPK agonist MK8722 (left panels) or in media lacking glucose in in the absence and presence of the AMPK inhibitor BAY3827 (right panels). For all plots the box indicates the interquartile range, the whiskers the 10-90% confidence interval, the square indicates the average, and the horizontal line is the median.

We further investigated whether direct modulation of AMPK activity would alter the biogenesis of autophagy factor foci, by analyzing their lifetimes. Similar to the findings under nutrient starvation (**Fig. S3**), we observed only minimal changes in the lifetime of autophagosome foci when AMPK was activated or inhibited with a small molecule drug (**Fig. 4C**). With the exception of a reduction in the duration of LC3-colocalization with Halo-ATG2A and Halo-WIPI2 (from ∼100s to ∼75s) when AMPK was inhibited with BAY3827 (**Fig. 3D**). This finding suggests that inhibition of AMPK signaling may increase the rate of phagophore expansion by ATG2 or LC3 conjugation mediated by WIPI2, shortening the time of autophagosome maturation.

Taken together, these experiments support the hypothesis that the decrease of autophagosome maturation observed under glucose starvation is directly mediated by the AMPK signaling, and further suggest that inhibition of AMPK activity can increase the efficiency of autophagosome formation. Furthermore, our results suggest that ATG2A activity and therefore tethering of the phagophore to a lipid donor membrane could be a critical target of AMPK signaling.

### ULK1 is required for ATG13 recruitment to phagophores after glucose starvation

While the formation of LC3 positive autophagosomes is inhibited by glucose withdrawal or AMPK activation, the formation of ATG13 foci is stimulated to a similar degree by amino acid and FBS starvation as glucose removal (**Fig. 3B**). Importantly, the vast majority (>90%) of ATG13 foci formed under glucose starvation are short lived (∼30 seconds) (**Fig. 3C, Fig. S3**). To define the role of the ULK1 complex in ATG13 foci formation in the absence of glucose, we imaged previously established Halo-ATG13 cell lines in which either ULK1, FIP200, or ATG101 were knocked out (**Movie S11-S13**) ^9^. The number of LC3 positive ATG13 foci was strongly reduced in ULK1, FIP200, and ATG101 knock-out cells compared to wildtype controls after removal of amino acids and FBS or glucose (**Fig. 5A**), consistent with the reduction in LC3 conjugation we previously observed in these cell lines ^9^. In contrast, the formation of short lived LC3 negative Halo-ATG13 foci was increased in the absence of amino acids and FBS when ULK1, FIP200, or ATG101 were knocked-out (**Fig. 5A**). In our prior study we observed a strong reduction in Halo-ATG13 foci in ULK1, FIP200, or ATG101 knock-out cells. However, in these experiments cells were only imaged every 15 seconds, which did not allow us to detect these short lived Halo-ATG13 foci ^9^. Under glucose starvation, the number of Halo-ATG13 was elevated in control cells, FIP200, and ATG101 knock-out cells (**Fig. 5A**). However, glucose removal did not lead to an increase in Halo-ATG13 foci in ULK1 knock-out cells (**Fig. 5A**). The lifetimes of LC3B positive and LC3B negative Halo-ATG13 foci were largely unaffected by the knockout of ULK1, FIP200, or ATG101 in all media conditions (**Fig. 5B**). Together, these observations suggest that ATG13 recruitment to the phagophore must be one of the first events in autophagosome biogenesis and its response to glucose starvation requires ULK1 but not ATG101 or FIP200.

**Figure 5.**
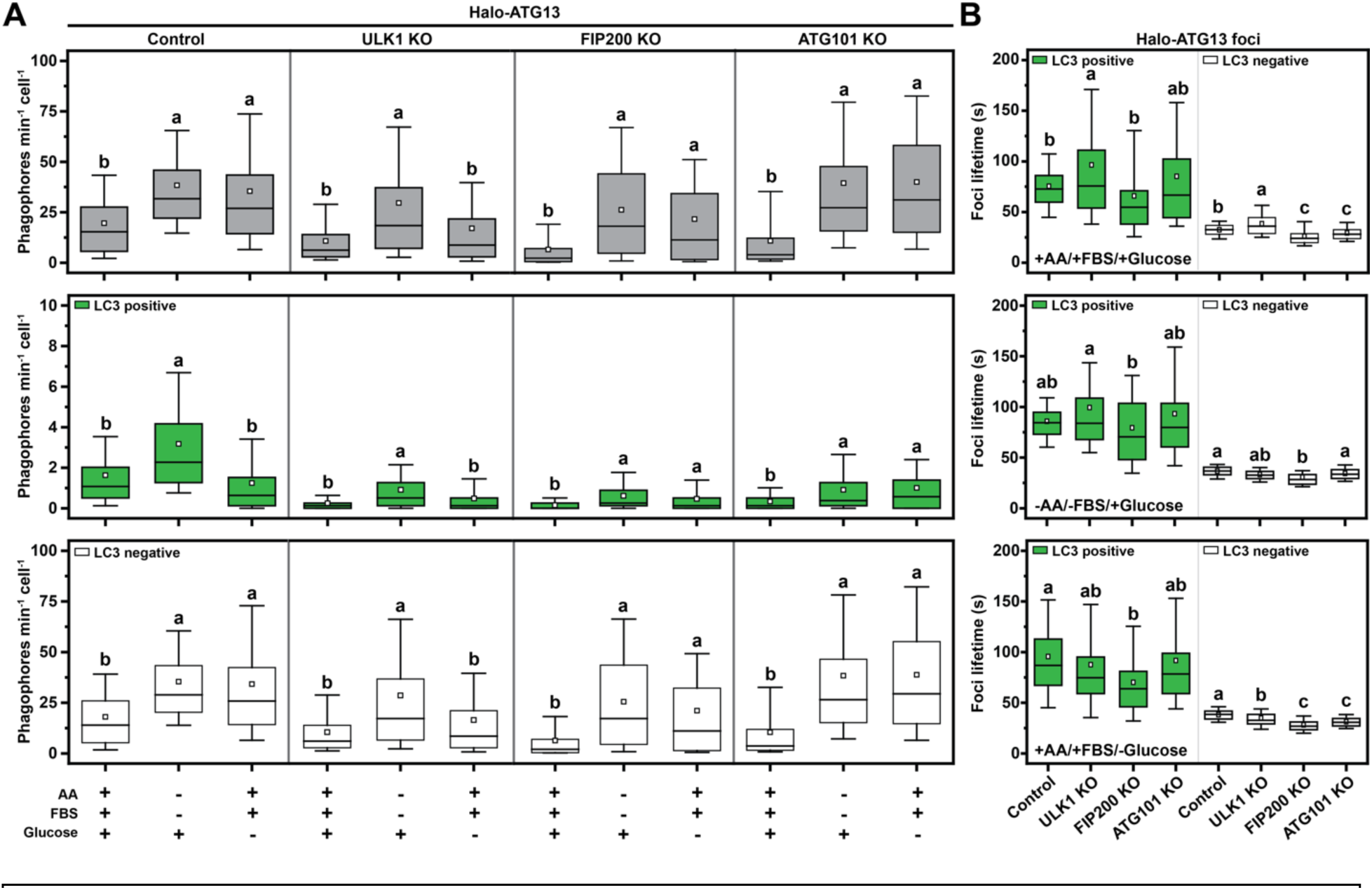
Contribution of the ULK1 complex to starvation induced autophagy protein foci formation. (**A**) Formation rate of total, LC3B positive, and LC3B negative Halo-ATG13 foci in control, ULK1, FIP200, and ATG101 knock out cells in various media conditions. **(B)** Lifetimes of LCB3 positive and LC3B negative Halo-ATG13 foci in control, ULK1, FIP200, and ATG101 knock out cells in various media conditions.

### AMPK modulates the membrane tethering of WIPI2 positive phagophores

An emerging model for autophagosome formation in human cells is the “vesicle seeding” model ^3,9–11,55^. In this model, an ATG9 vesicle is tethered to the phagophore initiation site by ATG2A. We have previously demonstrated that many autophagy factors, except for ATG2A, accumulate on mobile structures consistent with ATG9 vesicles ^9^. Using single-particle tracking, we have demonstrated that ATG9A vesicles exist in an untethered and tethered state with distinct diffusion properties (i.e. movement distances from frame to frame, **Fig. 6A**) ^9^. Our observations described in this study suggest that AMPK signaling could modulate this critical tethering step (**Fig. 4C)**. To investigate the role of AMPK signaling in vesicle tethering, we examined the mobility of autophagy factor foci under control and starvation conditions (–AA/-FBS/+Glucose and +AA/+FBS/-Glucose). Analysis of the step size distribution of autophagy factor trajectories allows us to assess the mobility of the underlying cellular structure (**Fig. 6A**). The step size distribution aborted phagophores (i.e. autophagy foci that never colocalize with LC3) and committed autophagosomes (i.e. autophagy foci that colocalize with LC3) are distinct. The step size distribution of LC3 positive autophagy foci is shifted towards smaller step sizes, which reflects their association with a donor membrane (**Fig. 6B-D, Fig. S5A**). This shift towards smaller step sizes upon LC3 accumulation was most dramatic for ULK1 foci and was also observed for WIPI2 and ATG2 foci (**Fig. 6B-D, Fig. S5A**). Surprisingly, the step size distribution of LC3 positive and negative ATG13 foci were comparable (**Fig. 6B-D, Fig. S5A**), suggesting that LC3 can accumulate on mobile ATG13 positive structures.

**Figure 6.**
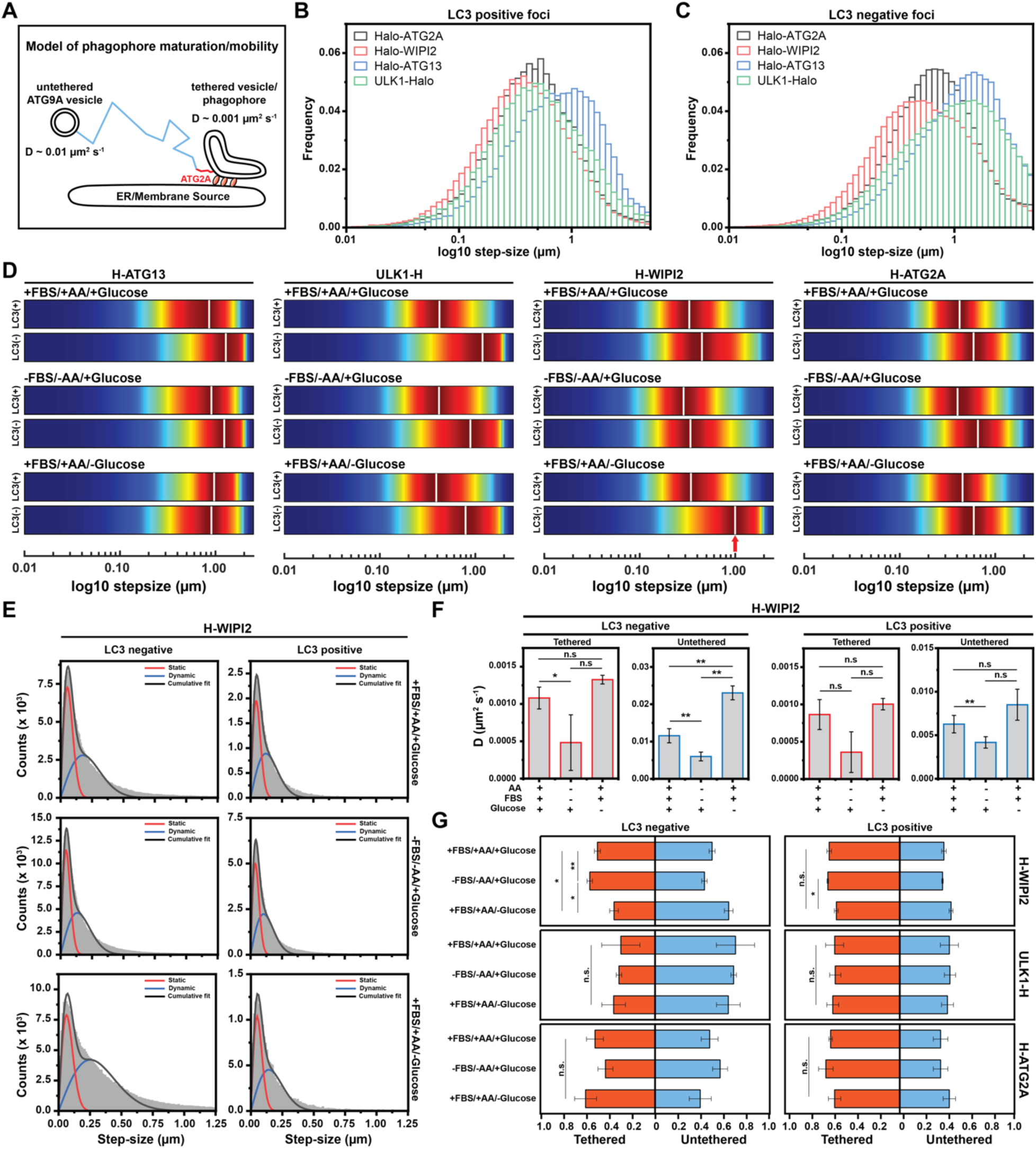
AMPK regulates tethering of WIPI2 positive phagophores. (**A**) Model depicting the diffusion properties of tethered phagophores and untethered ATG9 vesicles. (**B**-**C**) Step size distributions of Halo-ATG13, ULK1-Halo, Halo-WIPI2, and Halo-ATG2A trajectories for (**B**) LC3B positive and (**C**) LC3B negative autophagy factor foci. (**D**) Kernel density plots of the step size distributions of Halo-ATG13, ULK1-Halo, Halo-WIPI2, and Halo-ATG2A trajectories for LC3B positive and negative tracks in the indicated media conditions. (**E**) Step size distributions of LC3B positive and negative Halo-WIPI2 trajectories fit with a two-state diffusion model (black line) encompassing tethered (red) and untethered (blue) populations in the different media conditions. (**F**) Diffusion coefficients of the tethered and untethered populations of LC3B positive and negative Halo-WIPI2 trajectories in the different media conditions, derived from the fits shown in Fig. 6E. (**G**) Distribution of LC3B positive and negative Halo-WIPI2 trajectories between tethered and untethered populations, derived from the fits shown in Fig. 6E.

Using these populations as a baseline, we determined the impact of nutrient starvation on the movement pattern of foci formed by each autophagy protein. To better visualize changes in step size distributions, histograms were converted to 2D kernel density plots (**Fig. 6D, Fig. S5A**). Under control conditions LC3 negative foci of ATG13 and ULK1 moved faster than LC3 negative foci formed by ULK1 and WIPI2 (**Fig. 6D, Fig. S5A**), which likely reflects their early recruitment to mobile ATG9 vesicles. The step size distributions of ATG13 and ULK1 foci diverge when LC3 has been recruited. LC3 positive ULK1 foci move with smaller step sizes, while LC3 positive ATG13 foci exhibit both a slow– and fast-moving population (**Fig. 6D, Fig. S5A**), as previously observed ^9^. ATG13 was the only protein analyzed that displayed a mobile population that colocalized with LC3, and it is unclear whether these structures reflect and autophagosome biogenesis intermediate or a distinct organelle not involved in non-selective autophagy. WIPI2 and ATG2A foci, on the other hand, contain slower populations in the presence and absence of LC3, which likely signifies that their recruitment closely coincides with ATG9 vesicle tethering, which might precede detectable LC3 conjugation. Amino acid and FBS (–AA/– FBS/+Glucose) or glucose (+AA/+FBS/-Glucose) withdrawal had little impact on the step size distributions of LC3 positive and LC3 negative ATG13, ULK1, and ATG2 foci (**Fig. 6D, Fig. S5A**). While amino acid and FBS withdrawal did not change the mobility of WIPI2 foci, removal of glucose from the media led to a dramatic increase in the mobility of LC3 negative WIPI2 foci (**Fig. 6D, Fig. S5A**). AMPK activation in the presence of glucose mimicked this effect and AMPK inhibition in the absence of glucose reduced WIPI2 foci mobility (**Fig. S5B,C**). Importantly, the step size distribution of these mobile WIPI2 positive foci was comparable to ATG13 foci under the same conditions (**Fig. 6D, Fig. S5A**), consistent accumulation of these factors on mobile ATG9 vesicles. These observations suggest that phagophores that are rapidly tethered upon WIPI2 recruitment fail to be immobilized upon glucose starvation or AMPK activation.

To quantify the tethering defect of WIPI2 foci in the absence of glucose, the step-size distributions of foci trajectories were fit to Gaussian probability density functions to determine the diffusion coefficients and relative abundance of the tethered and untether foci populations ^56^. This two-state model (tethered and untethered) fit the step-size distribution with *R*^2^ > 0.98 in all cases and the calculated diffusion coefficients closely aligned with those recently reported (**Fig. 6A,E,F**) ^9^. Upon triggering autophagy by removing amino acids and FBS, we observed a substantial reduction (∼50%) in the diffusion coefficients of tethered and untethered WIPI2 foci (Fig. 5F). In contrast, glucose removal resulted in a significant increase in the mobility of untethered WIPI2 foci lacking LC3B signal (from D = 0.011 µm^2^ s^-1^ to D = 0.023 µm^2^ s^-1^). Furthermore, glucose depletion led to a noticeable rise in the fraction of untethered WIPI2 foci, increasing from 40% to approximately 60% (**Fig. 6G**). The distribution between untethered and tethered states of the other autophagy factors remained largely unchanged by amino acid and FBS or glucose withdrawal (**Fig. S6**).

In summary, these results suggest that AMPK exerts a negative regulatory influence on autophagy by directly impeding the membrane tethering process and WIPI2 accumulates on mobile ATG9 vesicles. Importantly, even though WIPI2 is recruited it appears that LC3B conjugation does not detectably take place under these circumstances.

## Discussion

In this study, we demonstrate that glucose starvation causes inhibition of autophagosome maturation, but not its initiation, in a human cancer cell model. Pivotal for this discovery is the implementation of K-FOCUS, a computational pipeline to quantitatively analyze multicolor live-cell imaging data. Our research offers mechanistic insights into the role of AMPK in regulating autophagosome maturation. These findings not only enhance our understanding of cellular responses to nutrient stress but also have implications for drug discovery efforts targeting AMPK, as modulating autophagosome maturation could be a promising avenue for therapeutic interventions in various diseases.

### Glucose starvation inhibits autophagosome maturation via AMPK

The role of AMPK in autophagy has been the subject of extensive research over the last few decades. Following the discovery that ULK1 is a direct substrate of AMPK ^27,57,58^, a consensus has emerged, establishing AMPK as a metabolic checkpoint and direct autophagy modulator, alongside mTORC1 ^59^. According to these early studies, under nutrient-rich conditions, mTORC1 phosphorylates ULK1, disrupting its interaction with AMPK and thereby leading to the inactivation of ULK1. Conversely, when levels of ATP are low AMPK activates ULK1 via direct phosphorylation on sites Ser^317^ and Ser^777^ ^27,57^. A few years after the discovery of ULK1 as direct AMPK substrate, reports indicated that glucose starvation has a protective effect on both human ^24^ and yeast cells ^25^. Particularly, Ramirez-Peinado *et al.* showed that removal of glucose from the media did not induce autophagy flux in four different cell lines, measured by determining the number of LC3 foci formed ^24^. Moreover, depriving the cells of glucose during autophagy induction did not confer protection against necrosis and apoptosis ^24^. Similarly, Nwadike et al. reported the suppression of autophagy under glucose starvation in mouse embryonic fibroblasts, both at the early and late stages of autophagy ^60^. These findings were further corroborated in a comprehensive study by Park *et al.* ^26^ on the AMPK-driven changes in ULK1 phosphorylation leading to inhibition of autophagosome formation.

To reconcile the discrepancies concerning the role of AMPK in autophagy, especially at early stages of phagophore initiation, we approached the problem from various experimental angles. First, we focused on improving the design of culture media conditions. To ensure that our results were not influenced by artifacts in commercial media, we utilized a consistent base for our media and systematically added nutrients. This approach aimed to clarify certain nuances that can affect experimental outcomes, such as the removal of not only glucose but also L-glutamine in previous work ^23^. In addition, we developed a single-cell imaging analysis pipeline – K-FOCUS – to measure the kinetics of autophagosome formation in living cells. K-FOCUS allowed us to assess the impact of nutrient starvation on early autophagy stages, including phagophore initiation and the recruitment of LC3 conjugation machinery by monitoring the onset of LC3 on autophagy factor foci ATG13, ULK1, WIPI2 and ATG2. Using K-FOCUS we were able to distinguish between long-lived autophagosome that show accumulation of LC3 (LC3-positive autophagosome) and short-lived LC3-negative autophagosome, or aborted autophagosomes, which represent the majority of autophagy protein foci formed in cells ^9^.

Our results demonstrate that glucose starvation causes a decrease in autophagy, by impeding the maturation of autophagosome. We observed this effect on all four of our genome-edited cell lines expressing different HaloTagged autophagy factors. Notably, the maturation rate of the small number of autophagosome that form in the absence of glucose is not affected under these conditions. While the formation of mature autophagosomes is reduced under glucose starvation, phagophore initiation, detected as short-lived autophagy factor foci, does not appear to be reduced, and might be increased for ATG13. Furthermore, the activation of AMPK using the agonist MK8722 mirrors the outcomes observed under glucose depletion, suggesting that AMPK is the driving force behind the observed reduction in autophagosome maturation when ATP availability is low. Our results contradict recent findings by Karabiyik et al. ^23^ who observed an increase of ATG16L1 puncta under glucose removal or direct activation of AMPK. It is worth noting that the authors used media that in addition to glucose also lacked L-glutamine, a major carbon source for cell growth, which when absent is a known autophagy inducer ^61^. To further substantiate the hypothesis that AMPK inhibits autophagosome maturation, we inhibited AMPK using BAY3827, resulting in enhanced autophagosome formation in response to nutrient stress. Interestingly, the absence of glucose had no impact on the lifespan of failed autophagosomes, and this trend remained consistent across all tested media conditions. Likewise, the lifespan of autophagosomes that exhibited LC3 accumulation remained unchanged, suggesting that AMPK’s effects occur prior to the recruitment of the LC3 conjugation machinery to the ATG9 vesicles.

### ATG13 recruitment to phagophores is independent of ATG101 and FIP200

In the current model for autophagosome biogenesis, ATG9 vesicles recruit phagophore initiation factors to their surface, including the ULK1 complex comprised of ATG13, ULK1, FIP200 and ATG101 ^21,62,63^. This complex initiate PI3K-mediated phospholipid signaling, leading to the accumulation of PI3P and other signaling phospholipids on the membrane of ATG9 vesicles ^1,9,11^. Our data show that the removal of amino acids and/or glucose resulted in an increase in the number of ATG13 foci, suggesting that phagophore initiation is increased (**Fig. 7**). It has been shown that colocalization of the ULK1 complex with the phagophore is required by both ATG13 and FIP200 ^21,64^, and PI3P signaling also contributes by promoting its recruitment via ATG13 membrane binding ^65,66^. Our findings indicate that ATG13 recruitment to ATG9 vesicles does not rely on ATG101, FIP200, or ULK1, since the number of ATG13 foci observed after autophagy induction is increased even when these three proteins are knocked out. Notably, both amino acid and glucose deprivation resulted in an increased ATG13-mediated phagophore formation in the knockout cell lines when compared to the parental cell line. However, the glucose starvation induced increase in the number of ATG13 foci was not observed in the ULK1 knockout. This suggests that ULK1 is required for the recruitment of ATG13 in response to AMPK activation.

**Figure 7.**
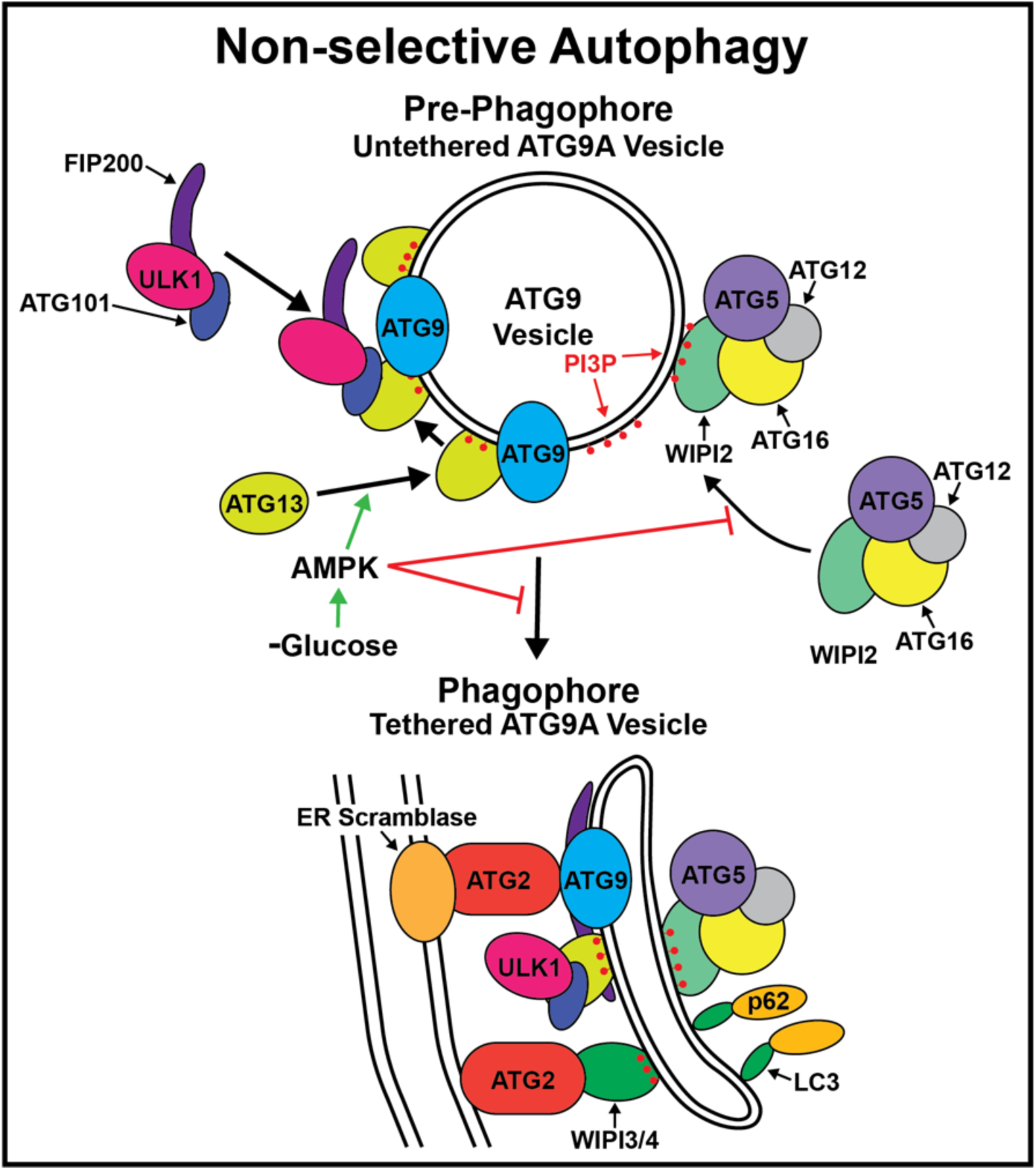
Model of the regulation of autophagosome biogenesis by AMPK. Glucose withdrawal leads to AMPK activation, which promotes the recruitment of ATG13 to mobile ATG9 vesicles. Under these conditions WIPI2 also accumulates on ATG9 vesicles but their tethering to donor membranes to allow phagophore maturation is inhibited by AMPK activation.

Importantly, in all knockout cell lines, we observed almost no mature LC3+ autophagosomes (<1 phagophore cell^-1^ min^-1^). The absence of long-lived autophagosomes is consistent with our previous report ^9^, and observations made by others ^62^, where loss of function of any of the ULK1 complex components impaired autophagy. It is important to note the absolute number of ATG13 foci formed after autophagy induction (∼30) is almost double of the other autophagy factors tested (∼10-15), suggesting that ATG13 is one of the first autophagy factors recruited to ATG9 vesicles. This is consistent with the recently observed interaction between ATG13 and ATG9 ^66–68^ This work also suggested that ATG13 binds to two distinct regions in ATG9, one independent of ATG101 and a second promoted by an ATG101 induced conformational change in the HORMA domain of ATG13 ^67,68^. Our results demonstrate that ATG13 is recruited to phagophores in the absence of ATG101, potentially reflecting ATG13 binding to the ATG101 independent binding site on ATG9. The slow kinetics for this interaction observed *in vitro* ^67^, might be overcome by the high expression levels of ATG13 in cells ^9^, assuring that the number of ATG13 molecules in a conformation capable of binding ATG9 is sufficiently high at all times. The increase in the number of ATG13 foci under amino acid or glucose starvation suggests that the interaction between ATG9 and ATG13 is promoted under these conditions potentially by phosphorylation of ATG9 or ATG13 ^69^ (**Fig. 7**).

### AMPK activation prevents tethering of mobile phagophores

Once primed as the seed for autophagosomes, ATG9 vesicles become tethered to the lipid source membrane by ATG2A ^9,10^. ATG2A, a rod-shaped protein, facilitates lipid transfer from the membrane source to the vesicle, enabling its rapid expansion ^15,16^. The recruitment of ATG2A to the vesicle seed is mediated by WIPI3/4 ^16,70^, which recognizes PI3P enrichments on the vesicle membrane, and potentially by the ATG9-ATG13 complex (**Fig. 7**) ^67^. Once tethered, ATG2A-mediated lipid transfer enables the vesicle to assume the characteristic cup-shaped form of the phagophore, ready to engulf the cargo designated for degradation (**Fig. 7**) ^1^. Our previous work, employing single-particle tracking approaches, has demonstrated that the recruitment of the LC3 conjugation machinery takes place just prior to vesicle tethering, as evidenced by the similar mobility patterns of WIPI2 foci and ATG2A foci ^9^. WIPI2 plays a crucial role in facilitating the recruitment of the E3-like ligase ATG16-5-12 complex, thus enhancing LC3 conjugation (**Fig. 7**)^45^. Our results indicate that AMPK signaling inhibits autophagosome maturation by reducing vesicle tethering to the phagophore. We have previously shown that phagophores exist in two distinct mobility states. A tethered state (D = 0.001 µm^2^ s^-1^) and an untethered vesicle state (D = 0.01 µm^2^ s^-1^). Importantly, both populations were present in foci that colocalized with LC3 and those that did not, suggesting that vesicle tethering does not require LC3 conjugation. Among the autophagy-related proteins examined, WIPI2 foci exhibited altered mobility after glucose starvation and AMPK activation. Specifically, LC3-negative WIPI2 foci, which likely represents the step immediately preceding tethering, displayed significantly increased mobility when AMPK was activated. Furthermore, our results also suggest that the amount of WIPI2 recruited to the phagophores is reduced in the absence of glucose (**Fig. 3**). This observation is also consistent with WIPI2 recruitment to mobile ATG9 vesicles, which are expected to have a limited capacity to bind WIPI2, compared to an expanding tether phagophore (**Fig. 7**). McAlpine and colleagues similarly reported a lack of WIPI2 (and WIPI1) recruitment in mouse embryonic under glucose-starvation conditions, which was attributed to a reduction in PI3P accumulation ^62^.

In total, our results support a model in which AMPK activation after glucose starvation leads to the accumulation of mobile phagophores bound by WIPI2, which fail to tether to donor membranes, leading to a reduction in autophagosome maturation. In contrast, when AMPK signaling is inhibited, we observed an increase in the rate of phagophore maturation, measured as a decrease in the time frame of LC3 colocalization with ATG2 and WIPI2 (**Fig. 4C**). This suggests that AMPK interferes with the lipid transfer machinery. Future studies will have to address the precise target of AMPK that leads to the reduction of autophagosome maturation we observed in this study. It has been suggested that WIPI4, which can recruit ATG2 to phagophores, is a direct AMPK interactor that is released from the AMPK/ULK1 complex upon autophagy induction ^70^. Another possibility is that AMPK regulates ATG9-ATG13 mediated ATG2 recruitment.

A key unaddressed question is why AMPK activation simultaneously increases autophagy foci formation and inhibits autophagosome maturation. One explanation is that the primary function of non-selective autophagy is to provide molecular building blocks but not metabolites that can be shunted into ATP production. Autophagy consumes ATP during all its stages, and inhibition of autophagy could preserve ATP under conditions that lead to AMPK activation. At the same time AMPK could increase the potency of early autophagy factors to facilitate targeted autophagy and maintain the ability to regulate mitochondrial function via mitophagy.

In summary, our study sheds light on the inhibitory role of AMPK signaling in autophagosome maturation during nutrient stress. Glucose starvation leads to a reduction in autophagosome maturation, a phenomenon consistently observed across multiple autophagy factor proteins. Importantly, this maturation rate is not influenced by the number of phagophores formed but by the accumulation of LC3-positive autophagosomes. Our results indicate that AMPK activation, whether triggered by glucose depletion or a pharmacological agonist, is a key driver of this inhibition. This observation contrasts with some previous reports but clarifies the intricate role of AMPK in autophagy regulation under nutrient stress. There is an emerging interest in activating AMPK for enhancing brain energetics and thus prevent neurological diseases, including Parkinson’s and Huntington’s diseases ^71^. In cancer, AMPK’s role remains a subject of intense debate, particularly because many cancers downregulate AMPK. This phenomenon appears counterintuitive, given AMPK’s putative role as an activator of catabolic pathways implicated in cancer growth and survival. To effectively target AMPK as a therapeutic approach it will be critical to precisely define all of its contributions to regulating metabolic processes, including autophagy.

## Supporting information

K-FOCUS Code

## Acknowledgments

We thank Luke D. Lavis (Janelia Research Campus, Ashburn, VA, USA) for generously providing the Janelia fluor dyes. We acknowledge the Flow Cytometry core (Research Technology Support Facility, Michigan State University) and the Confocal Laser Scanning Microscopy core (Center for Advanced Microscopy, Michigan State University) for supporting our cell sorting and microscopy experiments. The order of the authors C.B and D.B. was decided by a randomization process and both authors contributed equally to the paper; co-first authors reserve the right to list themselves first on their curriculum vitae. This work was supported by a grant from the NIH (DP2 GM142307) to J.C.S.

## Author contributions

Conceptualization: C.B., D.B., and J.C.S.; Experiments: C.B., D.B., and G.I.P.; Data Analysis: C.B. and D.B.; Writing – Original Draft: C.B. and D.B.; Writing – Review and Editing: C.B., D.B., and J.C.S.

## Materials and Methods Cell Lines

All cell lines used in this study were derived from human bone osteosarcoma epithelial cells (U2OS, ATCC HTB-96). The monoclonal cell lines edited to express the HaloTag at the endogenous loci of the autophagy factors ATG13, WIPI2, ATG2A, ATG9A and ULK1 were extensively characterized in our recent paper ^9^. The cell lines having knockouts of the genes belonging to the ULK1 complex (ULK1 KO, FIP200 KO and ATG101 KO) were also previously characterized ^9^. Cells were grown in RPMI cell culture media supplemented with 10% fetal bovine serum (FBS), 100 units/ml penicillin, 100 µg/ml streptomycin at 37°C with 5% CO_2_.

## Plasmid Construction and Genome Editing

GFP-tagged LC3 were stably expressed to parental HaloTagged cell lines (Halo-ATG2, Halo-ATG13, Halo-WIPI2, and ULK1-Halo), Halo-ATG13 knock-outs (ULK1 KO, ATG101 KO, FIP200 KO) by introducing the coding sequence at the AAVS1 safe-harbor locus (PPP1R12C) ^72^. Polyclonal GFP-LC3B expressing cell lines were obtained by fluorescence activated cell sorting (FACS) by selecting cells lying within 75^th^-95^th^ percentile of GFP-signal.

## Preparation of Growing Media and Treatment Conditions

The autophagy flux was characterized in in cells growing in four different media conditions: control RPMI media supplied with 10% dialyzed FBS (ThermoFisher Scientific), 25 mM glucose and amino acids (AA) (indicated as control), and media deprived of amino acids (AA), glucose (indicated as no glucose media in the paper), amino acids and FBS (analogues to an EBSS starvation media), and media deprived of all nutrients. All the media were prepared from a common base media (ThermoFisher Scientific) to which glucose, AA and FBS were added accordingly to the condition to be tested. All conditions media were filtered, and pH corrected before experiments and supplemented with 100 units/ml penicillin and 100 µg/ml streptomycin.

For the live-cell imaging experiments, 200,000 cells were seeded on glass coverslips (170 ± 5 μm, Schott) in control media. 24 hours after seeding, cells were labeled with JFX650 (100nM, 10 min), followed by dye bleeding (5 min); both steps were performed in control media. Afterward, cells were washed three times with PBS before being switched to the respective growing media conditions. The treatment duration for the different growing media conditions was 1 hour, except for the no glucose media, which had a treatment duration of 4 hours prior to imaging. For studying autophagy response after AMPK activity modulation, the compounds MK8722 (AMPK agonist) and BAY3827 (AMPK antagonist) were used. We designed a treatment/release experiment for addressing how cells responded to nutrient deficiency in the presence (treatment) or absence (control) of drugs after acute exposure to the drug itself. Cells were seeded at lower confluency (100,000 cells) on glass coverslips (170 ± 5 μm, Schott); 24h after seeding media was substituted with 1 mL of complete media containing MK8722 (1µM) or BAY3837 (1µM). After 18h of drug treatment, cells were labeled by adding 100 µl of media containing 1µM of JFX650 (final labeling concentration 100nM) for 10 min, followed by 5 min dye bleeding with control media supplemented with MK8722 (1µM) or BAY3837 (1µM). To analyze the response of cells to the AMPK agonist (MK8722), cells were washed three times with PBS and 1 mL of media deprived of amino acids and FBS (but not glucose) and supplemented with the agonist (1µM) was added. For the control experiment, the media (–AA/– FBS/+Glucose) was not supplemented with the agonist and replaced with fresh media after 30 min to guarantee complete removal of the AMPK agonist. Cells were imaged after 1 hour after the addition of agonist containing or control media. To analyze the response of cells to the AMPK antagonist (BAY3837) an identical approach was used, the only difference being the starvation media, which did not contain any nutrient (–AA/– FBS/-Glucose). For each live-cell imaging experiment, three biological replicates were performed and analyzed, with a total of 90-150 cells imaged per experiment.

## Live-cell Microscopy

Live cell imaging experiments were carried out using an Olympus microscope (IX83), with an X-Cite TURBO multi-wavelength LED illumination system (Excelitas Technologies). The microscope is equipped with an environmental chamber (cellVivo) to control humidity, temperature, and CO_2_ level, a 60× TIRF oil-immersion objective (Olympus UPlanApo, NA = 1.50), and the appropriate excitation and emission filters. The microscope is equipped with two Hamamatsu Orca Fusion BT sCMOS cameras attached to a Twin-cam beam splitter (Cairn Research). All live cell imaging was carried out at 37°C, and under 5% CO_2_-containing humidified air. For imaging the HaloTagged autophagy proteins labeled with JFX650 dye, the 630 nm LED source was set at 100% power. The GFP-edited autophagy marker LC3 was imaged with a 475 nm LED source at 30% laser power. Both optical settings led to negligible photobleaching across the entire experiment. Cells were imaged for 8 minutes total time, at 1 seconds per frame (total number of frames = 480) with 50 ms exposure time for both LED sources. Alternatively, we used an i3 spinning disc confocal microscope equipped with a CSU-W1 confocal spinning disk system (Yokogawa), five laser lines (100 mW 445nm, 150 mW 488nm, 175 mW 515 nm, 160 mW 561nm, and 140 mW 638 nm), a Prime 95B sCMOS camera (Photometrics), a 63X oil-immersion objective (Zeiss C Plan-Apo, NA = 1.4), and an incubation chamber to control humidity, temperature, and CO_2_ level. All live cell imaging was carried out at 37°C, and under 5% CO_2_-containing humidified air.

## K-FOCUS Imaging Analysis Pipeline

In the following sections, we will describe the K-FOCUS quantitative imaging pipeline with its three different steps: the CellPose-based segmentation step ^38^, the autophagy factor foci tracking using TrackIT ^40^, and the foci co-diffusion analysis algorithm.

### Cellular segmentation using CellPose

CellPose was trained to segment U2OS cells stably expressing GFP-LC3 from the AAVS1 locus and Halo-ATG9A expressed from the endogenous ATG9 locus. For the GFP-LC3, a training set was genrated by manually annotating >100 single-frame images obtained using the Olympus microscope described above. For Halo-ATG9A, single-frame images were obtained using the spinning-disk confocal microscope described above. The manual annotation and generation of the models were performed in the GUI version of CellPose and a preliminary segmentation was obtained using a pretrained model provided with the segmentation package. CellPose is robust to difference on cell confluency; however, optimal cellular segmentation was achieved when 200-300k cells were seeded 24h before imaging, corresponding to 50-60% plate confluency for U2OS cells.

Before cellular segmentation, we renamed the single-channel files to contain ‘C1’ (GFP-LC3) and ‘C2’ (HaloTagged autophagy factor) as part of their filename. Proper file renaming is essential for the pipeline to work since the image files will be recalled by the subsequent analysis algorithms. For dual-color imaging, only one channel has to be segmented, and the segmentation propagated to the second channel. In our experiments, cells were segmented in the GFP-LC3 channel. For performing cellular segmentation of large datasets, CellPose was implemented as a Python script using PyCharm environment (JetBrain). The script efficiently processes live-cell imaging videos (in TIFF format) by segmenting a single frame due to the negligible X-Y drift observed in our 8-minutes imaging experiments. The outputs of the cellular segmentation are single-cell ROI outlines. The script calls MatLab as computational engine using the MatLab Engine API package to convert the ROI outlines into ROI files to be implemented in the foci tracking algorithm.

### Single-cell Foci tracking using TrackIT

TrackIt is a GUI integrative tracking and analysis MatLab-based software developed by Gebhardt group ^40^. Movies were loaded in TrackIt, and ROI were imported and visually inspected to correct for misassigned cells. TrackIt fits single particle to 2D-Gaussian wavelet functions and links spots using a *nearest neighbor algorithm*. For our tracking analysis, we used the following optimized settings: threshold 1.9; tracking radius 5; minimum track length 5; gap frames 5. For tracking the LC3 foci, the threshold was set at 2.1. Tracking files were saved as a batch file containing single-cell tracking data.

### K-FOCUS co-diffusion analysis

K-FOCUS was developed to analyze foci co-diffusion and kinetics in 2-color live-cell imaging at a single-cell resolution. The core software consists in two scripts, *K-Focus_GUI* and *TrackColocalization_2CH. K-Focus_GUI* uses a GUI application for selecting batches files ^73^ for both channels and creates output folders for each condition tested. In this step, the algorithm allows for customization by specifying the minimum frame length of the foci tracks, which was set to 5 for the experiments discussed in this paper. The algorithm then performs background intensity correction for all the tracks. For calculating foci intensity, we centered an 11×11 pixel box on the particle coordinated obtained by TrackIt. We then divided this box into two subregions. The central 5×5 pixel box – corresponding to the foci signal – was used to obtain the intensity of the particle as the max intensity (I_max_). The pixel intensity of the remaining subregion surrounding the 5×5 pixel area was averaged to compute the background intensity (I_background_). The corrected particle intensity (I_max, corr_) was calculated as following I_max, corr_ = I_max_ –I_background_. Finally, for each foci track, the algorithm calculates the foci intensity by averaging the I_max, corr_ associated with each localization. At this stage, the algorithm computes the mean distance of the foci track from the ROI edge (cell border), as well as the ROI area (cell area). Once this step is complete, *K-FOCUS_GUI* will store single-cell analysis for each condition in a separate folder.

Next, *TrackColocalization_2CH* performs the colocalization analysis in ‘C1’ and ‘C2’. The algorithm runs multiple instances in parallel and saves each experimental condition as separate output file. To define two foci as colocalized, two parameters are considered: the *minimum distance* between the foci at any point during their lifetime, and the *extent of colocalization*, which measures the duration of time that the two foci are colocalized within that distance. For our analysis, the minimum distance was set at 3 pixel (0.65 μm for our camera), and the minimum extent of colocalization to 10 seconds. The algorithm further calculates several metrics at single-cell resolution, including the count of C2-positive foci (HaloTag autophagy factor foci not colocalized with GFP-LC3), the count of C2-negative foci (HaloTag autophagy factor foci colocalized with GFP-LC3), the conversion ratio (fraction of C2-positive foci out of the total C1 foci), and the lifetime of both fractions. These metrics are then stored in a MatLab structure for further analysis.

Additional filtering algorithms are applied to exclude preexisting foci within the initial 5 seconds. This step helps in calculating the ‘firing’ rate, which quantifies the number of newly formed foci per cell per minute. Moreover, we examined the extent of co-diffusion by assessing the duration of foci before and after the appearance of the C1 signal (i.e., before and after GFP-LC3B appearance). This analysis provides insights into the length of foci pre– and post-appearance of the C1 signal.

## Single-Cell Threshold– and Objective-based Foci Colocalization Analysis

To prepare the movies for the Pearson, Manders, and SODA colocalization algorithms, the channels Halo-ATG and GFP-LC3 were merged, and the first frame was isolated and segmented using an adapted script calling the CellPose API with the output being the ROIs per image in an Imagej format. Next, using a custom Imagej script the signal outside the ROI was then deleted, and a single image was saved per cell. A colocalization batch analysis was then designed in ICY ^74^, which separated the channels, detected the foci within individual cell in both channels using ICY’s innate detector algorithms. We used the SODA suite ^34^ to perform the Pearson, Manders, and SODA colocalization analysis.

## Step-size analysis

The step-size distributions obtained from single-particle tracking were fitted to a probability distribution to obtain the diffusion coefficients ^56^. The equation used for the fitting was the following:

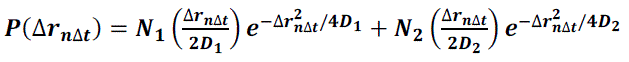

In this equation **P**(Δ_*r*Δ*t*_) is the probability of the protein displacing with a certain step-size. In our analysis, an autophagy factor focus can be associated to a mobile (dynamic) ATG9A vesicle, or to a tethered (static) ATG9A vesicle. *N*_1_ is the number of observed steps associated with the population 1 (dynamic) having a diffusion coefficient *D*_1_, and *N*_2_ is the number of observed steps associated with the population 2 (static) having a diffusion coefficient *D*_2_. For fitting the step-size distribution, a custom algorithm was written in MatLab. The algorithm computed the histogram using the Freedman-Diaconis rule ^75^ to select the optimal bin for each experiment and performed the fitting using a non-linear least square method with 1000 max iterations.

## Quantification and statistical analysis

Data are expressed as mean ± confidence interval (90-10) unless stated otherwise. Statistical differences were assessed using a two-tail t-test or one-way ANOVA at p < 0.05 significance level followed by Bonferroni post-hoc test. All the statistical analyses were performed using OriginPro (v. 2023b, OriginLab).

## Supplemental movies legends

**Movie S1**. U2OS cells expressing LAMP1-mNeonGreen and Halo-ATG9A (JFX650) imaged at 1 frame per second using a spinning disc confocal acquired with Photometrics Prime 95B sCMOS camera, either immediately after Halo-ATG9A labeling (top row), or 24 hours after Halo-ATG9A labeling (bottom row).

**Movie S2.** U2OS cells expressing Halo-ATG13(JFX650, left column, magenta in merge, right column) and GFP-LC3B (middle column, green in merge left column) imaged at 1 fps using widefield illumination and a Hamamatsu BT fusion camera (2×2 binning) cultured in complete media (top row) and EBSS (bottom row).

**Movie S3.** U2OS cells expressing Halo-WIPI2 (JFX650, left column, magenta in merge, right column) and GFP-LC3B (middle column, green in merge right column) imaged at 1 fps using widefield illumination and a Hamamatsu BT fusion camera (2×2 binning) cultured in different media conditions (rows, top to bottom: Complete Media; No Amino Acids; No Amino Acids and FBS; No Glucose; No Amino Acids, FBS, and Glucose).

**Movie S4.** U2OS cells expressing Halo-ATG13(JFX650, left column, magenta in merge, right column) and GFP-LC3B (middle column, green in merge left column) imaged at 1 fps using widefield illumination and a Hamamatsu BT fusion camera (2×2 binning) cultured in different media conditions (rows, top to bottom: Complete Media; No Amino Acids; No Amino Acids and FBS; No Glucose; No Amino Acids, FBS, and Glucose).

**Movie S5.** U2OS cells expressing Halo-ULK1 (JFX650, left column, magenta in merge, right column) and GFP-LC3B (middle column, green in merge left column) imaged at 1 fps using widefield illumination and a Hamamatsu BT fusion camera (2×2 binning) cultured in different media conditions (rows, top to bottom: Complete Media; No Amino Acids; No Amino Acids and FBS; No Glucose; No Amino Acids, FBS, and Glucose).

**Movie S6.** U2OS cells expressing Halo-ATG2A (JFX650, left column, magenta in merge, right column) and GFP-LC3B (middle column, green in merge left column) imaged at 1 fps using widefield illumination and a Hamamatsu BT fusion camera (2×2 binning) cultured in different media conditions (rows, top to bottom: Complete Media; No Amino Acids; No Amino Acids and FBS; No Glucose; No Amino Acids, FBS, and Glucose).

**Movie S7.** U2OS cells expressing Halo-ATG13 and GFP-LC3B (top row), Halo-ULK1 and GFP– LC3B (second row from the top), Halo-WIPI2 and GFP-LC3B (third row from the top), and Halo– ATG2A and GFP-LC3B (bottom row) in media lacking amino acids and FBS but in the presence of glucose without the AMPK agonist MK8722. Imaged at 1 fps using widefield illumination and a Hamamatsu BT fusion camera (2×2 binning). HaloTagged protein in the left column (magenta in the merge, right column) and GFP-LC3B in the middle column (green in the merge, right column).

**Movie S8.** U2OS cells expressing Halo-ATG13 and GFP-LC3B (top row), Halo-ULK1 and GFP– LC3B (second row from the top), Halo-WIPI2 and GFP-LC3B (third row from the top), and Halo– ATG2A and GFP-LC3B (bottom row) in media lacking amino acids and FBS but in the presence of glucose with the AMPK agonist MK8722. Imaged at 1 fps using widefield illumination and a Hamamatsu BT fusion camera (2×2 binning). HaloTagged protein in the left column (magenta in the merge, right column) and GFP-LC3B in the middle column (green in the merge, right column).

**Movie S9.** U2OS cells expressing Halo-ATG13 and GFP-LC3B (top row), Halo-ULK1 and GFP– LC3B (second row from the top), Halo-WIPI2 and GFP-LC3B (third row from the top), and Halo– ATG2A and GFP-LC3B (bottom row) in media lacking amino acids, FBS, and glucose without the AMPK inhibitor BAY3827. Imaged at 1 fps using widefield illumination and a Hamamatsu BT fusion camera (2×2 binning). HaloTagged protein in the left column (magenta in the merge, right column) and GFP-LC3B in the middle column (green in the merge, right column).

**Movie S10.** U2OS cells expressing Halo-ATG13 and GFP-LC3B (top row), Halo-ULK1 and GFP–LC3B (second row from the top), Halo-WIPI2 and GFP-LC3B (third row from the top), and Halo-ATG2A and GFP-LC3B (bottom row) in media lacking amino acids, FBS, and glucose with the AMPK inhibitor BAY3827. Imaged at 1 fps using widefield illumination and a Hamamatsu BT fusion camera (2×2 binning). HaloTagged protein in the left column (magenta in the merge, right column) and GFP-LC3B in the middle column (green in the merge, right column).

**Movie S11.** U2OS cells with a ULK1 knockout expressing Halo-ATG13 (JFX650, left column, magenta in merge, right column) and GFP-LC3B (middle column, green in merge left column) imaged at 1 fps using widefield illumination and a Hamamatsu BT fusion camera (2×2 binning) cultured in different media conditions (rows, top to bottom: Complete Media; No Amino Acids and FBS; No Glucose).

**Movie S12.** U2OS cells with a FIP200 knockout expressing Halo-ATG13 (JFX650, left column, magenta in merge, right column) and GFP-LC3B (middle column, green in merge left column) imaged at 1 fps using widefield illumination and a Hamamatsu BT fusion camera (2×2 binning) cultured in different media conditions (rows, top to bottom: Complete Media; No Amino Acids and FBS; No Glucose).

**Movie S13.** U2OS cells with a ATG101 knockout expressing Halo-ATG13 (JFX650, left column, magenta in merge, right column) and GFP-LC3B (middle column, green in merge left column) imaged at 1 fps using widefield illumination and a Hamamatsu BT fusion camera (2×2 binning) cultured in different media conditions (rows, top to bottom: Complete Media; No Amino Acids and FBS; No Glucose).

## Supplemental Material

**Figure S1.**
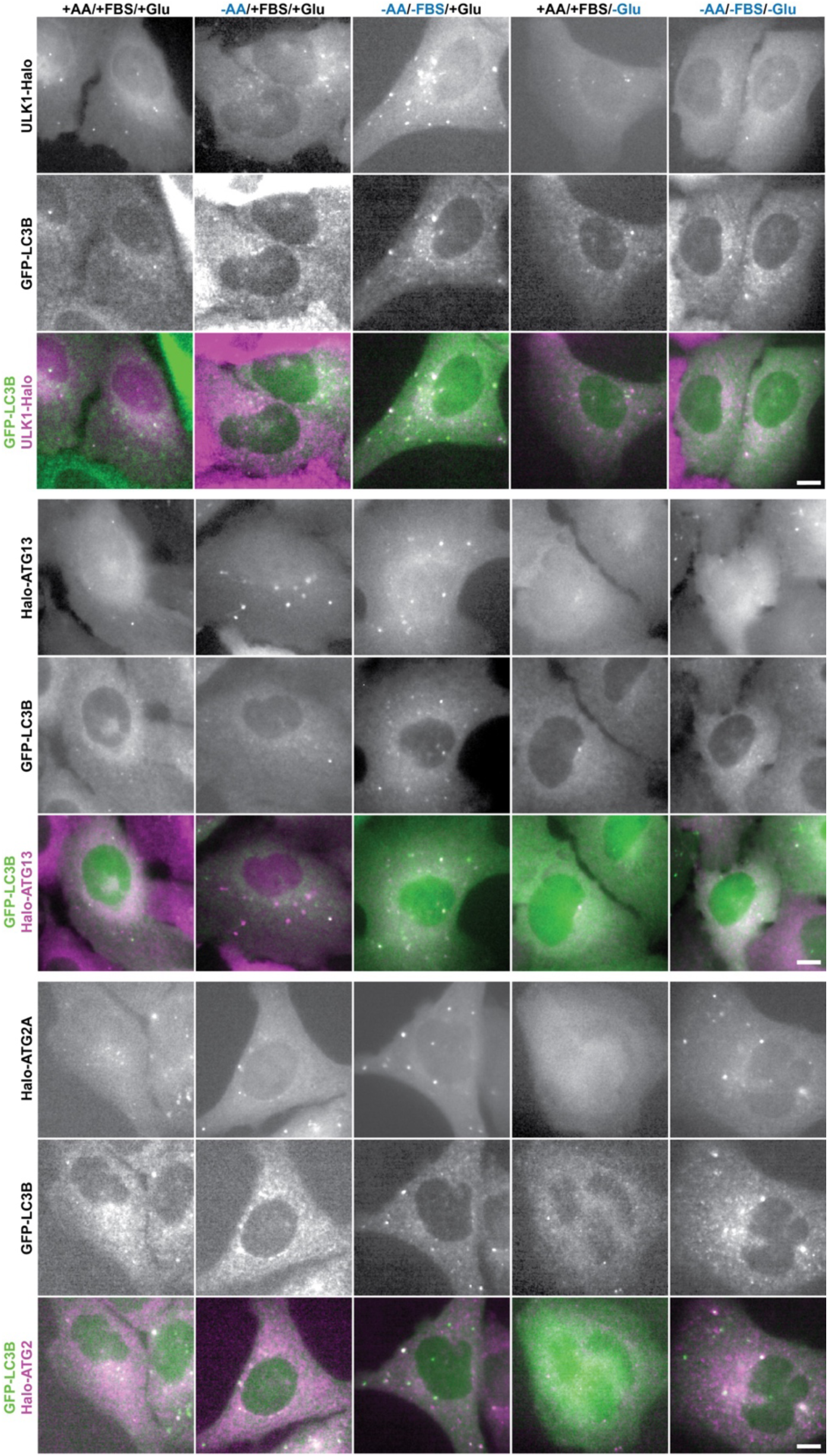
K-FOCUS analysis of the kinetics of autophagy factor foci formation under nutrient starvation. Live-Cell images of ULK1-Halo, Halo-ATG13, or Halo-ATG2A and GFPLC3B in control and starvation medium (Scale bar = 5 μm).

**Figure S2.**
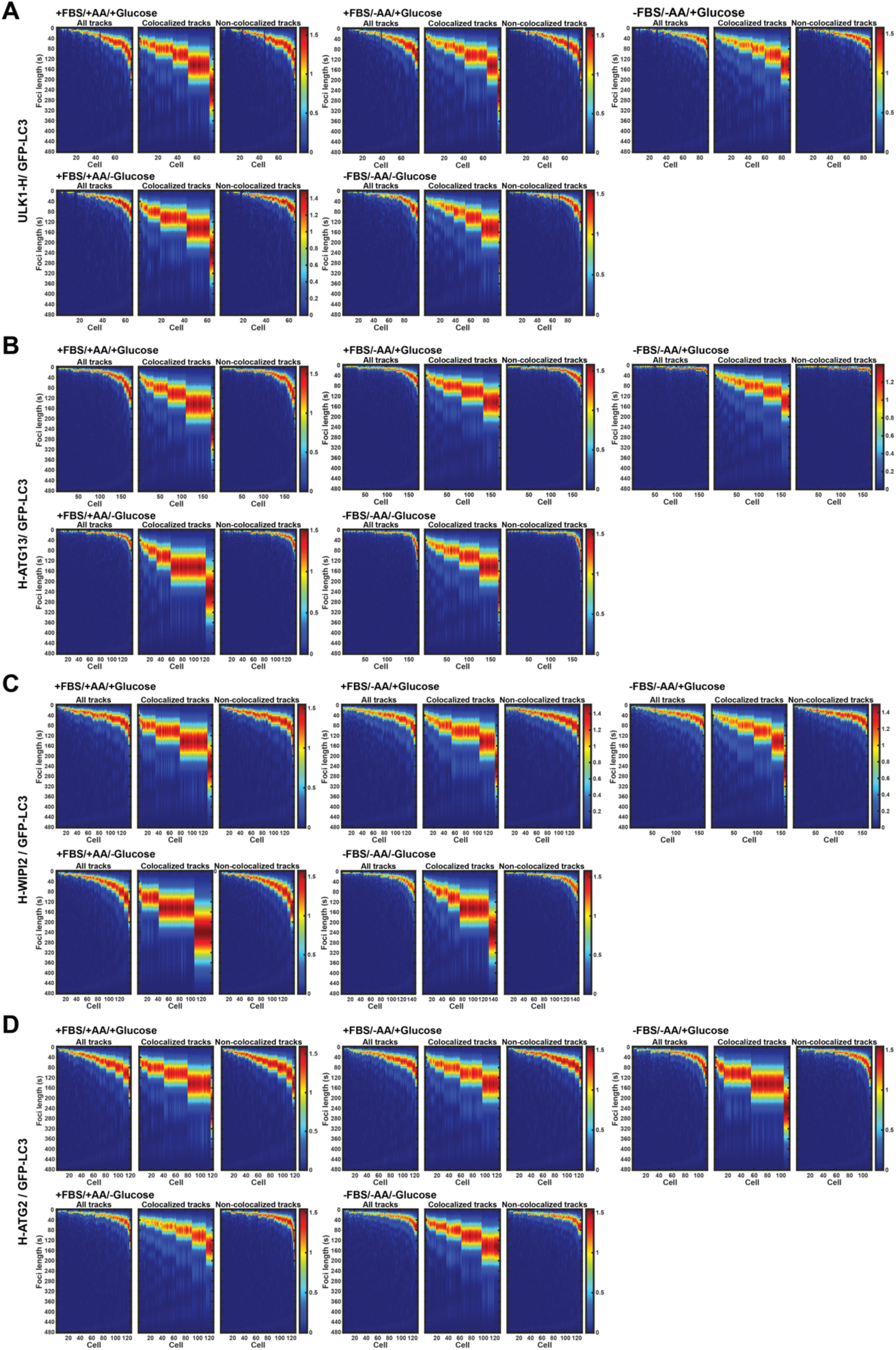
K-FOCUS analysis of the kinetics of autophagy factor foci formation under nutrient starvation. (**A-D**) Kernel density plots of the foci lifetimes per cell for (**A**) ULK1-Halo, (**B**) Halo-ATG13, (**C**) Halo-WIPI2, and (**D**) Halo-ATG2 in the indicated media conditions.

**Figure S3.**
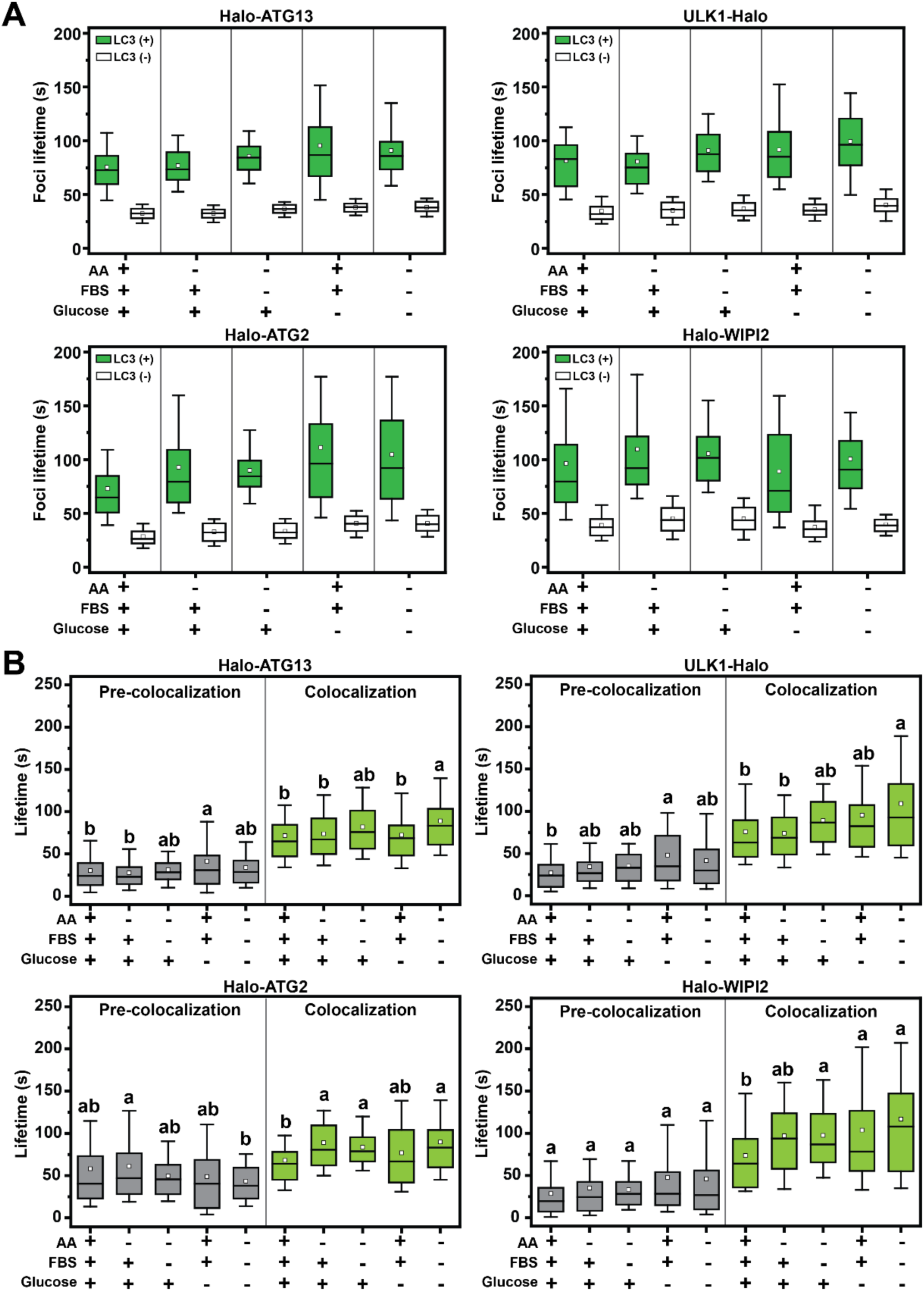
K-FOCUS analysis of autophagy factor foci lifetimes. (**A**) Lifetimes of LC3B positive and LC3B negative trajectories of Halo-ATG13, ULK1-Halo, Halo-WIPI2, and Halo-ATG2A in different media conditions. **(B)** Lifetimes of LC3B positive Halo-ATG13, ULK1-Halo, Halo-WIPI2, and Halo-ATG2A trajectories before and during colocalization with LC3B in different media conditions.

**Figure S4.**
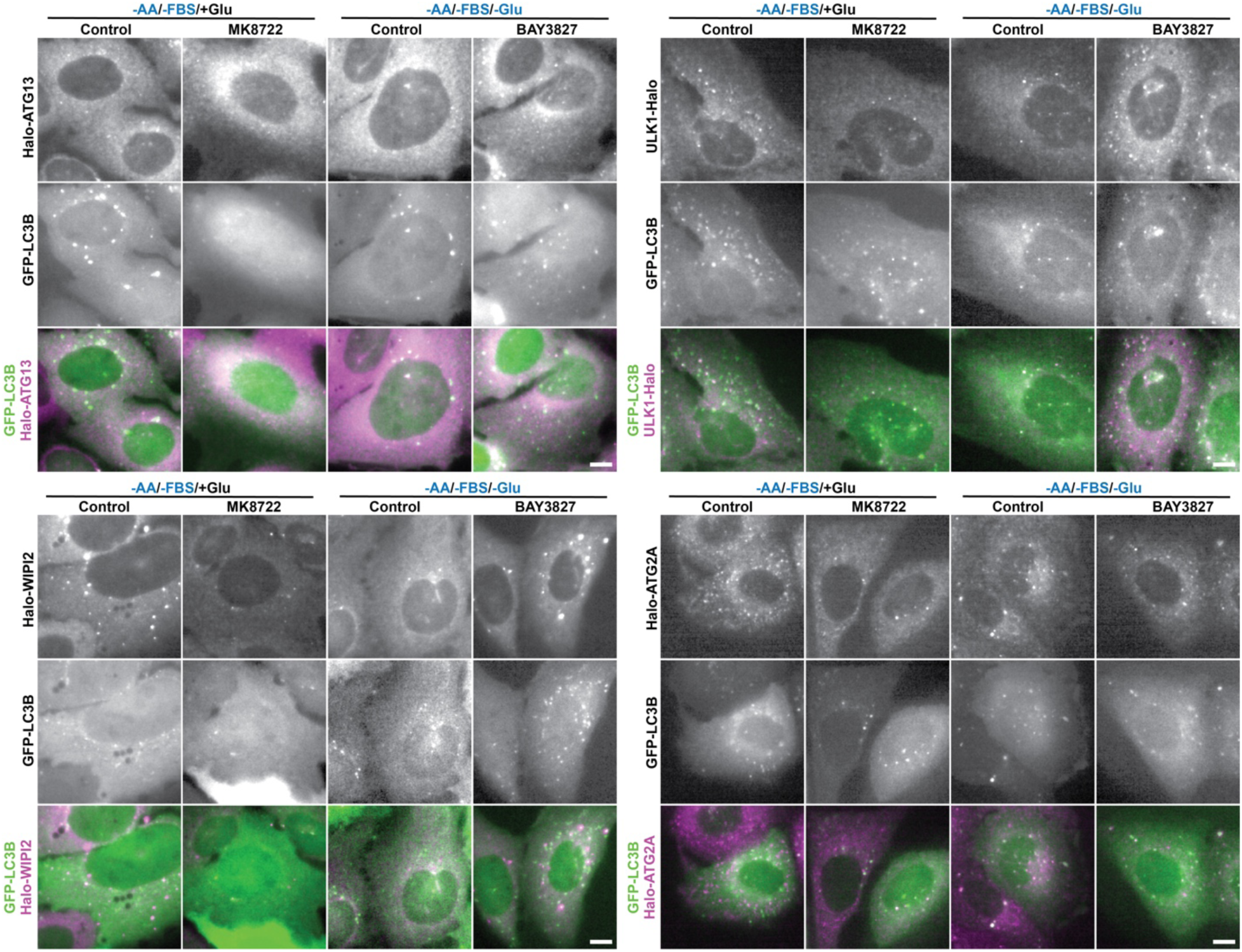
AMPK inhibits autophagosome maturation. Live-Cell images of ULK1-Halo, Halo-ATG13, or Halo-ATG2A and GFP-LC3B in the indicated media conditions in the presence and absence of the AMPK activator MK8722 or the AMPK inhibitor BAY3827 (Scale bar = 5 μm).

**Figure S5.**
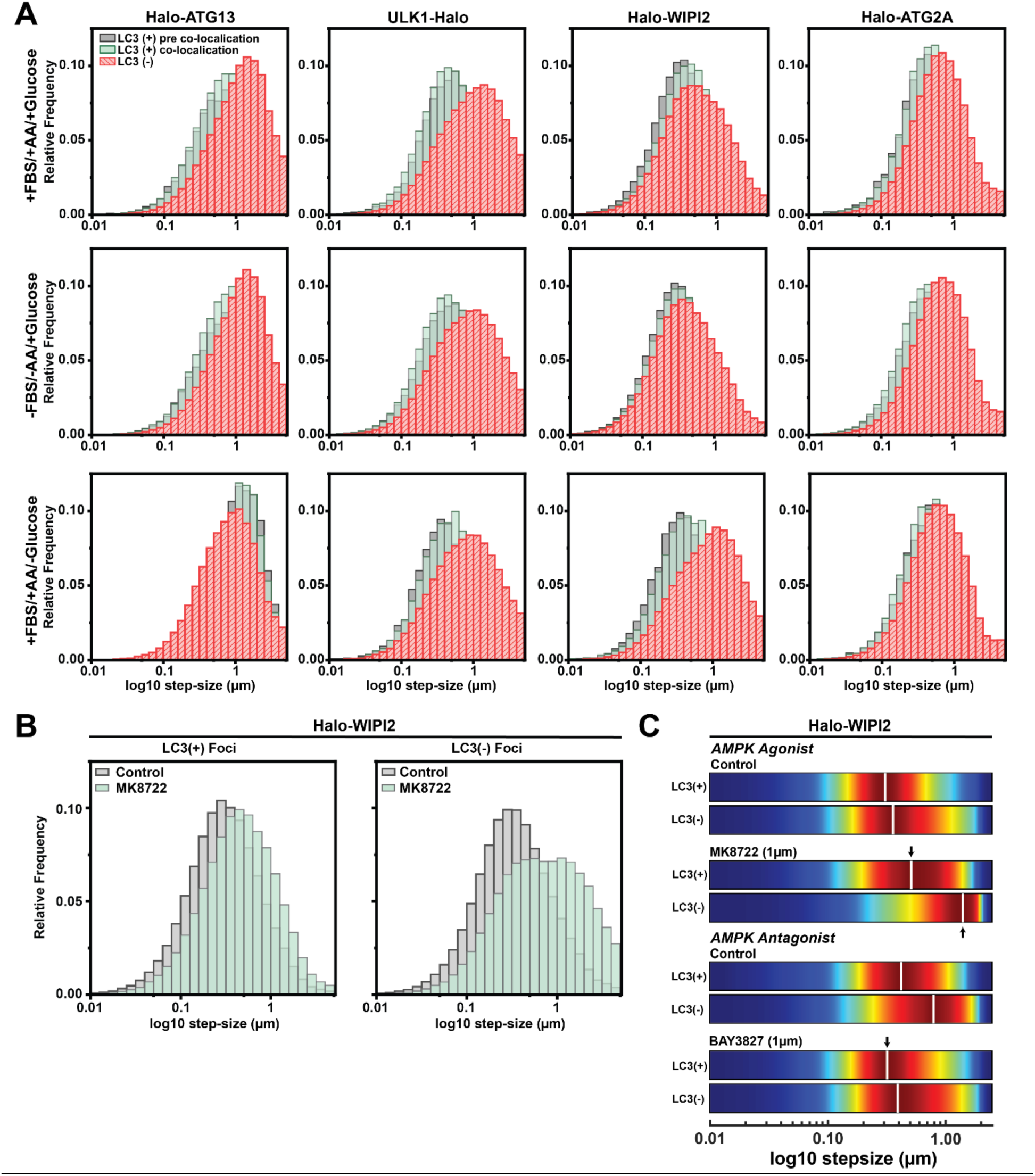
Diffusion dynamics of autophagy factor foci. (**A**) Step size distributions of LC3B negative and LC3B positive autophagy factor trajectories before and during colocalization with LC3B in indicated media conditions. **(B)** Step size distributions of LC3B negative and LC3B positive Halo-WIPI2 foci in the presence and absence of the AMPK agonist MK8722. (**C**) Kernel density plot of the step size distribution of LC3B negative and LC3B positive Halo-WIPI2 foci in the presence and absence of the AMPK agonist MK8722 (in media with glucose) or the presence and absence of the AMPK inhibitor BAY3827 (in media lacking glucose).

**Figure S6.**
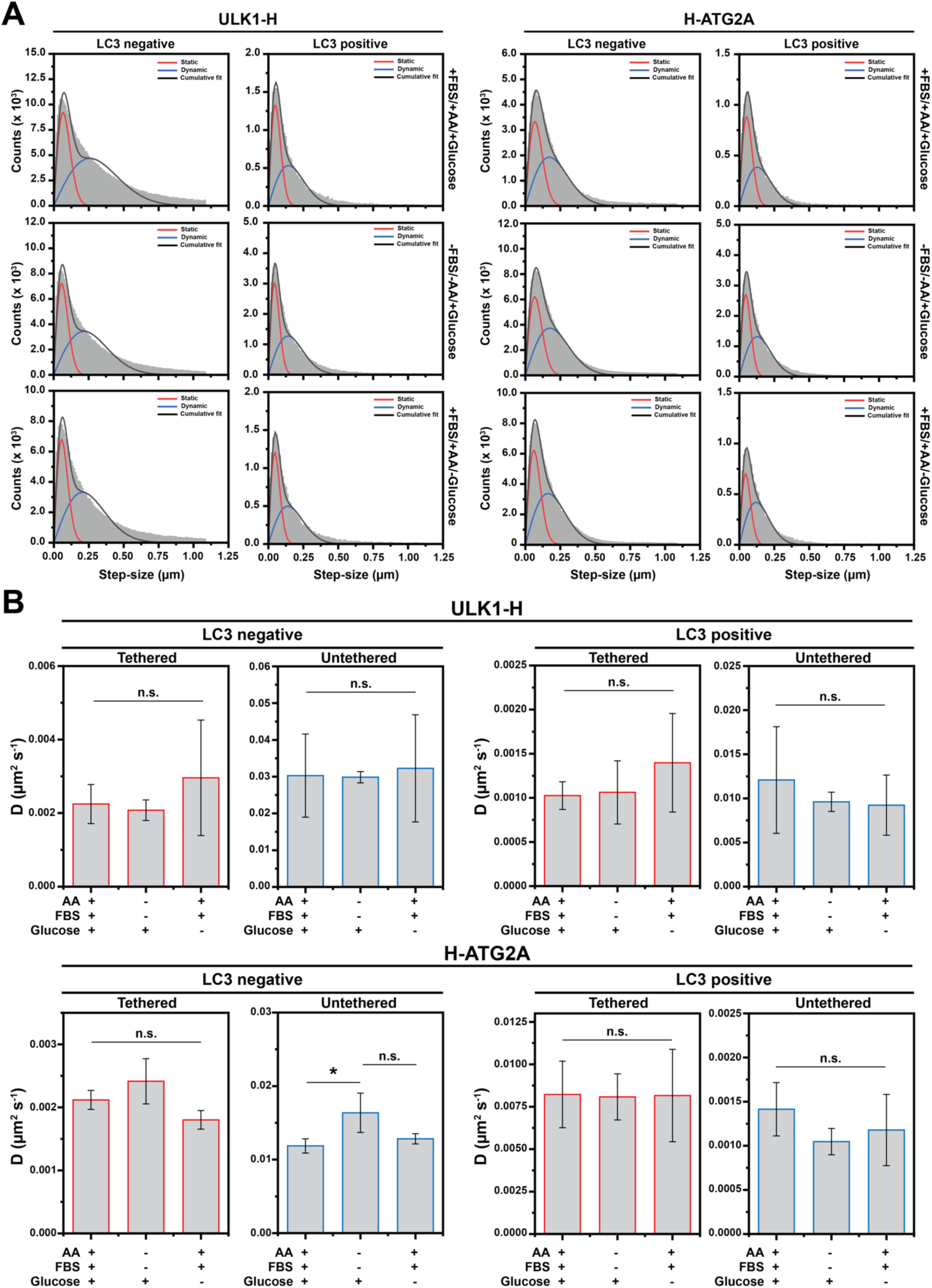
Diffusion dynamics of autophagy factor foci. (**A**) Step size distributions of LC3B positive and negative ULK1-Halo, and Halo-ATG2A trajectories fit with a two-state diffusion model (black line) encompassing tethered (red) and untethered (blue) populations in the different media conditions. (**B**) Diffusion coefficients of the tethered and untethered populations of LC3B positive and negative ULK1-Halo, and Halo-ATG2A trajectories in the different media conditions, derived from the fits shown in Fig. S6A.

