## Supplementary material for "AMPK Regulates Phagophore-to-Autophagosome Maturation": K-FOCUS Code: cellposetoMat.m

```
function cellposetoMat(filename,dir,dir2)

newdir = strcat(dir,'ROI');
dir_filename = strcat(dir2,filename);

if exist(newdir,'dir')
    % OK, output_path exists and is a directory (== 7). 
    disp('The given output foder exists. MATLAB workspaces will be save to:');
    disp(dir);
else
    mkdir(newdir);
    disp('The given output folder did not exist, but was just created. MATLAB workspaces will be save to:');
    disp(dir);
end

data = readmatrix(dir_filename,'Delimiter',{','}, 'Range', 'A1');
newfilename = strcat(newdir, '/', filename(1:end-16),'.roi')
ATGfilename = strcat(newdir, '/', filename(1:end-17),'1.roi')
subROI{1,1}{1,1}={};
ROI{1,1} = [.5 .5; .5 512.5; 512.5 512.5; 512.5 .5; .5 .5]; %Olympus
%ROI{1,1} = [.5 .5; .5 600.5; 600.5 600.5; 600.5 .5; .5 .5];

for k = 1:size(data,1);
    
    datatemp=data(k,:)
    datatempx=datatemp(1:2:end-1)';
    datatempy=datatemp(2:2:end)';
    datatemp=horzcat(datatempx,datatempy);
    subROI{1,k}{1,1}{1,1}=datatemp;
        
end

save(newfilename,'ROI','subROI');
save(ATGfilename,'ROI','subROI');
```
