## Supplementary material for "AMPK Regulates Phagophore-to-Autophagosome Maturation": K-FOCUS Code: main.py

```
from typing import List
import numpy as np
import matplotlib.pyplot as plt
from cellpose import models, io
from cellpose.io import imread
import os
import matlab.engine
import tkinter
from tkinter import filedialog
import tifffile
import itertools

#obtain directory path
tkinter.Tk().withdraw() # prevents an empty tkinter window from appearing
dir = filedialog.askdirectory()+'/'
dir2 = dir+'singleframe/'

# Create a singleframe Directory to save frame 50 of movies
try:
    os.mkdir(dir+'singleframe')
    print("Directory " , dir+'singleframe' ,  " Created ")
except FileExistsError:
    print("Directory " , dir+'singleframe' ,  " already exists")

files = [f for f in os.listdir(dir) if f.endswith('C1.tif')]
#files: List[str] = [os.path.join(dir, file) for file in
#         os.listdir(dir) if file.endswith('.tif')]


x = [io.imread(dir+f) for f in files]

#save single frame videos by using itertools.count for dynamic name saving

counter = itertools.count(0)
[tifffile.imwrite(dir2 + f, x[next(counter)][2]) for f in files]

# model_type='cyto' or 'nuclei' or 'cyto2' alternatively pretrained_model='absolutepathtomodel'
# modeldir = 'C:/Users/David/Desktop/cellpose/models/U2OSCellSegmentation01'
# modeldir = 'C:/Users/David/.cellpose/models/LC3_Virus'
modeldir = 'C:/Users/barnabac/Desktop/CellPose trial/LC3B-BTFusion_4'
model = models.CellposeModel(pretrained_model=modeldir, net_avg=False)

# channels = [0,0]
# list of files
# PUT PATH TO YOUR FILES HERE!
#files: List[str] = [os.path.join(dir, file) for file in
#         os.listdir(dir) if file.endswith('.tif')]

x = [io.imread(dir2+f) for f in files]
nimg = len(x)

# define CHANNELS to run segementation on
# grayscale=0, R=1, G=2, B=3
# channels = [cytoplasm, nucleus]
# if NUCLEUS channel does not exist, set the second channel to 0
# channels = [[0,0]]
# IF ALL YOUR IMAGES ARE THE SAME TYPE, you can give a list with 2 elements
channels = [0, 0]  # IF YOU HAVE GRAYSCALE
# channels = [2,3] # IF YOU HAVE G=cytoplasm and B=nucleus
# channels = [2,1] # IF YOU HAVE G=cytoplasm and R=nucleus

# if diameter is set to None, the size of the cells is estimated on a per image basis
# you can set the average cell `diameter` in pixels yourself (recommended)
# diameter can be a list or a single number for all images
# Input diameter from GUI model
masks, flows, styles = model.eval(x, diameter=135, channels=channels, model_loaded=True)

files: List[str] = [os.path.join(dir2, file) for file in
         os.listdir(dir2) if file.endswith('.tif')]
io.save_masks(x, masks, flows, files, tif=True)

eng = matlab.engine.start_matlab()
#text_files: List[str] = [os.path.join('C:/Users/David/Desktop/cellpose/singleframemovie/', file) for file in os.listdir('C:/Users/David/Desktop/cellpose/singleframemovie/') if os.path.isfile(os.path.join('C:/Users/David/Desktop/cellpose/singleframemovie/', file)) and file.endswith(".txt")]

text_files = [f for f in os.listdir(dir2) if f.endswith('.txt')]

[eng.cellposetoMat(f,dir,dir2,nargout=0) for f in text_files]
```
