## Supplementary material for "AMPK Regulates Phagophore-to-Autophagosome Maturation": K-FOCUS Code: multiread.py

```
import numpy as np
import matplotlib.pyplot as plt
from cellpose import models
from cellpose.io import imread
import os

def load_images_from_folder('C:/Users/David/Desktop/cellpose/singleframemovie'):
    images = []
    for filename in os.listdir('C:/Users/David/Desktop/cellpose/singleframemovie'):
        img = imread(os.path.join(folder,filename))
        if img is not None:
            images.append(img)
    return images
```
