## Supplementary material for "AMPK Regulates Phagophore-to-Autophagosome Maturation": K-FOCUS Code: KFocus_Master_GUI.m

```
%% SpotAnalysis script by Carlo Barnaba and David Broadbent 04/26/2023 %%

close all
clear all

FileList = uipickfiles;

for iter = 1:length(FileList);

selectedFile = FileList{iter};
[filepath,name,ext] = fileparts(selectedFile)

analysisFolder = fullfile(filepath, 'Kfocus_analysis'); % replace with the filepath to your folder
if ~exist(analysisFolder, 'dir')
    mkdir(analysisFolder);
    disp('Directory created');
else
    disp('Directory already exists');
end

% Check for settings file, proceed to dialog if not present
settingsFile = fullfile(filepath, 'KFocus_Settings.mat');
if exist(settingsFile, 'file') == 2
    load(settingsFile);
else
    disp('No Settings File Present, User Input Required');
    
    % Create a dialog box to get the number of conditions from the user
    numConditions = str2double(inputdlg('How many conditions do you want to assign?', 'User Input', 1));

    % Create a cell array to store the signed conditions
    settings.Conditions = cell(numConditions, 1);

    % Create a dialog box to get the conditions and minimum track length from the user
    prompt = {'Minimum track length:'};
    defaultAnswers = {'5'};
    dlgTitle = 'User Input';
    LineNo = 1;
    AddOpts.Resize = 'on';
    AddOpts.WindowStyle = 'normal';
    AddOpts.Interpreter = 'tex';

    for i = 1:numConditions
        prompt{i + 1} = ['Condition ', num2str(i), ':'];
        defaultAnswers{i + 1} = '';  % Prefilled values are initially empty
    end

    % Opens dialog menu
    userInputs = inputdlgcol(prompt, dlgTitle, LineNo, defaultAnswers, AddOpts, 2);

    % Extract the minimum track length
    settings.minTrackLength = str2double(userInputs{1});

    % Loop through each condition and process the input
    for i = 1:numConditions
        settings.Conditions{i} = userInputs{i + 1};
        currentCondition = settings.Conditions{i};

        % Check if a directory exists for the current condition
        conditionDir = fullfile(analysisFolder, currentCondition);
        if ~exist(conditionDir, 'dir')
            % Create the directory if it doesn't exist
            mkdir(conditionDir);
            disp(['Directory created for condition ', currentCondition]);
        else
            % Display a message if the directory already exists
            disp(['Directory already exists for condition ', currentCondition]);
        end
    end

    disp(['Minimum track length set to ', num2str(settings.minTrackLength)]);

    % Save the settings to the settings file
    save(settingsFile, 'settings', '-mat');
end

data = load(selectedFile);
batch = data.batch;
filesTable = data.filesTable;

batch = IntensityCorrectionParallel(batch);

modifiedFilename = strcat(selectedFile(1:end-4), '_IntensCorrected.mat');
save(modifiedFilename, 'batch', 'filesTable', '-mat');

BatchToSingleCellParallel(modifiedFilename , analysisFolder, settings)

modifiedFilename2 = fullfile(filepath, 'KFocus_Settings.mat');
save(modifiedFilename2, 'settings', '-mat');

end
```
