## Supplementary material for "AMPK Regulates Phagophore-to-Autophagosome Maturation": K-FOCUS Code: BatchToSingleCellParallel.m

```
function BatchToSingleCell(modifiedfilename,Analysis_folder,settings);

data = load(modifiedfilename);
batch = data.batch;
filesTable = data.filesTable;
dbstop if error

for movieIdx = 1:size(batch,1)

    trackSubRegionAssignment = batch(movieIdx).results.tracksSubRoi;
    nRegions = batch(movieIdx).results.nSubRegions+1;

    tracksInSubRoiIdx = batch(movieIdx).results.tracksSubRoi;
    title = strrep(filesTable{movieIdx,1}, '.tif','-');
                
tic
for subRegionIdx = 1:nRegions-1
    data = struct('TrackID', [],'xy', [], 'stepsize', [], 'frame',[], 'TimeStamp',[], 'movie',[], 'cellnumber', [], 'PeakIntensity', [], 'ROISize', [], 'minDistEdge', []);
    tracksInSubRegion  = subRegionIdx-1 == trackSubRegionAssignment;
    temptracks = batch(movieIdx).results.tracks(tracksInSubRegion);


    %calculate ROI Size
    Xcoordinates = rmmissing(batch(movieIdx).subROI{1, subRegionIdx}{1,1}{1,1}(:,1));
    Ycoordinates = rmmissing(batch(movieIdx).subROI{1, subRegionIdx}{1,1}{1,1}(:,2));
    ROISize = round(polyarea(Xcoordinates, Ycoordinates));
    
    %filter out small tracks and reasign temptracks using the filter if
    %there are any tracks at all
   for iter = 1:size(temptracks,2)
   filteredtracks(iter) = size(temptracks{1,iter},1) > 5;
   end
   

    
    if 1:size(temptracks,2) > 0
    filteredtracks = filteredtracks == 1; 
    temptracks = temptracks(filteredtracks);
    end
    
    if 1:size(temptracks,2) > 0

    parfor k = 1:size(temptracks,2)
    tempxy = temptracks{1,k}(:,2:3);
    temp(k).TrackID = k;
    temp(k).xy = tempxy;
    temp(k).tempIntensity = temptracks{1,k}(:,12);
    
    %calculate stepsize
    steps = zeros(length(temptracks{1,k}(:,2:3))-1, 1);
    for kk = 1:(length(temptracks{1,k}(:,2:3))-1)
        temptrack = temptracks{1,k}(:,2:3);
        loc = temptrack(kk,1:2); % current particle location
        locplusone = temptrack(kk+1,1:2); % next location
        steps(kk) = markerdistance(loc, locplusone); % determine stepsize and store in steps vector
    end
    temp(k).stepsizetemp = steps;
    
    
    %calcaulte minDistEdge
    P = mean(tempxy);
    temp(k).minDistEdge = min(abs(p_poly_dist1(P(1), P(2), batch(movieIdx).subROI{1, subRegionIdx}{1, 1}{1, 1}(:,1), batch(movieIdx).subROI{1, subRegionIdx}{1,1}{1,1}(:,2))));

    tempframe = temptracks{1,k}(:,1);
    temp(k).frame = tempframe;
    
    %change time depending on framerate 3 for 3 seconds
    temp(k).TimeStamp = tempframe*1;
    temp(k).movie = movieIdx;
    temp(k).cellnumber = subRegionIdx-1;

    end
    
    
    data = struct('TrackID', {temp.TrackID},'xy', {temp.xy}, 'stepsize', {temp.stepsizetemp}, 'frame', {temp.frame}, 'TimeStamp', {temp.TimeStamp}, 'movie', {temp.movie}, 'cellnumber', {temp.cellnumber}, 'PeakIntensity', {temp.tempIntensity}, 'ROISize', {ROISize}, 'minDistEdge', {temp.minDistEdge});
    
    else
    
    data = struct('TrackID', [],'xy', [], 'stepsize', [], 'frame',[], 'TimeStamp',[], 'movie',[], 'cellnumber', [], 'PeakIntensity', [], 'ROISize', [], 'minDistEdge', []);

    end
    
    
    
    for i = 1:length(settings.Conditions)
        
    if contains(lower(title), lower(settings.Conditions{i}))
       
        new_title = fullfile(Analysis_folder,settings.Conditions{i},'\',char(append(title,string(subRegionIdx-1),'.mat')))
        save(fullfile(new_title),'data');
    end
    end
       

    clear TrackID xy frame TimeStamp movie cellnumber tempIntensity ROISize minDistEdge filteredtracks stepsize temptracks temp
    toc
end

end
```
