## Supplementary material for "AMPK Regulates Phagophore-to-Autophagosome Maturation": K-FOCUS Code: inputdlgcol.m

```
function Answer=inputdlgcol(Prompt, Title, NumLines, DefAns,Resize,NumCols)
%INPUTDLGCOL Input dialog box.
%  Answer = INPUTDLGCOL(Prompt) creates a modal dialog box that returns
%  user input for multiple prompts in the cell array Answer.  Prompt
%  is a cell array containing the Prompt strings.
%
%  INPUTDLGCOL uses WAITFOR to suspend execution until the user responds.
%
%  Answer = INPUTDLGCOL(Prompt,Title) specifies the Title for the dialog.
%
%  Answer = INPUTDLGCOL(Prompt,Title,LineNo) specifies the number of lines
%  for each answer in LineNo.  LineNo may be a constant value or a 
%  column vector having one element per Prompt that specifies how many
%  lines per input.  LineNo may also be a matrix where the first
%  column specifies how many rows for the input field and the second
%  column specifies how many columns wide the input field should be.
%
%  Answer = INPUTDLGCOL(Prompt,Title,LineNo,DefAns) specifies the default
%  answer to display for each Prompt.  DefAns must contain the same
%  number of elements as Prompt and must be a cell array.
%
%  Answer = INPUTDLGCOL(Prompt,Title,LineNo,DefAns,AddOpts) specifies whether
%  the dialog may be resized or not.  Acceptable values for AddOpts are 
%  'on' or 'off'.  If the dialog can be resized, then the dialog is
%  not modal.  
%
%  AddOpts may also be a data structure with fields Resize,
%  WindowStyle and Interpreter.  Resize may be 'on' or 'off'.
%  WindowStyle may be 'modal' or 'normal' and Interpreter may be
%  'tex' or 'none'.  The interpreter applies to the prompt strings.
%
%  Answer = INPUTDLGCOL(Prompt,Title,LineNo,DefAns,AddOpts,columnsize)
%  specifies the number of columns of edit boxes.  By default it is set 
%  to 1.
%
%  Examples:
%
%     prompt={'Enter the matrix size for x^2:','Enter the colormap name:'};
%     def={'20','hsv'};
%     dlgTitle='Input for Peaks function';
%     lineNo=1;
%     answer=inputdlgcol(prompt,dlgTitle,lineNo,def);
%
%     AddOpts.Resize='on';
%     AddOpts.WindowStyle='normal';
%     AddOpts.Interpreter='tex';
%     answer=inputdlgcol(prompt,dlgTitle,lineNo,def,AddOpts);
%
%     prompt={'Enter your first name:','Enter your street address:',...
%     'Enter your city and state and zip code:','Enter your last name:','Enter your apartment number:'};
%     def={'','','','','','',''};
%     dlgTitle='Input for address information';
%     largest_question_length = size((strvcat(prompt')),2);
%     lineNo=[1,largest_question_length+1];
%     AddOpts.Resize='on';
%     AddOpts.WindowStyle='normal';
%     AddOpts.Interpreter='tex';
%     answer=inputdlgcol(prompt,dlgTitle,lineNo,def,AddOpts,2);
%
%  See also TEXTWRAP, QUESTDLG, WAITFOR.

%  Loren Dean   May 24, 1995.
%  Copyright 1998-2002 The MathWorks, Inc.
%  $Revision: 1.58 $

%%%%%%%%%%%%%%%%%%%%%
%%% General Info. %%%
%%%%%%%%%%%%%%%%%%%%%
Black      =[0       0        0      ]/255;
LightGray  =[192     192      192    ]/255;
LightGray2 =[160     160      164    ]/255;
MediumGray =[128     128      128    ]/255;
White      =[255     255      255    ]/255;

%%%%%%%%%%%%%%%%%%%%
%%% Nargin Check %%%
%%%%%%%%%%%%%%%%%%%%
if nargin == 1 && nargout == 0
  if strcmp(Prompt,'InputDlgResizeCB')
    LocalResizeFcn(gcbf)
    return
  end
end

%********************** EDITTED ************
% Change the number of maximum inputs to 6
error(nargchk(1,6,nargin));
%*******************************************

error(nargoutchk(1,1,nargout));
if nargin==1
  Title=' ';
end

if nargin<=2, NumLines=1;end

if ~iscell(Prompt)
  Prompt={Prompt};
end

NumQuest=prod(size(Prompt));    

if nargin<=3
  DefAns=cell(NumQuest,1);
  for lp=1:NumQuest, DefAns{lp}=''; end
end

WindowStyle='modal';
Interpreter='none';
if nargin<=4
  Resize = 'off';
end

%********************** EDITTED ************
% Set up sizes for 5 and 6 inputs
if nargin>=5
    if isstruct(Resize)
        Interpreter=Resize.Interpreter;
        WindowStyle=Resize.WindowStyle;
        Resize=Resize.Resize;
    end
end

if nargin < 6
    NumCols = 1;
end
%*******************************************

if strcmp(Resize,'on')
  WindowStyle='normal';
end

% Backwards Compatibility
if isstr(NumLines)
  warning(['Please see the INPUTDLG help for correct input syntax.' 10 ...
           '         OKCallback no longer supported.' ]);
  NumLines=1;
end

[rw,cl]=size(NumLines);
OneVect = ones(NumQuest,1);
if (rw == 1 && cl == 2)
  NumLines=NumLines(OneVect,:);
elseif (rw == 1 && cl == 1)
  NumLines=NumLines(OneVect);
elseif (rw == 1 && cl == NumQuest)
  NumLines = NumLines';
elseif rw ~= NumQuest || cl > 2
  error('NumLines size is incorrect.')
end

if ~iscell(DefAns)
  error('Default Answer must be a cell array in INPUTDLG.');  
end

%%%%%%%%%%%%%%%%%%%%%%%
%%% Create InputFig %%%
%%%%%%%%%%%%%%%%%%%%%%%
FigWidth=300;FigHeight=100;
FigPos(3:4)=[FigWidth FigHeight];
FigColor=get(0,'Defaultuicontrolbackgroundcolor');
TextForeground = Black;
if sum(abs(TextForeground - FigColor)) < 1
    TextForeground = White;
end
InputFig=dialog(                               ...
               'Visible'         ,'off'      , ...
               'Name'            ,Title      , ...
               'Pointer'         ,'arrow'    , ...
               'Units'           ,'points'   , ...
               'UserData'        ,''         , ...
               'Tag'             ,Title      , ...
               'HandleVisibility','on'       , ...
               'Color'           ,FigColor   , ...
               'NextPlot'        ,'add'      , ...
               'WindowStyle'     ,WindowStyle, ...
               'Resize'          ,Resize       ...
               );
  

%%%%%%%%%%%%%%%%%%%%%
%%% Set Positions %%%
%%%%%%%%%%%%%%%%%%%%%
DefOffset=5;
SmallOffset=2;

DefBtnWidth=50;
BtnHeight=20;
BtnYOffset=DefOffset;
BtnFontSize=get(0,'FactoryUIControlFontSize');
BtnWidth=DefBtnWidth;

TextInfo.Units              ='points'   ;   
TextInfo.FontSize           =BtnFontSize;
TextInfo.HorizontalAlignment='left'     ;
TextInfo.HandleVisibility   ='callback' ;

StInfo=TextInfo;
StInfo.Style              ='text'     ;
StInfo.BackgroundColor    =FigColor;
StInfo.ForegroundColor    =TextForeground ;

TextInfo.VerticalAlignment='bottom';

EdInfo=StInfo;
EdInfo.Style='edit';
EdInfo.BackgroundColor=White;

BtnInfo=StInfo;
BtnInfo.Style='pushbutton';
BtnInfo.HorizontalAlignment='center';

% Determine # of lines for all Prompts
ExtControl=uicontrol(StInfo, ...
                     'String'   ,''         , ...    
                     'Position' ,[DefOffset                  DefOffset  ...
                                 0.96*(FigWidth-2*DefOffset) BtnHeight  ...
                                ]            , ...
                     'Visible'  ,'off'         ...
                     );
                     
WrapQuest=cell(NumQuest,1);
QuestPos=zeros(NumQuest,4);

for ExtLp=1:NumQuest
  if size(NumLines,2)==2
    [WrapQuest{ExtLp},QuestPos(ExtLp,1:4)]= ...
        textwrap(ExtControl,Prompt(ExtLp),NumLines(ExtLp,2));
  else
    [WrapQuest{ExtLp},QuestPos(ExtLp,1:4)]= ...
        textwrap(ExtControl,Prompt(ExtLp),80);
  end
end % for ExtLp

delete(ExtControl);
QuestHeight=QuestPos(:,4);

TxtHeight=QuestHeight(1)/size(WrapQuest{1,1},1);
EditHeight=TxtHeight*NumLines(:,1);
EditHeight(NumLines(:,1)==1)=EditHeight(NumLines(:,1)==1)+4;

%********************** EDITTED ************
% Get the total number of rows
NumRows = ceil(NumQuest/NumCols);

% map edit box numbers to a matrix
% to factor in the position in the columns
% that are blank
QuestInd = zeros(NumRows,NumCols);
questvals = 1:NumQuest;
NumBlanks = numel(QuestInd) - NumQuest;
for n=1:NumCols
    if n<=(NumCols-NumBlanks)
        QuestInd(:,n) = questvals(1:NumRows)';
        questvals(1:NumRows) = [];
    else
        QuestInd(1:end-1,n) = questvals(1:NumRows-1)';
        questvals(1:NumRows-1) = [];
    end
end

% Get the max height for questions and 
% edit boxes per row
tempHeight = zeros(NumRows,NumCols);
tempHeight(QuestInd>0) = EditHeight(:);
EditHeight = tempHeight;
tempHeight(QuestInd>0) = QuestHeight(:);
QuestHeight = tempHeight;
if NumCols == 1
    maxEditHeight = EditHeight';
    maxQuestHeight = QuestHeight';
else
    maxEditHeight = max(EditHeight');
    maxQuestHeight = max(QuestHeight');
end
for n=1:NumRows 
    EditHeight(n,:) = maxEditHeight(n);
    QuestHeight(n,:) = maxQuestHeight(n);
end

% Store figure height
if NumRows == 1
    FigHeight=(NumRows+2)*DefOffset    + ...
        BtnHeight + max(EditHeight) + ...
    max(QuestHeight);
else
    
    FigHeight=(NumRows+2)*DefOffset    + ...
        BtnHeight + max(sum(EditHeight)) + ...
    max(sum(QuestHeight));
end
%*******************************************
TxtXOffset=DefOffset;

%********************** EDITTED ************
% Default text width is set to fit the figure width
TxtWidth=FigWidth/NumCols-2*DefOffset;
%*******************************************

QuestYOffset=zeros(NumQuest,1);
EditYOffset=zeros(NumQuest,1);
QuestYOffset(1)=FigHeight-DefOffset-QuestHeight(1);
EditYOffset(1)=QuestYOffset(1)-EditHeight(1);% -SmallOffset;

for YOffLp=2:NumQuest
  QuestYOffset(YOffLp)=EditYOffset(YOffLp-1)-QuestHeight(YOffLp)-DefOffset;
  EditYOffset(YOffLp)=QuestYOffset(YOffLp)-EditHeight(YOffLp); %-SmallOffset;
end % for YOffLp

QuestHandle=[];
EditHandle=[];
FigWidth =1;

AxesHandle=axes('Parent',InputFig,'Position',[0 0 1 1],'Visible','off');

%********************** EDITTED ************
% Set mulitiple columns boxes in inputdlg
column_flag = 1;
shape_rows = [];
NumQuest_remaining = NumQuest;
num_col_left = NumCols;
oldlp = 1;
%*******************************************

for lp=1:NumQuest
    QuestTag=['Prompt' num2str(lp)];
    EditTag=['Edit' num2str(lp)];
    if ~ischar(DefAns{lp})
        delete(InputFig);
        error('Default answers must be strings in INPUTDLG.');
    end
    
    %********************** EDITTED ************
    % Move edit box to start a new column
    
    if num_col_left ~= 0
        if (lp-oldlp) >= (NumQuest_remaining/num_col_left) 
            if column_flag == 1
                shape_rows = (lp+1)-oldlp;
                column_flag = 2;
            end
            pos_edit = get(EditHandle(1),'Position');
            TxtXOffset = TxtXOffset + pos_edit(3)+ DefOffset;
            EditYOffset(lp:(lp+(shape_rows-1))) = EditYOffset(1:shape_rows);
            QuestYOffset(lp:(lp+(shape_rows-1))) = QuestYOffset(1:shape_rows);
            NumQuest_remaining = NumQuest-round(lp-1);
            num_col_left = num_col_left - 1;
            oldlp = lp;
        end
    end
    %*******************************************
    
    QuestHandle(lp)=text('Parent',AxesHandle, ...
        TextInfo     , ...
        'Position'   ,[ TxtXOffset QuestYOffset(lp)], ...
        'String'     ,WrapQuest{lp}                 , ...
        'Color'      ,TextForeground                , ...
        'Interpreter',Interpreter                   , ...
        'Tag'        ,QuestTag                        ...
    );
    
    EditHandle(lp)=uicontrol(InputFig   ,EdInfo     , ...
        'Max'       ,NumLines(lp,1)       , ...
        'Position'  ,[ TxtXOffset EditYOffset(lp) ...
            TxtWidth   EditHeight(lp)  ...
    ]                    , ...
        'String'    ,DefAns{lp}           , ...
        'Tag'       ,QuestTag               ...
    );               
    if size(NumLines,2) == 2
        set(EditHandle(lp),'String',char(ones(1,NumLines(lp,2))*'x'));
        Extent = get(EditHandle(lp),'Extent');
        NewPos = [TxtXOffset EditYOffset(lp)  Extent(3) EditHeight(lp) ];
        
        NewPos1= [TxtXOffset QuestYOffset(lp)];
        set(EditHandle(lp),'Position',NewPos,'String',DefAns{lp})
        set(QuestHandle(lp),'Position',NewPos1)
        
        FigWidth=max(FigWidth,Extent(3)+2*DefOffset);
    else
        FigWidth=max(175,TxtWidth+2*DefOffset);
    end
    
end % for lp
set(InputFig ,'Visible','on')
FigPos=get(InputFig,'Position');

Temp=get(0,'Units');
set(0,'Units','points');
ScreenSize=get(0,'ScreenSize');
set(0,'Units',Temp);

%********************** EDITTED ************
% Set size of the figure window width
pos_edit = get(EditHandle(end),'Position');
FigWidth = DefOffset+pos_edit(1)+pos_edit(3);
%*******************************************

FigWidth=max(FigWidth,2*(BtnWidth+DefOffset)+DefOffset);
FigPos(1)=(ScreenSize(3)-FigWidth)/2;
FigPos(2)=(ScreenSize(4)-FigHeight)/2;
FigPos(3)=FigWidth;
FigPos(4)=FigHeight;

set(InputFig,'Position',FigPos);

CBString='set(gcbf,''UserData'',''Cancel'');uiresume';

CancelHandle=uicontrol(InputFig   ,              ...
                      BtnInfo     , ...
                      'Position'  ,[FigWidth-BtnWidth-DefOffset DefOffset ...
                                    BtnWidth  BtnHeight  ...
                                   ]           , ...
                      'String'    ,'Cancel'    , ...
                      'Callback'  ,CBString    , ...
                      'Tag'       ,'Cancel'      ...
                      );
                                   
                                   
CBString='set(gcbf,''UserData'',''OK'');uiresume';

OKHandle=uicontrol(InputFig    ,              ...
                   BtnInfo     , ...
                   'Position'  ,[ FigWidth-2*BtnWidth-2*DefOffset DefOffset ...
                                  BtnWidth                    BtnHeight ...
                                ]           , ...
                  'String'     ,'OK'        , ...
                  'Callback'   ,CBString    , ...
                  'Tag'        ,'OK'          ...
                  );
    
Data.OKHandle = OKHandle;
Data.CancelHandle = CancelHandle;
Data.EditHandles = EditHandle;
Data.QuestHandles = QuestHandle;
Data.LineInfo = NumLines;
Data.ButtonWidth = BtnWidth;
Data.ButtonHeight = BtnHeight;
Data.EditHeight = TxtHeight+4;
Data.Offset = DefOffset;
set(InputFig ,'Visible','on','UserData',Data);
% This drawnow is a hack to work around a bug
drawnow
h_temp = findall(InputFig); 
set(h_temp(isprop(h_temp,'Units')),'Units','normalized');
set(h_temp,'HandleVisibility','callback');
set(InputFig,'Units','points')
try
    uiwait(InputFig);
catch
    delete(InputFig);
end

TempHide=get(0,'ShowHiddenHandles');
set(0,'ShowHiddenHandles','on');

if any(get(0,'Children')==InputFig)
  Answer={};
  if strcmp(get(InputFig,'UserData'),'OK')
    Answer=cell(NumQuest,1);
    for lp=1:NumQuest
      Answer(lp)=get(EditHandle(lp),{'String'});
    end % for
  end % if strcmp
  delete(InputFig);
else
  Answer={};
end % if any

set(0,'ShowHiddenHandles',TempHide);


function LocalResizeFcn(FigHandle)
  Data=get(FigHandle,'UserData');
  
  %Data.ButtonHandles = [ OKHandles CancelHandle];
  %Data.EditHandles = EditHandle;
  %Data.QuestHandles = QuestHandle;
  %Data.LineInfo = NumLines;
  %Data.ButtonWidth = BtnWidth;
  %Data.ButtonHeight = BtnHeight;
  %Data.EditHeight = TxtHeight;
  
  set(findall(FigHandle),'Units','points');
  
  FigPos = get(FigHandle,'Position');
  FigWidth = FigPos(3); FigHeight = FigPos(4);
  
  OKPos = [ FigWidth-Data.ButtonWidth-Data.Offset Data.Offset ...
	    Data.ButtonWidth                      Data.ButtonHeight ];
  CancelPos =[Data.Offset Data.Offset Data.ButtonWidth  Data.ButtonHeight];
  set(Data.OKHandle,'Position',OKPos);
  set(Data.CancelHandle,'Position',CancelPos);

  % Determine the height of all question fields
  YPos = sum(OKPos(1,[2 4]))+Data.Offset;
  QuestPos = get(Data.QuestHandles,{'Extent'});
  QuestPos = cat(1,QuestPos{:});
  QuestPos(:,1) = Data.Offset;
  RemainingFigHeight = FigHeight - YPos - sum(QuestPos(:,4)) - ...
                       Data.Offset - size(Data.LineInfo,1)*Data.Offset;
  
  Num1Liners = length(find(Data.LineInfo(:,1)==1));
  
  RemainingFigHeight = RemainingFigHeight - ...
      Num1Liners*Data.EditHeight;
  
  Not1Liners = find(Data.LineInfo(:,1)~=1);

  %Scale the 1 liner heights appropriately with remaining fig height
  TotalLines = sum(Data.LineInfo(Not1Liners,1));
  
  % Loop over each quest/text pair
  
  for lp = 1:length(Data.QuestHandles)
   CurPos = get(Data.EditHandles(lp),'Position');
   NewPos = [Data.Offset YPos  CurPos(3) Data.EditHeight ];
   if Data.LineInfo(lp,1) ~= 1
     NewPos(4) = RemainingFigHeight*Data.NumLines(lp,1)/TotalLines;
   end
    
   set(Data.EditHandles(lp),'Position',NewPos)
   YPos = sum(NewPos(1,[2 4]));
   QuestPos(lp,2) = YPos;QuestPos(lp,3) = NewPos(3);
   set(Data.QuestHandles(lp),'Position',QuestPos(lp,:));
   YPos = sum(QuestPos(lp,[2 4]))+Data.Offset;
 end
 
 if YPos>FigHeight - Data.Offset
   FigHeight = YPos+Data.Offset;
   FigPos(4)=FigHeight;
   set(FigHandle,'Position',FigPos);  
   drawnow
 end
 set(FigHandle,'ResizeFcn','inputdlg InputDlgResizeCB');
 
 set(findall(FigHandle),'Units','normalized')
 
```
