## Supplementary material for "AMPK Regulates Phagophore-to-Autophagosome Maturation": K-FOCUS Code: IntensityCorrectionParallel.m

```
%% SpotAnalysis script by Carlo Barnaba 07202022 %%

function [batch] = IntensityCorrectionParallel(batch)

intensity_max_nocorrection_tot = [];
intensity_max_raw_tot = [];
intensity_max_max_tot = [];

% Get the Max intensity for all the tracks from TrackIT
parfor i = 1:length(batch)
    tracks = batch(i).results.tracks;
    intensity_max_nocorrection = [];
    
    for j = 1:length(tracks)
        intensity_max_nocorrection = [intensity_max_nocorrection; mean(tracks{1, j}(:, 5))];
    end
    
    intensity_max_nocorrection_tot = [intensity_max_nocorrection_tot; intensity_max_nocorrection];
end

% Background correction for the tracks
parfor iter = 1:length(batch)
    tracks = batch(iter).results.tracks;

    % Specify tif file
    filename = strcat(batch(iter).movieInfo.pathName, batch(iter).movieInfo.fileName);

    % Extract the number of frames in the TIFF file
    num_frames = length(imfinfo(filename));

    % Pre-allocate an empty cell array to store the frames
    image = cell(num_frames, 1);

    % Read the TIFF file using the built-in MATLAB function "Tiff"
    for i = 1:num_frames
        image{i} = flip(imread(filename, i));
    end

    for i = 1:length(tracks)
        x_coord = tracks{1, i}(:, 2); % olympus BT-fusion (2X2 bin, 6.5 um, 60X objective)
        y_coord = tracks{1, i}(:, 3); % olympus BT-fusion
        max_track = [];
        intensity_max_raw = [];
        intensity_max = [];
        intensity_max_max = [];
        background = [];
        tempbackground = [];
        
        for j = 1:length(x_coord)
            frame_num = tracks{1, i}(j, 1);
            frame = image{frame_num};
            xCenter = round(x_coord(j));
            yCenter = round(y_coord(j));
            
            if xCenter >= 6 && xCenter <= 506 && yCenter >= 6 && yCenter <= 506
                sub_region1 = frame(xCenter - 5 : xCenter + 5, yCenter - 5 : yCenter + 5); % 122 pixels
                center = frame(xCenter - 3 : xCenter + 3, yCenter - 3 : yCenter + 3);
                sub_region1(3 : 9, 3 : 9) = 0; % 25 pixels
                maxi = max(center, [], 'all');
                max_track = vertcat(max_track, maxi);
                tempbackground = sum(sub_region1, 'all') / 86; % average background
                background = vertcat(background, tempbackground);
            else
                maxi = NaN;
                max_track = vertcat(max_track, maxi);
                tempbackground = NaN;
                background = vertcat(background, tempbackground);
            end
        end
        
        batch(iter).results.tracks{1, i}(:, 10) = max_track;
        batch(iter).results.tracks{1, i}(:, 11) = background;
        batch(iter).results.tracks{1, i}(:, 12) = double(max_track) - background;
    end
end

end
```
