## Supplementary material for "AMPK Regulates Phagophore-to-Autophagosome Maturation": K-FOCUS Code: markerdistance.m

```
function D = markerdistance(A, varargin)
% `MARKERDISTANCE` Pairwise distance between two marker sets.
% This function behaves like the built-in 'pdist2', but the code was vectorized and extended to nD-array (i.g., i-by-j-by-k).
% The efficiency of this function is about 10x~100x times compared to `pdist2`.
%
%   D = markerdistance(A, Q) returns the Euclidean distance between each pair of markers in A and Q.
%   D = markerdistance(A) is equivalent to D = markerdistance(A, A), which returns the Euclidean distance between pairs of markers in A.
%   Columns of the marker set correspond to dimensions (e.g., 3)
%   D is an MX-by-MY-by-NSlice matrix, with the (I,J, P) entry equal to the distance between marker I in A set and marker J in Q set at P-th time frame.
%
%   D = markerdistance(A, Q, metric) computes D using specified distance measure.
%   Currently avaliable distance measures are `euclidean`,
%   `cityblock`,`chebychev`, `minkowski`, `cosine`, `correlation`,
%   'mahalanobis', 'chisquare', 'emd'
%
% |      Measure       | Generalized vector p-norm |                                                                     Syntax                                                                     |                                                                           Description                                                                            |
% | :-----------------: | :----------------------------------: | :------------------------------------------------------------------------------------------------: | :--------------------------------------------------------------------------------------------------------------: |
% |    'euclidean'    |                        p = 2                       |                `markerdistance(_, 2)`, `markerdistance(_, 'euclidean')`                  |                                                          Euclidean distance (default)                                                           |
% |    'cityblock'     |                        p = 0                        |                 `markerdistance(_, 0)`, `markerdistance(_, 'cityblock')`                  |                                                      City block (Manhattan) distance                                                       |
% |   'chebychev'   |                         p = inf                    |              `markerdistance(_, inf)`, `markerdistance(_, 'chebychev')`               |                               Chebychev distance (maximum coordinate difference)                               |
% |   'minkowski'   |           p = positive scalar         |                                                    `markerdistance(_, p)`                                                     |                                                                   Minkowski distance                                                                   |
% |       'cosine'        |                         NA                          |                                              `markerdistance(_, 'cosine')`                                              |      One minus the cosine of the included angle between points (treated as vectors)     |
% |   'correlation'   |                         NA                         |                                         `markerdistance(_, 'correlation')`                                          |  One minus the sample correlation between points (treated as sequences of values) |
% |    'chisquare'    |                         NA                          |               `markerdistance(_, 'chisquare')`, `markerdistance(_, 'x2')`               |                      The chi-squared distance is useful when comparing histograms.                     |
% |          'emd'         |                         NA                          |                 `markerdistance(_, 'emd')`, `markerdistance(_, 'earth')`                 |                    Earth Mover's Distance between positive 1D vectors (histograms).                    |
%
%   For other options of distance measures, use the built-in 'pdist2'.
%
% Note.
%   This function heavly depends on the calculation of vector-wise norm.
%   If you don't have the builtin `vecnorm` that was introduced in R2017b,
%   use local subfunction `ndnorm` as an alternative (all you need to do is simply replace `vecnorm` with `ndnorm`).
%
% I don't really know much about the measurements above as I am not working at these research fields.
% I just found some useful pieces of code in FEX or builtin 'pdist2', and realized that they can be vectorized.
% All I did is vectorize the codes I found, and then ensured the results were identical to the original codes.
% Don't blame on me if the results are not what you expected. It might not be my problems.
% 
% 
% See also pdist2, pdist, norm, vecnorm.
%
% ---------------------------------------------------------------------------------------------------------------------------
%
% Created by Wen-Feng Huang, OTMSATLab, Dec, 2022.
%   TODO: completely implement the Mahalanobis distance
% Revised by Wen-Feng Huang, OTMSATLab, Dec, 2022.
%   feat(measure): support Mahalanobis distance, Chi-squared distance, and Earth Mover's distance.
%   doc(help): typo & table of measures.
% ---------------------------------------------------------------------------------------------------------------------------

NIs = nargin;

if NIs == 1 % markerdistance(X)
    D = fn_wrapper(A, A, 2);

elseif NIs == 2 % markerdistance(X, p), markerdistance(X, Y)

    if (isscalar(varargin{1}) && (isnumeric(varargin{1}) || isstring(varargin{1}))) || ischar(varargin{1})
        D = fn_wrapper(A, A, varargin{:});

    else
        D = fn_wrapper(A, varargin{:}, 2);
    end

elseif NIs == 3 % markerdistance(X, Y, p)
    D = fn_wrapper(A, varargin{:});
end
end

function D = coredist(A, Q, sz, pn)

% it might have memory issues

xrow = 1:sz(1);
ycol = 1:sz(2);
xi = repmat(xrow(:), [1, sz(2)]);
yi = repmat(ycol, [sz(1), 1]);
X = A(xi(:), :, :);
Y = Q(yi(:), :, :);

D = reshape(vecnorm(X-Y, pn, 2), sz(1), sz(2), []);
% If you don't have the `vecnorm`, use `ndnorm` instead.
% D = reshape(ndnorm(X-Y, pn, 2), sz(1), sz(2), []);

%{
% special case: pn == 2
if size(D, 3) == 1
    % D = sqrt(dot(Q, Q, 2).'+dot(A, A, 2)-2*A*Q.');
else
    % If you don't have the `pagemtimes`, use `mtimesx` instead.
    % If you don't have the `pagetranspose`, use `permute` instead.
    % D = sqrt((pagetranspose(dot(Q, Q, 2))+dot(A, A, 2))-2*pagemtimes(A, 'n', Q, 't'));
end
%}
end

function D = specialdist(A, Q, sz, pn)
xrow = 1:sz(1);
ycol = 1:sz(2);
xi = repmat(xrow(:), [1, sz(2)]);
yi = repmat(ycol, [sz(1), 1]);
xi = xi(:);
yi = yi(:);

if ~isscalar(pn)
    CX = pn;
    pn = -6;
end

% I don't really know much about the measurements below as I am not working at these research fields.
% I just found some pieces of code in FEX or builtin 'pdist2', and awared that they can be vectorized.
% All I did is vectorize the codes I found, and then ensured the results were identical to the original codes.
% Don't blame on me if the result is not as you expected. I might not be my problems.
switch pn

    case -4 % cosine
        X = A(xi, :, :);
        Y = Q(yi, :, :);
        X = X./vecnorm(X, 2, 2);
        Y = Y./vecnorm(Y, 2, 2);
        D = 1-reshape(sum(X.*Y, 2), sz(1), sz(2), []);
        % D = 1-reshape(dot(X, Y, 2), sz(1), sz(2), []);
        D(D<0) = 0;

    case -5 % correlation
        X0 = A-mean(A, 2);
        Q0 = Q-mean(Q, 2);
        X = X0(xi, :, :);
        Y = Q0(yi, :, :);
        X = X./vecnorm(X, 2, 2);
        Y = Y./vecnorm(Y, 2, 2);
        D = 1-reshape(sum(X.*Y, 2), sz(1), sz(2), []);
        % D = 1-reshape(dot(X, Y, 2), sz(1), sz(2), []);
        D(D<0) = 0;

    case -6 % mahalanobis
        % Obtain Mahalanobis distance from Q to the center of reference samples A.
        % The following pieces of code are equivalent.
        % `sqrt(mahal(Q, A))`
        % `pdist2(Q, mean(A), 'mah', cov(A))`
        % `markerdistance(Q, mean(A), cov(A))`
        % `for ii = 1:size(Q, 2), cv.Mahalanobis(Q(ii, :), mean(A), inv(cov(A))), end`

        if ~exist('CX', 'var')
            mu = mean(A, 1);
            A0 = A-mu;
            CX = pagemtimes(A0, 't', A0, 'n')./(sz(1)-1); % cov(A)
        end
        ci = pageinv(CX);
        npages = max(size(A, 3), size(Q, 3));

        %         % This seems to be slower as it needs more runs in a for-loop.
        %         X = A(xi, :, :);
        %         Y = Q(yi, :, :);
        %         N = sz(1)*sz(2);
        %         delta = X-Y;
        %         tmpD = zeros(N, npages);
        %         for ii = 1:sz(1)*sz(2)
        %             cdelta = delta(ii, :, :);
        %             tmpD(ii, :) = pagemtimes(pagemtimes(cdelta, ci), 'n', cdelta, 't');
        %         end
        %         D = sqrt(reshape(tmpD, sz(1), sz(2), []));

        tmpD = zeros(sz(1), sz(2), npages);
        pci = permute(ci, [1, 2, 4, 3]); % expand dimension

        if sz(2) > 2 % prevent from too-many permutations
            X = A(xi, :, :);
            Y = Q(yi, :, :);
            delta = X-Y;
            delta = permute(reshape(permute(delta, [2, 1, 3]), [], sz(1), sz(2), npages), [1, 3, 2, 4]);

            for p = 1:sz(2)
                cdelta = delta(:, p, :, :);
                tmpD(:, p, :) = pagemtimes(pagemtimes(cdelta, 't', pci, 'n'), cdelta);
                % tmpD(:, p, :) = reshape(pagemtimes(pagemtimes(cdelta, 't', pci, 'n'), cdelta), sz(1), 1, []);
            end

        else

            for p = 1:sz(2)
                cdelta = permute( A-Q(p, :, :), [2, 4, 1, 3]);
                tmpD(:, p, :) = pagemtimes(pagemtimes(cdelta, 't', pci, 'n'), cdelta);
                % tmpD(:, p, :) = reshape(pagemtimes(pagemtimes(cdelta, 't', pci, 'n'), cdelta), sz(1), 1, []);
            end
        end

        D = sqrt(tmpD);

    case -7 % chi-squared % `distChiSq` in Piotr's Image & Video Matlab Toolbox
        X = A(xi, :, :);
        Y = Q(yi, :, :);
        D = sum(((Y-X).^2)./(Y+X), 2)/2;
        D = reshape(D, sz(1), sz(2), []);

    case -8 % chi-squared % `distEmd` in Piotr's Image & Video Matlab Toolbox
        Xcdf = cumsum(A, 2);
        Ycdf = cumsum(Q,2);
        X = Xcdf(xi, :, :);
        Y = Ycdf(yi, :, :);
        D = reshape(sum(abs(X - Y),2), sz(1), sz(2), []);
end
end

function D = fn_wrapper(A, Q, pn)
szA = size(A);
szQ = size(Q);

if (szA(2) == szQ(2))
    % in case A or Q is a 2D array.
    szA(3) = size(A, 3);
    szQ(3) = size(Q, 3);

    if szA(3) == szQ(3) || szA(3) == 1 || szQ(3) == 1

        if ~isnumeric(pn)
            pn = regexprep(pn, '[^a-zA-Z0-9]', '');
            strl = length(pn);
            strl = min(strl, 3);
            ix = strncmpi(pn, ...
                {'Cityblock', 'Manhattan', 'L1';...
                'Euclidean', '', 'L2'; ...
                'Chebychev', 'chessboard', 'inf'; ...
                'cosine', 'cosine', 'cosine'; ...
                'correlations', 'correlations', 'correlations'; ...
                'mahalanobis', 'mahalanobis', 'mahalanobis'; ...
                'chisquared', 'chi2', 'x2'; ...
                'EarthMover', 'emd', 'emd'; ...
                }, ...
                strl);

            if any(ix(:))
                pn = find(any(ix, 2));
                pn(pn > 3) = -pn(pn > 3);
                pn(pn == 3) = inf;

            else
                warning('wfH:ChkAIO:Val:UnrecognizedDistance', 'Unrecognized Distance measure: "%s". Compute the Euclidean distance by default.', pn);
                pn = 2;
            end

            if pn > 0
                D = coredist(A, Q, [szA(1), szQ(1)], pn);

            else
                D = specialdist(A, Q, [szA(1), szQ(1)], pn);
            end

        else

            if isscalar(pn)
                D = coredist(A, Q, [szA(1), szQ(1)], pn);

            else % Mahalanobis
                % given covariance of A
                if all(abs(pn-eye(szA(2))) <= eps, "all")
                    % Degenerate to Eucliden
                    D = coredist(A, Q, [szA(1), szQ(1)], 2);
                    
                else
                    D = specialdist(A, Q, [szA(1), szQ(1)], pn);
                end
            end
        end

    else
        error('wfH:ChkAIO:Size', 'X and Y must have the same number of slice.');
    end

else
    error('wfH:ChkAIO:Size', 'X and Y must have the same number of columns.');
end
end

function B = ndnorm(A, varargin) %#ok<DEFNU>
% -------------------------------------------------------------------------
% NDNORM calculates norm over specified dimension of an array.
%   For vector, B is the magnitude of A (built-in function "NORM" is recommended for one-by-n vector).
%   For matrices, B is a vector listing the  norm of the vectors constructed along the specified dimension.
%
% Syntax:
%       B = NDNORM(A)
%       B = NDNORM(A, p)
%       B = NDNORM(A, p, DIM)
%   The first non-singleton dimension is used if DIM is not specified.
%
% See also NORM, VECNORM, DOT.
% -------------------------------------------------------------------------
NIs = nargin;

if NIs < 2 % Euclidian norm over specified dimension
    % B = vecnorm(A);
    B = sqrt(sum(conj(A).*A));

else % generalized vector p-norm over specified dimension
    % B = vecnorm(A, varargin{:});
    B = sum(abs(A).^varargin{1}, varargin{2}).^(1/varargin{1});
end
end
```
