## Supplementary material for "AMPK Regulates Phagophore-to-Autophagosome Maturation": K-FOCUS Code: sqdist.m

```
function D = sqdist(X1, X2)
% Pairwise square Euclidean distance between two sample sets
% Input:
%   X1, X2: dxn1 dxn2 sample matrices
% Output:
%   D: n1 x n2 square Euclidean distance matrix
% Written by Mo Chen.
% X1 = track(1,1);
% X2 = track(2,1);
D = bsxfun(@plus,dot(X2,X2,1),dot(X1,X1,1)')-2*(X1'*X2);
```
