## Supplementary material for "AMPK Regulates Phagophore-to-Autophagosome Maturation": K-FOCUS Code: stepsize.m

```
function [steps] = stepsize(track)
    steps = zeros(length(track)-1, 1); % preallocate steps vector
    for a = 1:length(track)-1 % loop through all points of the track except the last
        loc = track(a,1:2); % current particle location
        locplusone = track(a+1,1:2); % next location
        steps(a) = markerdistance(loc, locplusone); % determine stepsize and store in steps vector
    end
end
```
