## Supplementary material for "AMPK Regulates Phagophore-to-Autophagosome Maturation": K-FOCUS Code: uipickfiles.m

```
function out = uipickfiles(varargin)
%uipickfiles: GUI program to select files and/or folders.
%
% Syntax:
%   files = uipickfiles('PropertyName',PropertyValue,...)
%
% The current folder can be changed by operating in the file navigator:
% double-clicking on a folder in the list or pressing Enter to move further
% down the tree, using the popup menu, clicking the up arrow button or
% pressing Backspace to move up the tree, or typing a path in the box to
% move to any folder.  Right-clicking on the path box (control-click on the
% Mac) will pop up a context menu listing previously-visited folders. These
% folders are listed in order of when they were last visited (most recent
% at the top) and the list is saved between calls to uipickfiles.  The list
% can be cleared or its maximum length changed with the items at the bottom
% of the menu. Also included (and unclearable) are the user's home folder
% and one or more MATLAB startup folders.
%
% (Windows only: To go to a UNC-named resource you will have to type the
% UNC name in the path box, but all such visited resources will be
% remembered and listed along with the mapped drives.)  The items in the
% file navigator can be sorted by name, modification date or size by
% clicking on the headers, though neither date nor size are displayed.  All
% folders are considered to have zero size.
%
% Files can be added to the list by double-clicking or selecting files
% (non-contiguous selections are possible with the control key) and
% pressing the Add button.  Control-F will select all the files listed in
% the navigator while Control-A will select everything (Command instead of
% Control on the Mac).  Since double-clicking a folder will open it,
% folders can be added only by selecting them and pressing the Add button.
% Files/folders in the list can be removed or re-ordered.  Recall button
% will insert into the Selected Files list whatever files were returned the
% last time uipickfiles was run.  When finished, a press of the Done button
% will return the full paths to the selected items in a cell array,
% structure array or character array.  If the Cancel button or the escape
% key is pressed then zero is returned.
%
% The figure can be moved and resized in the usual way and this position is
% saved and used for subsequent calls to uipickfiles.  The default position
% can be restored by double-clicking in a vacant region of the figure.
%
% The following optional property/value pairs can be specified as arguments
% to control the indicated behavior:
%
%   Property    Value
%   ----------  ----------------------------------------------------------
%   FilterSpec  String to specify starting folder and/or file filter.
%               Ex:  'C:\bin' will start up in that folder.  '*.txt'
%               will list only files ending in '.txt'.  'c:\bin\*.txt' will
%               do both.  Default is to start up in the current folder and
%               list all files.  Can be changed with the GUI.
%
%   REFilter    String containing a regular expression used to filter the
%               file list.  Ex: '\.m$|\.mat$' will list files ending in
%               '.m' and '.mat'.  Default is empty string.  Can be used
%               with FilterSpec and both filters are applied.  Can be
%               changed with the GUI.
%
%   REDirs      Logical flag indicating whether to apply the regular
%               expression filter to folder names.  Default is false which
%               means that all folders are listed.  Can be changed with the
%               GUI.
%
%   Type        Two-column cell array where the first column contains file
%               filters and the second column contains descriptions.  If
%               this property is specified an additional popup menu will
%               appear below the File Filter and selecting an item will put
%               that item into the File Filter.  By default, the first item
%               will be entered into the File Filter.  For example,
%                   { '*.m',   'M-files'   ;
%                     '*.mat', 'MAT-files' }.
%               Can also be a cell vector of file filter strings in which
%               case the descriptions will be the same as the file filters
%               themselves.
%               Must be a cell array even if there is only one entry.
%
%   Prompt      String containing a prompt appearing in the title bar of
%               the figure.  Default is 'Select files'.
%
%   NumFiles    Scalar or vector specifying number of files that must be
%               selected.  A scalar specifies an exact value; a two-element
%               vector can be used to specify a range, [min max].  The
%               function will not return unless the specified number of
%               files have been chosen.  Default is [] which accepts any
%               number of files.
%
%   Append      Cell array of strings, structure array or char array
%               containing a previously returned output from uipickfiles.
%               Used to start up program with some entries in the Selected
%               Files list.  Any included files that no longer exist will
%               not appear.  Default is empty cell array, {}.
%
%   Output      String specifying the data type of the output: 'cell',
%               'struct' or 'char'.  Specifying 'cell' produces a cell
%               array of strings, the strings containing the full paths of
%               the chosen files.  'Struct' returns a structure array like
%               the result of the dir function except that the 'name' field
%               contains a full path instead of just the file name.  'Char'
%               returns a character array of the full paths.  This is most
%               useful when you have just one file and want it in a string
%               instead of a cell array containing just one string.  The
%               default is 'cell'.
%
% All properties and values are case-insensitive and need only be
% unambiguous.  For example,
%
%   files = uipickfiles('num',1,'out','ch')
%
% is valid usage.
% Version: 1.22, 15 June 2020
% Author:  Douglas M. Schwarz
% Email:   dmschwarz=ieee*org, dmschwarz=urgrad*rochester*edu
% Real_email = regexprep(Email,{'=','*'},{'@','.'})
% Define properties and set default values.
prop.filterspec = '*';
prop.refilter = '';
prop.redirs = false;
prop.type = {};
prop.prompt = 'Select files';
prop.numfiles = [];
prop.append = [];
prop.output = 'cell';
% Process inputs and set prop fields.
prop = parsepropval(prop,varargin{:});
% Validate FilterSpec property.
if isempty(prop.filterspec)
	prop.filterspec = '*';
end
if ~ischar(prop.filterspec)
	error('FilterSpec property must contain a string.')
end
% Validate REFilter property.
if ~ischar(prop.refilter)
	error('REFilter property must contain a string.')
end
% Validate REDirs property.
if ~isscalar(prop.redirs)
	error('REDirs property must contain a scalar.')
end
% Validate Type property.
if isempty(prop.type)
elseif iscellstr(prop.type) && isvector(prop.type)
	prop.type = repmat(prop.type(:),1,2);
elseif iscellstr(prop.type) && size(prop.type,2) == 2
else
	error(['Type property must be empty or a cellstr vector or ',...
		'a 2-column cellstr matrix.'])
end
% Validate Prompt property.
if ~ischar(prop.prompt)
	error('Prompt property must contain a string.')
end
% Validate NumFiles property.
if numel(prop.numfiles) > 2 || any(prop.numfiles < 0)
	error('NumFiles must be empty, a scalar or two-element vector.')
end
prop.numfiles = unique(prop.numfiles);
if isequal(prop.numfiles,1)
	numstr = 'Select exactly 1 item.';
elseif length(prop.numfiles) == 1
	numstr = sprintf('Select exactly %d items.',prop.numfiles);
else
	numstr = sprintf('Select %d to %d items.',prop.numfiles);
end
% Validate Append property and initialize pick data.
if isstruct(prop.append) && isfield(prop.append,'name')
	prop.append = {prop.append.name};
elseif ischar(prop.append)
	prop.append = cellstr(prop.append);
end
if isempty(prop.append)
	file_picks = {};
	full_file_picks = {};
	dir_picks = repmat(dir(char(127)),0,1);  % Create empty directory structure.
elseif iscellstr(prop.append) && isvector(prop.append)
	num_items = length(prop.append);
	file_picks = cell(1,num_items);
	full_file_picks = cell(1,num_items);
	dir_fn = fieldnames(repmat(dir(char(127)),0,1));
	dir_picks = repmat(cell2struct(cell(length(dir_fn),1),dir_fn(:)),...
		num_items,1);
	for item = 1:num_items
		if fdexist(prop.append{item},'dir') && ...
				~any(strcmp(full_file_picks,prop.append{item}))
			full_file_picks{item} = prop.append{item};
			[unused,fn,ext] = fileparts(prop.append{item});
			file_picks{item} = [fn,ext];
			path_c_tmp = path2cell(prop.append{item});
			temp = dir(cell2path(path_c_tmp(1:end-1)));
			if ispc || ismac
				thisdir = strcmpi({temp.name},[fn,ext]);
			else
				thisdir = strcmp({temp.name},[fn,ext]);
			end
			dir_picks(item) = temp(thisdir);
			dir_picks(item).name = prop.append{item};
		elseif fdexist(prop.append{item},'file') && ...
				~any(strcmp(full_file_picks,prop.append{item}))
			full_file_picks{item} = prop.append{item};
			[unused,fn,ext] = fileparts(prop.append{item});
			file_picks{item} = [fn,ext];
			dir_picks(item) = dir(prop.append{item});
			dir_picks(item).name = prop.append{item};
		else
			continue
		end
	end
	% Remove items which no longer exist.
	missing = cellfun(@isempty,full_file_picks);
	full_file_picks(missing) = [];
	file_picks(missing) = [];
	dir_picks(missing) = [];
else
	error('Append must be a cell, struct or char array.')
end
% Validate Output property.
legal_outputs = {'cell','struct','char'};
out_idx = find(strncmpi(prop.output,legal_outputs,length(prop.output)));
if length(out_idx) == 1
	prop.output = legal_outputs{out_idx};
else
	error(['Value of ''Output'' property, ''%s'', is illegal or '...
		'ambiguous.'],prop.output)
end
% Set style preference for display of folders.
%   1 => folder icon before and filesep after
%   2 => bullet before and filesep after
%   3 => filesep after only
folder_style_pref = 1;
fsdata = set_folder_style(folder_style_pref);
% Set style preference for context menu.
%   1 => home and logo icons before home and MATLAB folders
%   2 => no icons
cmenu_style_pref = 1;
csdata = set_cmenu_style(cmenu_style_pref);
% Get canonical names of current directory and filter.
jUserDir = java.lang.System.setProperty('user.dir',pwd);
if any(prop.filterspec == '*')
	jFile = java.io.File(prop.filterspec).getAbsoluteFile;
	current_dir  = char(jFile.getParentFile().getCanonicalPath());
	filter = char(jFile.getName());
else
	jFile = java.io.File(prop.filterspec).getCanonicalFile();
	if jFile.isDirectory()
		current_dir = char(jFile);
		filter = '*';
	else
		jDir = java.io.File(jFile.getParent());
		if jDir.isDirectory()
			current_dir = char(jDir);
			filter = char(jFile.getName());
		else
			java.lang.System.setProperty('user.dir',jUserDir);
			error('Path does not exist.')
		end
	end
end
java.lang.System.setProperty('user.dir',jUserDir);
% Initialize file lists.
re_filter = prop.refilter;
full_filter = fullfile(current_dir,filter);
network_volumes = {};
[path_cell,new_network_vol] = path2cell(current_dir);
if fdexist(new_network_vol,'dir')
	network_volumes = unique([network_volumes,{new_network_vol}]);
end
fdir = filtered_dir(full_filter,re_filter,prop.redirs,...
	@(x,c)file_sort(x,[1 0 0],c));
filenames = {fdir.name}';
[filenames,fdir] = annotate_file_names(filenames,fdir,fsdata);
% Initialize some data.
show_full_path = false;
nodupes = true;
% Get history preferences and set history.
history = getpref('uipickfiles','history',struct('name',{},'time',{}));
default_history_size = 15;
history_size = getpref('uipickfiles','history_size',default_history_size);
% Get favorites.
favorites = getpref('uipickfiles','favorites',{});
% Set history.
history = update_history(history,current_dir,now,history_size);
% Get figure position preference and create figure.
gray = [220 220 220]/255;
if ispref('uipickfiles','figure_position')
	fig_pos = getpref('uipickfiles','figure_position');
	create_fcn = '';
else
	fig_pos = [0 0 740 494];
	create_fcn = {@movegui,'center'};
end
fig = figure('Position',fig_pos,...
	'MenuBar','none',...
	'WindowStyle','modal',...
	'Color',gray,...
	'Resize','on',...
	'NumberTitle','off',...
	'Name',prop.prompt,...
	'IntegerHandle','off',...
	'CloseRequestFcn',@cancel,...
	'CreateFcn',create_fcn,...
	'ButtonDownFcn',@reset_figure_size,...
	'KeyPressFcn',@keypressmisc,...
	'Visible','off');
fig_color = fig.Color;
% Set system-dependent items.
if ismac
% 	ver = char(java.lang.System.getProperty('os.version'));
	set(fig,'DefaultUIControlFontName','Lucida Grande')
	set(fig,'DefaultUIControlFontSize',13)
	sort_ctrl_size = 12;
	mod_key = 'command';
	action = 'Control-click';
elseif ispc
% 	ver = str2double(java.lang.System.getProperty('os.version'));
	set(fig,'DefaultUIControlFontName','Segoe UI')
	set(fig,'DefaultUIControlFontSize',9)
	sort_ctrl_size = 7;
	mod_key = 'control';
	action = 'Right-click';
else
	set(fig,'DefaultUIControlFontName','Dialog')
	sort_ctrl_size = get(fig,'DefaultUIControlFontSize') - 1;
	mod_key = 'control';
	action = 'Right-click';
end
% Create uicontrols.
frame1 = uicontrol('Style','frame',...
	'BackgroundColor',fig_color,...
	'Position',[255 260 110 70]);
frame2 = uicontrol('Style','frame',...
	'BackgroundColor',fig_color,...
	'Position',[275 135 110 100]);
navlist = uicontrol('Style','listbox',...
	'Position',[10 10 250 320],...
	'String',filenames,...
	'Value',[],...
	'BackgroundColor','w',...
	'Callback',@clicknav,...
	'KeyPressFcn',@keypressnav,...
	'Max',2);
tri_up = repmat([1 1 1 1 0 1 1 1 1;1 1 1 0 0 0 1 1 1;1 1 0 0 0 0 0 1 1;...
	1 0 0 0 0 0 0 0 1],[1 1 3]);
tri_up(tri_up == 1) = NaN;
tri_down = tri_up(end:-1:1,:,:);
tri_null = NaN(4,9,3);
tri_icon = {tri_down,tri_null,tri_up};
sort_state = [1 0 0];
last_sort_state = [1 1 1];
sort_cb = zeros(1,3);
sort_cb(1) = uicontrol('Style','checkbox',...
	'Position',[15 331 70 15],...
	'String','Name',...
	'FontSize',sort_ctrl_size,...
	'BackgroundColor',fig_color,...
	'Value',sort_state(1),...
	'CData',tri_icon{sort_state(1)+2},...
	'KeyPressFcn',@keypressmisc,...
	'Callback',{@sort_type,1});
sort_cb(2) = uicontrol('Style','checkbox',...
	'Position',[85 331 70 15],...
	'String','Date',...
	'FontSize',sort_ctrl_size,...
	'BackgroundColor',fig_color,...
	'Value',sort_state(2),...
	'CData',tri_icon{sort_state(2)+2},...
	'KeyPressFcn',@keypressmisc,...
	'Callback',{@sort_type,2});
sort_cb(3) = uicontrol('Style','checkbox',...
	'Position',[155 331 70 15],...
	'String','Size',...
	'FontSize',sort_ctrl_size,...
	'BackgroundColor',fig_color,...
	'Value',sort_state(3),...
	'CData',tri_icon{sort_state(3)+2},...
	'KeyPressFcn',@keypressmisc,...
	'Callback',{@sort_type,3});
pickslist = uicontrol('Style','listbox',...
	'Position',[380 10 350 320],...
	'String',file_picks,...
	'BackgroundColor','w',...
	'Callback',@clickpicks,...
	'KeyPressFcn',@keypresslist,...
	'Max',2,...
	'Value',[]);
openbut = uicontrol('Style','pushbutton',...
	'Position',[265 295 90 30],...
	'String','Open',...
	'Enable','off',...
	'KeyPressFcn',@keypressmisc,...
	'Callback',@open);
arrow = [ ...
	'        1   ';
	'        10  ';
	'         10 ';
	'000000000000';
	'         10 ';
	'        10  ';
	'        1   '];
cmap = NaN(128,3);
cmap(double('10'),:) = [0.5 0.5 0.5;0 0 0];
arrow_im = NaN(7,76,3);
arrow_im(:,45:56,:) = ind2rgb(double(arrow),cmap);
add_cm = uicontextmenu;
add_cm_items(1) = uimenu(add_cm,...
	'Label','Add with path subfolders',...
	'Callback',{@add_with_subfolders,'pathsub'},...
	'Visible','off');
add_cm_items(2) = uimenu(add_cm,...
	'Label','Add with subfolders',...
	'Callback',{@add_with_subfolders,'sub'},...
	'Visible','off');
add_cm_items(3) = uimenu(add_cm,...
	'Label','<html>Add with <i>all</i> subfolders</html>',...
	'Separator','on',...
	'Callback',{@add_with_subfolders,'allsub'},...
	'Visible','off');
addbut = uicontrol('Style','pushbutton',...
	'Position',[265 265 90 30],...
	'String','Add    ',...
	'Enable','off',...
	'CData',arrow_im,...
	'KeyPressFcn',@keypressmisc,...
	'Callback',@add,...
	'UIContextMenu',add_cm);
removebut = uicontrol('Style','pushbutton',...
	'Position',[285 200 90 30],...
	'String','Remove',...
	'Enable','off',...
	'KeyPressFcn',@keypressmisc,...
	'Callback',@remove);
moveupbut = uicontrol('Style','pushbutton',...
	'Position',[285 170 90 30],...
	'String','Move Up',...
	'Enable','off',...
	'KeyPressFcn',@keypressmisc,...
	'Callback',@moveup);
movedownbut = uicontrol('Style','pushbutton',...
	'Position',[285 140 90 30],...
	'String','Move Down',...
	'Enable','off',...
	'KeyPressFcn',@keypressmisc,...
	'Callback',@movedown);
dir_popup = uicontrol('Style','popupmenu',...
	'Position',[10 350 225 20],...
	'BackgroundColor','w',...
	'String',path_cell,...
	'Value',length(path_cell),...
	'KeyPressFcn',@keypressmisc,...
	'Callback',@dirpopup);
uparrow = [ ...
	'  0     ';
	' 000    ';
	'00000   ';
	'  0     ';
	'  0     ';
	'  0     ';
	'  000000'];
cmap = NaN(128,3);
cmap(double('0'),:) = [0 0 0];
uparrow_im = ind2rgb(double(uparrow),cmap);
up_dir_but = uicontrol('Style','pushbutton',...
	'Position',[240 350 20 20],...
	'CData',uparrow_im,...
	'KeyPressFcn',@keypressmisc,...
	'Callback',@dir_up_one,...
	'ToolTip','Go to parent folder');
if length(path_cell) > 1
	up_dir_but.Enable = 'on';
else
	up_dir_but.Enable = 'off';
end
hist_cm = uicontextmenu;
pathbox = uicontrol('Style','edit',...
	'Position',[10 375 250 26],...
	'BackgroundColor','w',...
	'String',current_dir,...
	'HorizontalAlignment','left',...
	'TooltipString',[action,' to display folder history'],...
	'KeyPressFcn',@keypressmisc,...
	'Callback',@change_path,...
	'UIContextMenu',hist_cm);
label1 = uicontrol('Style','text',...
	'Position',[10 401 250 16],...
	'String','Current Folder',...
	'HorizontalAlignment','center',...
	'BackgroundColor',fig_color,...
	'TooltipString',[action,' to display folder history'],...
	'UIContextMenu',hist_cm);
hist_menus = [];
make_history_cm(csdata)
label2 = uicontrol('Style','text',...
	'Position',[10 440+36 80 17],...
	'String','File Filter',...
	'HorizontalAlignment','left',...
	'BackgroundColor',fig_color);
label3 = uicontrol('Style','text',...
	'Position',[100 440+36 160 17],...
	'String','Regular Expression Filter',...
	'HorizontalAlignment','left',...
	'BackgroundColor',fig_color);
label3.Position(3) = label3.Extent(3);
% 	'String','Reg. Exp. Filter',...
showallfiles = uicontrol('Style','checkbox',...
	'Position',[270 420+32 110 20],...
	'String','Show All Files',...
	'Value',0,...
	'HorizontalAlignment','left',...
	'BackgroundColor',fig_color,...
	'KeyPressFcn',@keypressmisc,...
	'Callback',@togglefilter);
showallfiles.Position(3) = showallfiles.Extent(3) + 20;
refilterdirs = uicontrol('Style','checkbox',...
	'Position',[270 420+10 100 20],...
	'String','RE Filter Dirs',...
	'Value',prop.redirs,...
	'HorizontalAlignment','left',...
	'BackgroundColor',fig_color,...
	'KeyPressFcn',@keypressmisc,...
	'Callback',@toggle_refiltdirs);
refilterdirs.Position(3) = refilterdirs.Extent(3) + 20;
filter_ed = uicontrol('Style','edit',...
	'Position',[10 420+30 80 26],...
	'BackgroundColor','w',...
	'String',filter,...
	'HorizontalAlignment','left',...
	'KeyPressFcn',@keypressmisc,...
	'Callback',@setfilspec);
refilter_ed = uicontrol('Style','edit',...
	'Position',[100 420+30 160 26],...
	'BackgroundColor','w',...
	'String',re_filter,...
	'HorizontalAlignment','left',...
	'KeyPressFcn',@keypressmisc,...
	'Callback',@setrefilter);
type_value = 1;
type_popup = uicontrol('Style','popupmenu',...
	'Position',[10 422 250 20],...
	'String','',...
	'BackgroundColor','w',...
	'Value',type_value,...
	'KeyPressFcn',@keypressmisc,...
	'Callback',@filter_type_callback,...
	'Visible','off');
if ~isempty(prop.type)
	filter_ed.String = prop.type{type_value,1};
	setfilspec()
	type_popup.String = prop.type(:,2);
	type_popup.Visible = 'on';
end
viewfullpath = uicontrol('Style','checkbox',...
	'Position',[380 335 230 20],...
	'String','Show full paths',...
	'Value',show_full_path,...
	'HorizontalAlignment','left',...
	'BackgroundColor',fig_color,...
	'KeyPressFcn',@keypressmisc,...
	'Callback',@showfullpath);
remove_dupes = uicontrol('Style','checkbox',...
	'Position',[380 360 280 20],...
	'String','Remove duplicates (as per full path)',...
	'Value',nodupes,...
	'HorizontalAlignment','left',...
	'BackgroundColor',fig_color,...
	'KeyPressFcn',@keypressmisc,...
	'Callback',@removedupes);
recall_button = uicontrol('Style','pushbutton',...
	'Position',[665 335 65 30],...
	'String','Recall',...
	'KeyPressFcn',@keypressmisc,...
	'Callback',@recall,...
	'ToolTip','Add previously selected items');
label4 = uicontrol('Style','text',...
	'Position',[380 405 350 20],...
	'String','Selected Items',...
	'HorizontalAlignment','center',...
	'BackgroundColor',fig_color);
done_button = uicontrol('Style','pushbutton',...
	'Position',[280 80 80 30],...
	'String','Done',...
	'KeyPressFcn',@keypressmisc,...
	'Callback',@done);
cancel_button = uicontrol('Style','pushbutton',...
	'Position',[280 30 80 30],...
	'String','Cancel',...
	'KeyPressFcn',@keypressmisc,...
	'Callback',@cancel);
% If necessary, add warning about number of items to be selected.
num_files_warn = uicontrol('Style','text',...
	'Position',[380 385 350 16],...
	'String',numstr,...
	'ForegroundColor',[0.8 0 0],...
	'BackgroundColor',fig_color,...
	'HorizontalAlignment','center',...
	'Visible','off');
if ~isempty(prop.numfiles)
	num_files_warn.Visible = 'on';
end
resize()
% Make figure visible and hide handle.
set(fig,'HandleVisibility','off',...
	'Visible','on',...
	'ResizeFcn',@resize)
% Wait until figure is closed.
uiwait(fig)
% Compute desired output.
switch prop.output
	case 'cell'
		out = full_file_picks;
	case 'struct'
		out = dir_picks(:);
	case 'char'
		out = char(full_file_picks);
	case 'cancel'
		out = 0;
end
% Update history preference.
setpref('uipickfiles','history',history)
if ~isempty(full_file_picks) && ~strcmp(prop.output,'cancel')
	setpref('uipickfiles','full_file_picks',full_file_picks)
end
% Update favorites preference.
setpref('uipickfiles','favorites',favorites)
% Update figure position preference.
setpref('uipickfiles','figure_position',fig_pos)
% ----------------- Callback nested functions ----------------
	function add(varargin)
		values = navlist.Value;
		for i = 1:length(values)
			dir_pick = fdir(values(i));
			pick = dir_pick.name;
			pick_full = fullfile(current_dir,pick);
			dir_pick.name = pick_full;
			if ~nodupes || ~any(strcmp(full_file_picks,pick_full))
				file_picks{end + 1} = pick; %#ok<AGROW>
				full_file_picks{end + 1} = pick_full; %#ok<AGROW>
				dir_picks(end + 1) = dir_pick; %#ok<AGROW>
			end
		end
		% Added 3/2/14 - 2 lines
		history = update_history(history,current_dir,now,history_size);
		make_history_cm(csdata)
		if show_full_path
			set(pickslist,'String',full_file_picks,'Value',[]);
		else
			set(pickslist,'String',file_picks,'Value',[]);
		end
		set([removebut,moveupbut,movedownbut],'Enable','off');
	end
	function add_with_subfolders(varargin)
		% Use final argument to determine exclusions for subdirs.
		switch varargin{end}
			case 'pathsub'
				exclusions = '.@+p';
			case 'sub'
				exclusions = '.';
			case 'allsub'
				exclusions = '';
		end
		values = navlist.Value;
		for i = 1:length(values)
			dir_pick = fdir(values(i));
			pick = dir_pick.name;
			pick_full = fullfile(current_dir,pick);
			subd = subdirs(pick_full,[],exclusions);
			for j = 1:length(subd)
				pick_full = subd{j};
				if ~nodupes || ~any(strcmp(full_file_picks,pick_full))
					[~,pick] = fileparts(pick_full);
					dir_temp = dir(pick_full);
					dir_pick = dir_temp(strcmp({dir_temp.name},'.'));
					dir_pick.name = pick_full;
					file_picks{end + 1} = pick; %#ok<AGROW>
					full_file_picks{end + 1} = pick_full; %#ok<AGROW>
					dir_picks(end + 1) = dir_pick; %#ok<AGROW>
				end
			end
		end
		% Added 3/2/14 - 2 lines
		history = update_history(history,current_dir,now,history_size);
		make_history_cm(csdata)
		if show_full_path
			set(pickslist,'String',full_file_picks,'Value',[]);
		else
			set(pickslist,'String',file_picks,'Value',[]);
		end
		set([removebut,moveupbut,movedownbut],'Enable','off');
	end
	function remove(varargin)
		values = pickslist.Value;
		file_picks(values) = [];
		full_file_picks(values) = [];
		dir_picks(values) = [];
		top = pickslist.ListboxTop;
		num_above_top = sum(values < top);
		top = top - num_above_top;
		num_picks = length(file_picks);
		new_value = min(min(values) - num_above_top,num_picks);
		if num_picks == 0
			new_value = [];
			set([removebut,moveupbut,movedownbut],'Enable','off')
		end
		if show_full_path
			set(pickslist,'String',full_file_picks,'Value',new_value,...
				'ListboxTop',top)
		else
			set(pickslist,'String',file_picks,'Value',new_value,...
				'ListboxTop',top)
		end
	end
	function open(varargin)
		values = navlist.Value;
		if fdir(values).isdir
			fig.Pointer = 'watch';
			drawnow
			% Convert 'My Documents' to 'Documents' when necessary.
			if ispc && strcmp(fdir(values).name,'My Documents')
				if isempty(dir(fullfile(current_dir,fdir(values).name)))
					values = find(strcmp({fdir.name},'Documents'));
				end
			end
			current_dir = fullfile(current_dir,fdir(values).name);
			if ispc && ~isempty(regexpi(current_dir,'\.lnk')) && ...
					is_shortcut_to_dir(current_dir)
				JFile = java.io.File(current_dir);
				sf = sun.awt.shell.ShellFolder.getShellFolder(JFile);
				current_dir = char(sf.getLinkLocation());
			end
% 			history = update_history(history,current_dir,now,history_size);
% 			make_history_cm(csdata)
			full_filter = fullfile(current_dir,filter);
			path_cell = path2cell(current_dir);
			fdir = filtered_dir(full_filter,re_filter,prop.redirs,...
				@(x,c)file_sort(x,sort_state,c));
			filenames = {fdir.name}';
			[filenames,fdir] = annotate_file_names(filenames,fdir,fsdata);
			set(dir_popup,'String',path_cell,'Value',length(path_cell))
			if length(path_cell) > 1
				up_dir_but.Enable = 'on';
			else
				up_dir_but.Enable = 'off';
			end
			pathbox.String = current_dir;
			set(navlist,'ListboxTop',1,'Value',[],'String',filenames)
			addbut.Enable = 'off';
			openbut.Enable = 'off';
			fig.Pointer = 'arrow';
		end
	end
	function clicknav(varargin)
		value = navlist.Value;
		nval = length(value);
		dbl_click_fcn = @add;
		switch nval
			case 0
				set([addbut,openbut],'Enable','off')
			case 1
				addbut.Enable = 'on';
				if fdir(value).isdir
					openbut.Enable = 'on';
					dbl_click_fcn = @open;
				else
					openbut.Enable = 'off';
				end
			otherwise
				addbut.Enable = 'on';
				openbut.Enable = 'off';
		end
		if any([fdir(value).isdir])
			set(add_cm_items,'Visible','on')
		else
			set(add_cm_items,'Visible','off')
		end
		if strcmp(fig.SelectionType,'open')
			dbl_click_fcn();
		end
	end
	function keypressmisc(h,evt) %#ok<INUSL>
		if strcmp(evt.Key,'escape') && isequal(evt.Modifier,cell(1,0))
			% Escape key means Cancel.
			cancel()
		end
	end
	function keypressnav(h,evt) %#ok<INUSL>
		if length(path_cell) > 1 && strcmp(evt.Key,'backspace') && ...
				isequal(evt.Modifier,cell(1,0))
			% Backspace means go to parent folder.
			dir_up_one()
		elseif strcmp(evt.Key,'f') && isequal(evt.Modifier,{mod_key})
			% Control-F (Command-F on Mac) means select all files.
			value = find(~[fdir.isdir]);
			navlist.Value = value;
		elseif strcmp(evt.Key,'rightarrow') && ...
				isequal(evt.Modifier,cell(1,0))
			% Right arrow key means select the file.
			add()
		elseif strcmp(evt.Key,'escape') && isequal(evt.Modifier,cell(1,0))
			% Escape key means Cancel.
			cancel()
		end
	end
	function keypresslist(h,evt) %#ok<INUSL>
		if strcmp(evt.Key,'backspace') && isequal(evt.Modifier,cell(1,0))
			% Backspace means remove item from list.
			remove()
		elseif strcmp(evt.Key,'escape') && isequal(evt.Modifier,cell(1,0))
			% Escape key means Cancel.
			cancel()
		end
	end
	function clickpicks(varargin)
		value = pickslist.Value;
		if isempty(value)
			set([removebut,moveupbut,movedownbut],'Enable','off')
		else
			removebut.Enable = 'on';
			if min(value) == 1
				moveupbut.Enable = 'off';
			else
				moveupbut.Enable = 'on';
			end
			if max(value) == length(file_picks)
				movedownbut.Enable = 'off';
			else
				movedownbut.Enable = 'on';
			end
		end
		if strcmp(fig.SelectionType,'open')
			remove();
		end
	end
	function recall(varargin)
		if ispref('uipickfiles','full_file_picks')
			ffp = getpref('uipickfiles','full_file_picks');
		else
			ffp = {};
		end
		for i = 1:length(ffp)
			if fdexist(ffp{i},'dir') && ...
					(~nodupes || ~any(strcmp(full_file_picks,ffp{i})))
				full_file_picks{end + 1} = ffp{i}; %#ok<AGROW>
				[unused,fn,ext] = fileparts(ffp{i});
				file_picks{end + 1} = [fn,ext]; %#ok<AGROW>
				path_c_temp = path2cell(ffp{i});
				temp = dir(cell2path(path_c_temp(1:end-1)));
				if ispc || ismac
					thisdir = strcmpi({temp.name},[fn,ext]);
				else
					thisdir = strcmp({temp.name},[fn,ext]);
				end
				dir_picks(end + 1) = temp(thisdir); %#ok<AGROW>
				dir_picks(end).name = ffp{i};
			elseif fdexist(ffp{i},'file') && ...
					(~nodupes || ~any(strcmp(full_file_picks,ffp{i})))
				full_file_picks{end + 1} = ffp{i}; %#ok<AGROW>
				[unused,fn,ext] = fileparts(ffp{i});
				file_picks{end + 1} = [fn,ext]; %#ok<AGROW>
				dir_picks(end + 1) = dir(ffp{i}); %#ok<AGROW>
				dir_picks(end).name = ffp{i};
			end
		end
		if show_full_path
			set(pickslist,'String',full_file_picks,'Value',[]);
		else
			set(pickslist,'String',file_picks,'Value',[]);
		end
		set([removebut,moveupbut,movedownbut],'Enable','off');
	end
	function sort_type(h,evt,cb) %#ok<INUSL>
		if sort_state(cb)
			sort_state(cb) = -sort_state(cb);
			last_sort_state(cb) = sort_state(cb);
		else
			sort_state = zeros(1,3);
			sort_state(cb) = last_sort_state(cb);
		end
		set(sort_cb,{'CData'},tri_icon(sort_state + 2)')
		
		fdir = filtered_dir(full_filter,re_filter,prop.redirs,...
				@(x,c)file_sort(x,sort_state,c));
		filenames = {fdir.name}';
		[filenames,fdir] = annotate_file_names(filenames,fdir,fsdata);
		set(dir_popup,'String',path_cell,'Value',length(path_cell))
		if length(path_cell) > 1
			up_dir_but.Enable = 'on';
		else
			up_dir_but.Enable = 'off';
		end
		pathbox.String = current_dir;
		set(navlist,'String',filenames,'Value',[])
		addbut.Enable = 'off';
		openbut.Enable = 'off';
		fig.Pointer = 'arrow';
	end
	function dirpopup(varargin)
		value = dir_popup.Value;
		container = path_cell{min(value + 1,length(path_cell))};
		path_cell = path_cell(1:value);
		fig.Pointer = 'watch';
		drawnow
		if ispc && value == 1
			current_dir = '';
			full_filter = filter;
			drives = getdrives(network_volumes);
			num_drives = length(drives);
			temp = tempname;
			mkdir(temp)
			dir_temp = dir(temp);
			rmdir(temp)
			fdir = repmat(dir_temp(1),num_drives,1);
			[fdir.name] = deal(drives{:});
		else
			current_dir = cell2path(path_cell);
% 			history = update_history(history,current_dir,now,history_size);
% 			make_history_cm(csdata)
			full_filter = fullfile(current_dir,filter);
			fdir = filtered_dir(full_filter,re_filter,prop.redirs,...
				@(x,c)file_sort(x,sort_state,c));
		end
		filenames = {fdir.name}';
		selected = find(strcmp(filenames,container));
		[filenames,fdir] = annotate_file_names(filenames,fdir,fsdata);
		set(dir_popup,'String',path_cell,'Value',length(path_cell))
		if length(path_cell) > 1
			up_dir_but.Enable = 'on';
		else
			up_dir_but.Enable = 'off';
		end
		pathbox.String = current_dir;
		set(navlist,'String',filenames,'Value',selected)
		addbut.Enable = 'off';
		fig.Pointer = 'arrow';
	end
	function dir_up_one(varargin)
		value = length(path_cell) - 1;
		container = path_cell{value + 1};
		path_cell = path_cell(1:value);
		fig.Pointer = 'watch';
		drawnow
		if ispc && value == 1
			current_dir = '';
			full_filter = filter;
			drives = getdrives(network_volumes);
			num_drives = length(drives);
			temp = tempname;
			mkdir(temp)
			dir_temp = dir(temp);
			rmdir(temp)
			fdir = repmat(dir_temp(1),num_drives,1);
			[fdir.name] = deal(drives{:});
		else
			current_dir = cell2path(path_cell);
% 			history = update_history(history,current_dir,now,history_size);
% 			make_history_cm(csdata)
			full_filter = fullfile(current_dir,filter);
			fdir = filtered_dir(full_filter,re_filter,prop.redirs,...
				@(x,c)file_sort(x,sort_state,c));
		end
		filenames = {fdir.name}';
		selected = find(strcmp(filenames,container));
		[filenames,fdir] = annotate_file_names(filenames,fdir,fsdata);
		set(dir_popup,'String',path_cell,'Value',length(path_cell))
		if length(path_cell) > 1
			up_dir_but.Enable = 'on';
		else
			up_dir_but.Enable = 'off';
		end
		pathbox.String = current_dir;
		set(navlist,'String',filenames,'Value',selected)
		addbut.Enable = 'off';
		fig.Pointer = 'arrow';
	end
	function change_path(varargin)
		fig.Pointer = 'watch';
		drawnow
		proposed_path = pathbox.String;
		% Process any folders named '..'.
		proposed_path_cell = path2cell(proposed_path);
		ddots = strcmp(proposed_path_cell,'..');
		ddots(find(ddots) - 1) = true;
		proposed_path_cell(ddots) = [];
		proposed_path = cell2path(proposed_path_cell);
		% Check for existance of folder.
		if ~fdexist(proposed_path,'dir')
			fig.Pointer = 'arrow';
			uiwait(errordlg(['Folder "',proposed_path,...
				'" does not exist.'],'','modal'))
			return
		end
		current_dir = proposed_path;
% 		history = update_history(history,current_dir,now,history_size);
% 		make_history_cm(csdata)
		full_filter = fullfile(current_dir,filter);
		[path_cell,new_network_vol] = path2cell(current_dir);
		if fdexist(new_network_vol,'dir')
			network_volumes = unique([network_volumes,{new_network_vol}]);
		end
		fdir = filtered_dir(full_filter,re_filter,prop.redirs,...
				@(x,c)file_sort(x,sort_state,c));
		filenames = {fdir.name}';
		[filenames,fdir] = annotate_file_names(filenames,fdir,fsdata);
		set(dir_popup,'String',path_cell,'Value',length(path_cell))
		if length(path_cell) > 1
			up_dir_but.Enable = 'on';
		else
			up_dir_but.Enable = 'off';
		end
		pathbox.String = current_dir;
		set(navlist,'String',filenames,'Value',[])
		addbut.Enable = 'off';
		openbut.Enable = 'off';
		fig.Pointer = 'arrow';
	end
	function showfullpath(varargin)
		show_full_path = viewfullpath.Value;
		if show_full_path
			pickslist.String = full_file_picks;
		else
			pickslist.String = file_picks;
		end
	end
	function removedupes(varargin)
		nodupes = remove_dupes.Value;
		if nodupes
			num_picks = length(full_file_picks);
			[unused,rev_order] = unique(full_file_picks(end:-1:1)); %#ok<SETNU>
			order = sort(num_picks + 1 - rev_order);
			full_file_picks = full_file_picks(order);
			file_picks = file_picks(order);
			dir_picks = dir_picks(order);
			if show_full_path
				set(pickslist,'String',full_file_picks,'Value',[])
			else
				set(pickslist,'String',file_picks,'Value',[])
			end
			set([removebut,moveupbut,movedownbut],'Enable','off')
		end
	end
	function moveup(varargin)
		value = pickslist.Value;
		removebut.Enable = 'on';
		n = length(file_picks);
		omega = 1:n;
		index = zeros(1,n);
		index(value - 1) = omega(value);
		index(setdiff(omega,value - 1)) = omega(setdiff(omega,value));
		file_picks = file_picks(index);
		full_file_picks = full_file_picks(index);
		dir_picks = dir_picks(index);
		value = value - 1;
		if show_full_path
			set(pickslist,'String',full_file_picks,'Value',value)
		else
			set(pickslist,'String',file_picks,'Value',value)
		end
		if min(value) == 1
			moveupbut.Enable = 'off';
		end
		movedownbut.Enable = 'on';
	end
	function movedown(varargin)
		value = pickslist.Value;
		removebut.Enable = 'on';
		n = length(file_picks);
		omega = 1:n;
		index = zeros(1,n);
		index(value + 1) = omega(value);
		index(setdiff(omega,value + 1)) = omega(setdiff(omega,value));
		file_picks = file_picks(index);
		full_file_picks = full_file_picks(index);
		dir_picks = dir_picks(index);
		value = value + 1;
		if show_full_path
			set(pickslist,'String',full_file_picks,'Value',value)
		else
			set(pickslist,'String',file_picks,'Value',value)
		end
		if max(value) == n
			movedownbut.Enable = 'off';
		end
		moveupbut.Enable = 'on';
	end
	function togglefilter(varargin)
		fig.Pointer = 'watch';
		drawnow
		value = showallfiles.Value;
		if value
			filter = '*';
			re_filter = '';
			set([filter_ed,refilter_ed],'Enable','off')
		else
			filter = filter_ed.String;
			re_filter = refilter_ed.String;
			set([filter_ed,refilter_ed],'Enable','on')
		end
		full_filter = fullfile(current_dir,filter);
		fdir = filtered_dir(full_filter,re_filter,prop.redirs,...
				@(x,c)file_sort(x,sort_state,c));
		filenames = {fdir.name}';
		[filenames,fdir] = annotate_file_names(filenames,fdir,fsdata);
		set(navlist,'String',filenames,'Value',[])
		addbut.Enable = 'off';
		fig.Pointer = 'arrow';
	end
	function toggle_refiltdirs(varargin)
		fig.Pointer = 'watch';
		drawnow
		value = refilterdirs.Value;
		prop.redirs = value;
		full_filter = fullfile(current_dir,filter);
		fdir = filtered_dir(full_filter,re_filter,prop.redirs,...
				@(x,c)file_sort(x,sort_state,c));
		filenames = {fdir.name}';
		[filenames,fdir] = annotate_file_names(filenames,fdir,fsdata);
		set(navlist,'String',filenames,'Value',[])
		addbut.Enable = 'off';
		fig.Pointer = 'arrow';
	end
	function setfilspec(varargin)
		fig.Pointer = 'watch';
		drawnow
		filter = filter_ed.String;
		if isempty(filter)
			filter = '*';
			filter_ed.String = filter;
		end
		% Process file spec if a subdirectory was included.
		[p,f,e] = fileparts(filter);
		if ~isempty(p)
			newpath = fullfile(current_dir,p,'');
			pathbox.String = newpath;
			filter = [f,e];
			if isempty(filter)
				filter = '*';
			end
			filter_ed.String = filter;
			change_path();
		end
		full_filter = fullfile(current_dir,filter);
		fdir = filtered_dir(full_filter,re_filter,prop.redirs,...
				@(x,c)file_sort(x,sort_state,c));
		filenames = {fdir.name}';
		[filenames,fdir] = annotate_file_names(filenames,fdir,fsdata);
		set(navlist,'String',filenames,'Value',[])
		addbut.Enable = 'off';
		fig.Pointer = 'arrow';
	end
	function setrefilter(varargin)
		fig.Pointer = 'watch';
		drawnow
		re_filter = refilter_ed.String;
		fdir = filtered_dir(full_filter,re_filter,prop.redirs,...
				@(x,c)file_sort(x,sort_state,c));
		filenames = {fdir.name}';
		[filenames,fdir] = annotate_file_names(filenames,fdir,fsdata);
		set(navlist,'String',filenames,'Value',[])
		addbut.Enable = 'off';
		fig.Pointer = 'arrow';
	end
	function filter_type_callback(varargin)
		type_value = type_popup.Value;
		set(filter_ed,'String',prop.type{type_value,1})
		setfilspec()
	end
	function done(varargin)
		% Optional shortcut: click on a file and press 'Done'.
% 		if isempty(full_file_picks) && strcmp(addbut.Enable,'on')
% 			add();
% 		end
		numfiles = length(full_file_picks);
		if ~isempty(prop.numfiles)
			if numfiles < prop.numfiles(1)
				msg = {'Too few items selected.',numstr};
				uiwait(errordlg(msg,'','modal'))
				return
			elseif numfiles > prop.numfiles(end)
				msg = {'Too many items selected.',numstr};
				uiwait(errordlg(msg,'','modal'))
				return
			end
		end
		fig_pos = fig.Position;
		delete(fig)
	end
	function cancel(varargin)
		prop.output = 'cancel';
		fig_pos = fig.Position;
		delete(fig)
	end
	function history_cb(h,evt,arg) %#ok<INUSL>
		fig.Pointer = 'watch';
		drawnow
		if ischar(arg)
			current_dir = arg;
		else
			current_dir = history(arg).name;
		end
% 		history = update_history(history,current_dir,now,history_size);
% 		make_history_cm(csdata)
		full_filter = fullfile(current_dir,filter);
		path_cell = path2cell(current_dir);
		fdir = filtered_dir(full_filter,re_filter,prop.redirs,...
				@(x,c)file_sort(x,sort_state,c));
		filenames = {fdir.name}';
		[filenames,fdir] = annotate_file_names(filenames,fdir,fsdata);
		set(dir_popup,'String',path_cell,'Value',length(path_cell))
		if length(path_cell) > 1
			up_dir_but.Enable = 'on';
		else
			up_dir_but.Enable = 'off';
		end
		pathbox.String = current_dir;
		set(navlist,'ListboxTop',1,'Value',[],'String',filenames)
		addbut.Enable = 'off';
		openbut.Enable = 'off';
		fig.Pointer = 'arrow';
	end
	function add_to_favorites(varargin)
		favorites{end+1} = current_dir;
		make_history_cm(csdata)
	end
	function remove_from_favorites(varargin)
		favorites = setdiff(favorites,current_dir);
		make_history_cm(csdata)
	end
	function clear_history(varargin)
		history = update_history(history(1),'',[],history_size);
		make_history_cm(csdata)
	end
	function set_history_size(varargin)
		result_cell = inputdlg('Number of Recent Folders:','',1,...
			{sprintf('%g',history_size)});
		if isempty(result_cell)
			return
		end
		result = sscanf(result_cell{1},'%f');
		if isempty(result) || result < 1
			return
		end
		history_size = result;
		history = update_history(history,'',[],history_size);
		make_history_cm(csdata)
		setpref('uipickfiles','history_size',history_size)
	end
	function resize(varargin)
		% Get current figure size.
		P = 'Position';
		pos = fig.Position;
		w = pos(3); % figure width in pixels
		h = pos(4); % figure height in pixels
		
		% Enforce minimum figure size.
		w = max(w,564);
		h = max(h,443);
		if any(pos(3:4) < [w h])
			pos(3:4) = [w h];
			fig.Position = pos;
		end
		
		% Change positions of all uicontrols based on the current figure
		% width and height.
		navw_pckw = round([1 1;-350 250]\[w-140;0]);
		navw = navw_pckw(1);
		pckw = navw_pckw(2);
		navp = [10 10 navw h-174];
		pckp = [w-10-pckw 10 pckw h-174];
		set(navlist,P,navp)
		set(pickslist,P,pckp)
		
		set(frame1,P,[navw+5 h-234 110 70])
		set(openbut,P,[navw+17 h-199 90 30])
		set(addbut,P,[navw+17 h-229 90 30])
		
		frame2y = round((h-234 + 110 - 100)/2);
		set(frame2,P,[w-pckw-115 frame2y 110 100])
		set(removebut,P,[w-pckw-105 frame2y+65 90 30])
		set(moveupbut,P,[w-pckw-105 frame2y+35 90 30])
		set(movedownbut,P,[w-pckw-105 frame2y+5 90 30])
		
		set(done_button,P,[navw+30 80 80 30])
		set(cancel_button,P,[navw+30 30 80 30])
		
		set(sort_cb(1),P,[15 h-163 70 15])
		set(sort_cb(2),P,[85 h-163 70 15])
		set(sort_cb(3),P,[155 h-163 70 15])
		
		set(dir_popup,P,[10 h-144 navw-25 20])
		set(up_dir_but,P,[navw-10 h-144 20 20])
		set(pathbox,P,[10 h-119 navw 26])
		set(label1,P,[10 h-93 navw 16])
		
		set(viewfullpath,P,[pckp(1) h-159 230 20])
		set(remove_dupes,P,[pckp(1) h-134 280 20])
		set(recall_button,P,[w-75 h-159 65 30])
		set(label4,P,[w-10-pckw h-89 pckw 20])
		set(num_files_warn,P,[w-10-pckw h-109 pckw 16])
		
		label2.Position(2) = h - 18;
		label3.Position(2) = h - 18;
		showallfiles.Position(2) = h - 42;
		refilterdirs.Position(2) = h - 64;
		filter_ed.Position(2) = h - 44;
		refilter_ed.Position(2) = h - 44;
		set(type_popup,P,[10 h-72 250 20])
	end
	function reset_figure_size(varargin)
		if strcmp(fig.SelectionType,'open')
			root_units = get(groot,'Units');
			screen_size = get(groot,'ScreenSize');
			set(0,'Units',root_units)
			hw = [740 494];
			pos = [round((screen_size(3:4) - hw - [0 26])/2),hw];
			fig.Position = pos;
			resize()
		end
	end
% ------------------ Other nested functions ------------------
	function make_history_cm(csdata)
		% Make context menu for history.
		if ~isempty(hist_menus)
			delete(hist_menus)
		end
		num_hist = length(history);
		hist_menus = zeros(1,num_hist);
		for i = 1:num_hist
			hist_menus(i) = uimenu(hist_cm,'Label',history(i).name,...
				'Callback',{@history_cb,i});
		end
		hist_menus(end+1) = uimenu(hist_cm,...
			'Label','Favorites',...
			'Enable','off',...
			'Separator','on');
		hist_menus(end+1) = uimenu(hist_cm,...
			'Label',[csdata.pre_home,csdata.home_folder,csdata.post],...
			'Callback',{@history_cb,csdata.home_folder});
		for i = 1:length(csdata.matlab_folders)
			hist_menus(end+1) = uimenu(hist_cm,...
				'Label',[csdata.pre_logo,csdata.matlab_folders{i},csdata.post],...
				'Callback',{@history_cb,csdata.matlab_folders{i}}); %#ok<AGROW>
		end
		for i = 1:length(favorites)
			hist_menus(end+1) = uimenu(hist_cm,...
				'Label',favorites{i},...
				'Callback',{@history_cb,favorites{i}}); %#ok<AGROW>
		end
		hist_menus(end+1) = uimenu(hist_cm,...
			'Label','Add Current Folder to Favorites',...
			'Separator','on',...
			'Callback',@add_to_favorites);
		hist_menus(end+1) = uimenu(hist_cm,...
			'Label','Remove Current Folder from Favorites',...
			'Callback',@remove_from_favorites);
		hist_menus(end+1) = uimenu(hist_cm,...
			'Label','Clear Menu',...
			'Separator','on',...
			'Callback',@clear_history);
		hist_menus(end+1) = uimenu(hist_cm,'Label',...
			sprintf('Set Number of Recent Folders (%d) ...',history_size),...
			'Callback',@set_history_size);
	end
end
% -------------------- Subfunctions --------------------
function [c,network_vol] = path2cell(p)
% Turns a path string into a cell array of path elements.
if ispc
	p = strrep(p,'/','\');
	c1 = regexp(p,'(^\\\\[^\\]+\\[^\\]+)|(^[A-Za-z]+:)|[^\\]+','match');
	vol = c1{1};
	c = [{'My Computer'};c1(:)];
	if strncmp(vol,'\\',2)
		network_vol = vol;
	else
		network_vol = '';
	end
else
	c = textscan(p,'%s','delimiter','/');
	c = [{filesep};c{1}(2:end)];
	network_vol = '';
end
end
% --------------------
function p = cell2path(c)
% Turns a cell array of path elements into a path string.
if ispc
	p = fullfile(c{2:end},'');
	if p(end) == ':'
		p = [p,filesep];
	end
else
	p = fullfile(c{:},'');
end
end
% --------------------
function d = filtered_dir(full_filter,re_filter,filter_both,sort_fcn)
% Like dir, but applies filters and sorting.
p = fileparts(full_filter);
if isempty(p) && full_filter(1) == '/'
	p = '/';
end
if fdexist(full_filter,'dir')
	dfiles = repmat(dir(char(127)),0,1);
else
	dfiles = dir(full_filter);
end
if ~isempty(dfiles)
	dfiles([dfiles.isdir]) = [];
end
ddir = dir(p);
ddir = ddir([ddir.isdir]);
[unused,index0] = sort(lower({ddir.name})); %#ok<ASGLU>
ddir = ddir(index0);
ddir(strcmp({ddir.name},'.') | strcmp({ddir.name},'..')) = [];
% Additional regular expression filter.
if nargin > 1 && ~isempty(re_filter)
	if ispc || ismac
		no_match = cellfun('isempty',regexpi({dfiles.name},re_filter));
	else
		no_match = cellfun('isempty',regexp({dfiles.name},re_filter));
	end
	dfiles(no_match) = [];
end
if filter_both
	if nargin > 1 && ~isempty(re_filter)
		if ispc || ismac
			no_match = cellfun('isempty',regexpi({ddir.name},re_filter));
		else
			no_match = cellfun('isempty',regexp({ddir.name},re_filter));
		end
		ddir(no_match) = [];
	end
end
% Set navigator style:
%	1 => list all folders before all files, case-insensitive sorting
%	2 => mix files and folders, case-insensitive sorting
%	3 => list all files before all folders, case-insensitive sorting
%	4 => list all folders before all files, case-sensitive sorting
%	5 => mix files and folders, case-sensitive sorting
%	6 => list all files before all folders, case-sensitive sorting
nav_style = 1;
switch nav_style
	case 1
		[unused,index1] = sort_fcn(dfiles,false); %#ok<ASGLU>
		[unused,index2] = sort_fcn(ddir,false); %#ok<ASGLU>
		d = [ddir(index2);dfiles(index1)];
	case 2
		d = [dfiles;ddir];
		[unused,index] = sort_fcn(d,false); %#ok<ASGLU>
		d = d(index);
	case 3
		[unused,index1] = sort_fcn(dfiles,false); %#ok<ASGLU>
		[unused,index2] = sort_fcn(ddir,false); %#ok<ASGLU>
		d = [dfiles(index1);ddir(index2)];
	case 4
		[unused,index1] = sort_fcn(dfiles,true); %#ok<ASGLU>
		[unused,index2] = sort_fcn(ddir,true); %#ok<ASGLU>
		d = [ddir(index2);dfiles(index1)];
	case 5
		d = [dfiles;ddir];
		[unused,index] = sort_fcn(d,true); %#ok<ASGLU>
		d = d(index);
	case 6
		[unused,index1] = sort_fcn(dfiles,true); %#ok<ASGLU>
		[unused,index2] = sort_fcn(ddir,true); %#ok<ASGLU>
		d = [dfiles(index1);ddir(index2)];
end
end
% --------------------
function [files_sorted,index] = file_sort(files,sort_state,casesen)
switch find(sort_state)
	case 1
		if casesen
			[files_sorted,index] = sort({files.name});
		else
			[files_sorted,index] = sort(lower({files.name}));
		end
		if sort_state(1) < 0
			files_sorted = files_sorted(end:-1:1);
			index = index(end:-1:1);
		end
	case 2
		if sort_state(2) > 0
			[files_sorted,index] = sort([files.datenum]);
		else
			[files_sorted,index] = sort([files.datenum],'descend');
		end
	case 3
		if sort_state(3) > 0
			[files_sorted,index] = sort([files.bytes]);
		else
			[files_sorted,index] = sort([files.bytes],'descend');
		end
end
end
% --------------------
function drives = getdrives(other_drives)
% Returns a cell array of drive names on Windows.
letters = char('A':'Z');
num_letters = length(letters);
drives = cell(1,num_letters);
for i = 1:num_letters
	if fdexist([letters(i),':\'],'dir')
		drives{i} = [letters(i),':'];
	end
end
drives(cellfun('isempty',drives)) = [];
if nargin > 0 && iscellstr(other_drives)
	drives = [drives,unique(other_drives)];
end
end
% --------------------
function [filenames,dir_listing] = ...
	annotate_file_names(filenames,dir_listing,fsdata)
% Adds a trailing filesep character to folder names and, optionally,
% prepends a folder icon or bullet symbol.
if ispc
	for i = 1:length(filenames)
		if ~isempty(regexpi(filenames{i},'\.lnk')) && ...
				is_shortcut_to_dir(dir_listing(i).name)
			filenames{i} = sprintf('%s%s%s%s',fsdata.pre_sc,filenames{i},...
				fsdata.filesep,fsdata.post);
			dir_listing(i).isdir = true;
		elseif dir_listing(i).isdir
			filenames{i} = sprintf('%s%s%s%s',fsdata.pre,filenames{i},...
				fsdata.filesep,fsdata.post);
		end
	end
else
	for i = 1:length(filenames)
		if dir_listing(i).isdir
			filenames{i} = sprintf('%s%s%s%s',fsdata.pre,filenames{i},...
				fsdata.filesep,fsdata.post);
		end
	end
end
end
% --------------------
function history = update_history(history,current_dir,time,history_size)
if ~isempty(current_dir)
	% Insert or move current_dir to the top of the history.
	% If current_dir already appears in the history list, delete it.
	match = strcmp({history.name},current_dir);
	history(match) = [];
	% Prepend history with (current_dir,time).
	history = [struct('name',current_dir,'time',time),history];
end
% Trim history to keep at most <history_size> newest entries.
history = history(1:min(history_size,end));
end
% --------------------
function success = generate_folder_icon(icon_path)
% Black = 1, manila color = 2, transparent white = 3.
im = [ ...
	3 3 3 1 1 1 1 3 3 3 3 3;
	3 3 1 2 2 2 2 1 3 3 3 3;
	3 1 1 1 1 1 1 1 1 1 1 3;
	1 2 2 2 2 2 2 2 2 2 2 1;
	1 2 2 2 2 2 2 2 2 2 2 1;
	1 2 2 2 2 2 2 2 2 2 2 1;
	1 2 2 2 2 2 2 2 2 2 2 1;
	1 2 2 2 2 2 2 2 2 2 2 1;
	1 2 2 2 2 2 2 2 2 2 2 1;
	1 1 1 1 1 1 1 1 1 1 1 1];
cmap = [0 0 0;255 220 130;255 255 255]/255;
fid = fopen(icon_path,'w');
if fid > 0
	fclose(fid);
	imwrite(im,cmap,icon_path,'Transparency',[1 1 0])
end
success = fdexist(icon_path,'file');
end
% --------------------
function success = generate_foldersc_icon(icon_path)
% Black = 1, blue color = 2, darker blue = 3, transparent white = 4.
im = [ ...
	4 4 4 1 1 1 1 4 4 4 4 4;
	4 4 1 2 2 2 2 1 4 4 4 4;
	4 1 1 1 1 1 1 1 1 1 1 4;
	1 2 2 2 2 2 3 2 2 2 2 1;
	1 2 2 2 2 2 2 1 2 2 2 1;
	1 2 2 2 1 1 1 1 1 2 2 1;
	1 2 2 1 2 2 2 1 2 2 2 1;
	1 2 1 2 2 2 3 2 2 2 2 1;
	1 2 2 2 2 2 2 2 2 2 2 1;
	1 1 1 1 1 1 1 1 1 1 1 1];
cmap = [0 0 0;163 185 255;65 83 128;255 255 255]/255;
fid = fopen(icon_path,'w');
if fid > 0
	fclose(fid);
	imwrite(im,cmap,icon_path,'Transparency',[1 1 1 0])
end
success = fdexist(icon_path,'file');
end
% --------------------
function success = generate_house_icon(icon_path)
im = [6 6 6 6 6 6 6 6 6 6 6 6 6 6 6 6;
	6 6 6 6 6 6 6 5 5 6 6 6 6 6 6 6;
	6 6 6 6 6 6 5 8 8 5 6 8 8 8 6 6;
	6 6 6 6 6 5 8 3 3 8 5 2 9 2 6 6;
	6 6 6 6 5 8 3 3 3 3 8 5 9 2 6 6;
	6 6 6 5 8 3 7 4 4 7 3 8 5 2 6 6;
	6 6 5 8 3 3 1 1 1 1 3 3 8 5 6 6;
	6 5 8 3 3 3 2 4 4 2 3 3 3 8 5 6;
	5 8 2 3 3 3 3 3 3 3 3 3 3 2 8 5;
	6 6 2 3 3 3 1 1 1 1 3 3 3 2 6 6;
	6 6 2 3 3 3 1 8 8 1 3 3 3 2 6 6;
	6 6 2 3 3 3 1 8 8 1 3 3 3 2 6 6;
	6 6 2 3 3 3 1 1 2 1 3 3 3 2 6 6;
	6 6 2 3 3 3 1 5 5 1 3 3 3 2 6 6;
	6 6 2 3 3 3 1 7 7 1 3 3 3 2 6 6;
	6 6 8 8 8 8 4 4 4 4 8 8 8 8 6 6];
im = [im,6*ones(16,5)];
cmap = [70 24 9;174 172 166;244 240 230;93 96 97;32 33 33;...
	255 255 255;153 71 21;66 52 39;255 255 254]/255;
fid = fopen(icon_path,'w');
if fid > 0
	fclose(fid);
	imwrite(im,cmap,icon_path,'Transparency',[1 1 1 1 1 0 1 1 1])
end
success = fdexist(icon_path,'file');
end
% --------------------
function success = generate_logo_icon(icon_path)
im = [9 9 9 9 9 9 9 9 9 9 9 9 9 9 9 9;
	9 9 9 9 9 9 9 9 9 9 10 9 9 9 9 9;
	9 9 9 9 9 9 9 9 9 10 8 6 9 9 9 9;
	9 9 9 9 9 9 9 9 9 4 3 7 10 9 9 9;
	9 9 9 9 9 9 9 9 2 1 7 7 6 9 9 9;
	9 9 9 9 9 9 9 2 1 1 7 7 3 9 9 9;
	9 9 9 9 9 10 4 1 1 8 7 7 3 5 9 9;
	9 9 9 9 10 1 1 1 1 8 7 7 3 6 5 9;
	9 9 10 2 4 4 1 1 8 8 7 7 3 3 9 9;
	10 2 2 2 4 4 1 1 8 3 3 7 7 3 6 9;
	9 2 2 4 4 4 1 8 8 3 7 7 7 3 6 5;
	9 9 10 2 4 1 8 8 8 3 7 7 6 6 3 5;
	9 9 9 9 9 1 8 3 3 7 7 9 9 9 6 6;
	9 9 9 9 9 5 3 7 7 7 5 9 9 9 9 5;
	9 9 9 9 9 9 6 7 7 5 9 9 9 9 9 9;
	9 9 9 9 9 9 5 6 5 9 9 9 9 9 9 9];
im = [im,9*ones(16,5)];
cmap = [73 50 49;132 193 188;182 60 15;97 146 141;246 224 205;...
	223 153 109;230 113 15;123 33 18;255 255 255;202 212 210]/255;
fid = fopen(icon_path,'w');
if fid > 0
	fclose(fid);
	imwrite(im,cmap,icon_path,'Transparency',[1 1 1 1 1 1 1 1 0 1])
end
success = fdexist(icon_path,'file');
end
% --------------------
function fsdata = set_folder_style(folder_style_pref)
% Set style to preference.
fsdata.style = folder_style_pref;
% If style = 1, check to make sure icon image file exists.  If it doesn't,
% try to create it.  If that fails set style = 2.
if fsdata.style == 1
	icon1_path = fullfile(prefdir,'uipickfiles_folder_icon.png');
	icon2_path = fullfile(prefdir,'uipickfiles_foldersc_icon.png');
	if ~(fdexist(icon1_path,'file') && fdexist(icon2_path,'file'))
		success1 = generate_folder_icon(icon1_path);
		success2 = generate_foldersc_icon(icon2_path);
		if ~(success1 && success2)
			fsdata.style = 2;
		end
	end
end
% Set pre and post fields.
if fsdata.style == 1
	icon1_url = ['file:///',strrep(strrep(icon1_path,':','|'),'\','/')];
	icon2_url = ['file:///',strrep(strrep(icon2_path,':','|'),'\','/')];
	fsdata.pre = sprintf('<html><img width=12 height=10 src="%s">&nbsp;',icon1_url);
	fsdata.pre_sc = sprintf('<html><img width=12 height=10 src="%s">&nbsp;',icon2_url);
	fsdata.post = '</html>';
elseif fsdata.style == 2
	fsdata.pre = '<html><b>&#8226;</b>&nbsp;';
	fsdata.pre_sc = '<html><b>&#8226;</b>&nbsp;';
	fsdata.post = '</html>';
elseif fsdata.style == 3
	fsdata.pre = '';
	fsdata.pre_sc = '';
	fsdata.post = '';
end
fsdata.filesep = filesep;
end
% --------------------
function csdata = set_cmenu_style(cmenu_style_pref)
% Set style to preference.
csdata.style = cmenu_style_pref;
% If style = 1, check to make sure icon image files exist.  If they don't,
% try to create them.  If that fails set style = 2.
if csdata.style == 1
	icon1_path = fullfile(prefdir,'uipickfiles_home_icon.png');
	icon2_path = fullfile(prefdir,'uipickfiles_logo_icon.png');
	if ~(fdexist(icon1_path,'file') && fdexist(icon2_path,'file'))
		success1 = generate_house_icon(icon1_path);
		success2 = generate_logo_icon(icon2_path);
		if ~(success1 && success2)
			csdata.style = 2;
		end
	end
end
% Set pre and post fields.
if csdata.style == 1
	icon1_url = ['file:///',strrep(strrep(icon1_path,':','|'),'\','/')];
	icon2_url = ['file:///',strrep(strrep(icon2_path,':','|'),'\','/')];
	csdata.pre_home = sprintf('<html><img width=21 height=16 src="%s">',icon1_url);
	csdata.pre_logo = sprintf('<html><img width=21 height=16 src="%s">',icon2_url);
	csdata.post = '</html>';
elseif csdata.style == 2
	csdata.pre_home = '';
	csdata.pre_logo = '';
	csdata.post = '';
end
% Get MATLAB folders from userpath.
matlab_folders = regexp(userpath,pathsep,'split');
matlab_folders(cellfun(@isempty,matlab_folders)) = [];
if ispc
	csdata.home_folder = getenv('USERPROFILE');
else
	csdata.home_folder = getenv('HOME');
end
csdata.matlab_folders = matlab_folders;
end
% --------------------
function prop = parsepropval(prop,varargin)
% Parse property/value pairs and return a structure.
properties = fieldnames(prop);
arg_index = 1;
while arg_index <= length(varargin)
	arg = varargin{arg_index};
	if ischar(arg)
		prop_index = match_property(arg,properties);
		prop.(properties{prop_index}) = varargin{arg_index + 1};
		arg_index = arg_index + 2;
	elseif isstruct(arg)
		arg_fn = fieldnames(arg);
		for i = 1:length(arg_fn)
			prop_index = match_property(arg_fn{i},properties);
			prop.(properties{prop_index}) = arg.(arg_fn{i});
		end
		arg_index = arg_index + 1;
	else
		error(['Properties must be specified by property/value pairs',...
			' or structures.'])
	end
end
end
% --------------------
function prop_index = match_property(arg,properties)
% Utility function for parsepropval.
prop_index = find(strcmpi(arg,properties));
if isempty(prop_index)
	prop_index = find(strncmpi(arg,properties,length(arg)));
end
if length(prop_index) ~= 1
	error('Property ''%s'' does not exist or is ambiguous.',arg)
end
end
% --------------------
function r = fdexist(item_path,type)
%fdexist: Check if file or directory exists.  Does not search MATLAB path.
%  type must be 'dir' or 'file'.
if strncmpi(type,'dir',length(type))
	r = java.io.File(item_path).isDirectory();
elseif strncmpi(type,'file',length(type))
	r = java.io.File(item_path).isFile();
end
end
% --------------------
function r = is_shortcut_to_dir(filename)
r = false;
% jFile = java.io.File(filename);
% sf = sun.awt.shell.ShellFolder.getShellFolder(jFile);
% r = sf.isDirectory();
end
% --------------------
function d = subdirs(basedir,depth,exclusions)
%subdirs: Recursive directory finder.
% Recursively find all subdirectories of a specified directory.
%
% Syntax:
%   dirs = subdirs;
%
% will return all subdirectories of the current directory, including the
% current directory, in a cell array of strings.
%
%   dirs = subdirs(base);
%
% starts at the directory in the string, base, rather than the current
% directory.
%
%   dirs = subdirs(base,depth);
%
% only searches to a limited depth (default is Inf).  A depth of zero
% returns only the base directory, depth = 1 returns the base directory and
% its immediate decendants, etc.
% Version: 1.1, 8 November 2014
% Author:  Douglas M. Schwarz
% Email:   dmschwarz=ieee*org, dmschwarz=urgrad*rochester*edu
% Real_email = regexprep(Email,{'=','*'},{'@','.'})
% If base directory not specified, use current directory.
if nargin < 1
	basedir = pwd;
end
% If depth not specified, search to infinite depth.
if nargin < 2 || isempty(depth)
	depth = inf;
end
if nargin < 3 || isempty(exclusions)
	exclusions = '';
end
% If instructed not to search deeper, return basedir in a cell.
if depth == 0
	d = {basedir};
	return
end
% Check exclusions for 'p' => exclude 'private' directories.  Other
% exclusions characters exclude directories beginning with that character,
% e.g., '.@+'.
exclu = exclusions;
no_private = any(exclu == 'p');
exclu(exclu == 'p') = [];
% Get directory contents.
items = dir(basedir);
% Get the name of each item in basedir.  Remove items that are not
% directories and the special directories named '.' and '..', leaving only
% the desired subdirectories.
item_names = {items.name};
item_names(~[items.isdir] | strcmp(item_names,'.') | ...
	strcmp(item_names,'..')) = [];
% Remove private directories, if desired.
if no_private
	item_names(strcmpi(item_names,'private')) = [];
end
% Remove directories beginning with the characters in exclu, e.g., '.@+'.
for i = 1:length(exclu)
	item_names(strncmp(item_names,exclu(i),1)) = [];
end
% Run this function recursively on each subdirectory and return basedir and
% those subdirectories.
num_items = length(item_names);
subitems = cell(1,num_items);
for i = 1:num_items
	subitems{i} = subdirs(fullfile(basedir,item_names{i},''),...
		depth - 1,exclusions);
end
d = [{basedir},subitems{:}];
end
```
