## Supplementary material for "AMPK Regulates Phagophore-to-Autophagosome Maturation": K-FOCUS Code: TrackColocalization_2CH.m

```
%TrackColocalization using TrackedPar data, written by David Broadbent in the Schmidt
%Lab- 11/08/2022

clear all
close all 

%Customize analysis by changing these values
ColocalizationThreshold = 3;  %Changes the threshold that call colocalized tracks
TimeColocalized = 10;

%Get Directory
directory = uipickfiles;

for iter = 1:length(directory)
 
%Get ParentDirectory
parts = strsplit(directory{iter}, '\');
parentdirectory = strjoin(parts(1:end-1), '\');
clear parts

%Generate list of movies in directory per channel
Ch0FilesList = dir(fullfile(directory{iter}, '*C1*'));
Ch1FilesList = dir(fullfile(directory{iter}, '*C2*'));

%initalize
Colocalized = struct;
tic
%For loop to load individual movies from list for analysis
for movieID = 1:length(Ch0FilesList)
data = load(strcat(Ch0FilesList(movieID).folder, '\', Ch0FilesList(movieID).name));
Ch0Tracks = data.data;
data = load(strcat(Ch1FilesList(movieID).folder, '\', Ch1FilesList(movieID).name));
Ch1Tracks = data.data;

%initialize variables and structures
C=[];
D=[];
Distn=[];
ColocalizedTrackID=[];
empty= struct('TrackID', [],'xy', [], 'stepsize',[],'frame',[], 'TimeStamp',[], 'movie',[], 'cellnumber', [], 'PeakIntensity', [], 'ROISize', [], 'minDistEdge', []);
colocstruct = struct('TrackID', [], 'xy', [], 'stepsize',[],'frame',[], 'TimeStamp',[], 'movie',[], 'cellnumber', [], 'PeakIntensity', [], 'ROISize', [], 'minDistEdge', []);

%print movieID
fprintf(1, 'Now reading movie %s\n', char(strcat(num2str(movieID),':',{' '},Ch1FilesList(movieID).name)));

%If tracks exist in both channels proceed, else assign [], this was added
%to deal with matlab which says that empty structures have a size of 1, but
%an empty array is called 0. 
E = [Ch1Tracks.movie];
F = [Ch0Tracks.movie];

if length(E)>0 && length(F)>0
     
%Assigns Tracks to A from CH1
for trackID = 1:length(Ch1Tracks)
A = [Ch1Tracks(trackID).xy, Ch1Tracks(trackID).frame];
for iter = 1:length(Ch0Tracks)
B=[Ch0Tracks(iter).xy, Ch0Tracks(iter).frame];

%finds overlapping index for A and B
FrameOverlapIndexA = ismember(A(:,3),B(:,3));
FrameOverlapIndexB = ismember(B(:,3),A(:,3));

%makes new matrix with only overlapping track frames
OverlapA = A(FrameOverlapIndexA,1:3);
OverlapB = B(FrameOverlapIndexB,1:3);

%if the overlapping frames is greater than 1, calculate mean difference
%accross frames.
if size(OverlapA,1)>1
Distn = [];
for iteriter = 1:size(OverlapA,1)
        %Distn(iteriter)=pdist2_subfun(OverlapA(iteriter,1:2),OverlapB(iteriter,1:2),'euclidean','Smallest',1); % calculate the Euclidean distance
        Distn(iteriter)=markerdistance(OverlapA(iteriter,1:2),OverlapB(iteriter,1:2)); % calculate the Euclidean distance 
end

idx1 = Distn<=ColocalizationThreshold;
colocidx(iter) = sum(idx1);
else
colocidx(iter) = 0;
end
end

if max(colocidx) > TimeColocalized
   [M,I] = max(colocidx);
   D(trackID) = I;
else
    D(trackID) = [NaN];
end

if ~isnan(D(trackID)) 
ColocalizedTrackID(trackID) = D(trackID);
colocstruct(trackID) = Ch0Tracks(D(trackID));

else 
ColocalizedTrackID(trackID) = NaN;
colocstruct(trackID) = struct('TrackID', NaN, 'xy', NaN, 'stepsize',NaN,'frame', NaN, 'TimeStamp', NaN, 'movie', NaN, 'cellnumber', NaN, 'PeakIntensity', NaN, 'ROISize', NaN, 'minDistEdge', NaN);
end

%clear variables for another iteration
C=[];
D=[];
colocidx=[];

end


ColocalizedTrackIDs{movieID} = ColocalizedTrackID;
ColocalizedEvents{movieID} = length(ColocalizedTrackIDs{movieID}(~isnan(ColocalizedTrackIDs{movieID})));
ColocalizedTracks{movieID} = ColocalizedTrackIDs{movieID}(~isnan(ColocalizedTrackIDs{movieID}));
CH1AllEvents{movieID} = length(Ch1Tracks);
CH0AllEvents{movieID} = length(Ch0Tracks);
elseif length(E) == 0 && length(F)>0
ColocalizedTrackIDs{movieID} = [];
ColocalizedEvents{movieID} = 0;
CH1AllEvents{movieID} = length(Ch1Tracks);
CH0AllEvents{movieID} = length(Ch0Tracks);
else
ColocalizedTrackIDs{movieID} = [];
ColocalizedEvents{movieID} = [];
end

%combine colocalized CH0 tracks with CH1 tracks, only runs if marker has
%any tracks
if length(F)>0
Colocalized(movieID).tracks = cell2struct([struct2cell(Ch1Tracks);struct2cell(colocstruct)],[fieldnames(Ch1Tracks);strcat('CH2',fieldnames(colocstruct))]);
end

%clear ColocalizedTrackIDs for another iteration
ColocalizedTrackID=[];
colocstruct = struct('TrackID', [],'xy', [], 'stepsize',[],'frame',[], 'TimeStamp',[], 'movie',[], 'cellnumber', [], 'PeakIntensity', [], 'ROISize', [], 'minDistEdge', []);
end

%delete empty track spaces
fun = @(s) all(structfun(@isempty,s)); % check the fields of a scalar structure.
idx2 = arrayfun(fun,Colocalized); % indices of those structure elements with ALL fields empty.
Colocalized(idx2)=[];

%make final structure to save
for i=1:size(Colocalized,2)
    dataStruct(i).TrackIDs = ColocalizedTrackIDs{i};
    dataStruct(i).Tracks = Colocalized(i).tracks;
    dataStruct(i).ColocalizedTracks = Colocalized(i).tracks(~isnan([Colocalized(i).tracks.CH2TrackID]));
    dataStruct(i).NonColocalizedTracks = Colocalized(i).tracks(isnan([Colocalized(i).tracks.CH2TrackID]));
    dataStruct(i).ColocalizedEvents = ColocalizedEvents{i};
    dataStruct(i).NonColocalizedEvents = length(dataStruct(i).NonColocalizedTracks);
    dataStruct(i).CH1AllEvents = CH1AllEvents{i};
    dataStruct(i).CH0AllEvents = CH0AllEvents{i};

%   clear AverageLengthColoc AverageLengthNonColoc AverageColocCH1PeakIntensity AverageColocCH2PeakIntensity AverageNonColocCH1PeakIntensity NonColoctemplength Coloctemplength
    NonColoctemplength=[];
    Coloctemplength=[];
    AverageNonColocCH1PeakIntensity=[];
    AverageColocCH2PeakIntensity=[];
    AverageColocCH1PeakIntensity=[];
    AverageLengthNonColoc=[];
    AverageLengthColoc=[];
    ColoctempCH1FociPeakIntensity=[];
    ColoctempCH2FociPeakIntensity=[];
    nonColoctempCH1FociPeakIntensity=[];


    %Calculate conversion rate, average track length, and average Intensity
    for ii=1:size(dataStruct(i).ColocalizedTracks,2)
                
    Coloctemplength(ii) = length(dataStruct(i).ColocalizedTracks(ii).xy);
    for iiii=1:size(dataStruct(i).ColocalizedTracks,2)
    ColoctempCH1FociPeakIntensity(iiii) = mean(dataStruct(i).ColocalizedTracks(iiii).PeakIntensity);
    ColoctempCH2FociPeakIntensity(iiii) = mean(dataStruct(i).ColocalizedTracks(iiii).CH2PeakIntensity);
    end
    for iiii=1:size([dataStruct(i).NonColocalizedTracks],2)
    nonColoctempCH1FociPeakIntensity(iiii) = mean(dataStruct(i).NonColocalizedTracks(iiii).PeakIntensity);
    end
    
    
    end
    
    for iii=1:size(dataStruct(i).NonColocalizedTracks,2)
    NonColoctemplength(iii) = length(dataStruct(i).NonColocalizedTracks(iii).xy);
    end
        dataStruct(i).ConversionRatio = (ColocalizedEvents{i}/CH1AllEvents{i});
        dataStruct(i).AverageLengthColoc = mean(Coloctemplength,"omitnan");
        dataStruct(i).AverageLengthNonColoc = mean(NonColoctemplength,"omitnan");
        dataStruct(i).AverageColocCH1PeakIntensity = mean(ColoctempCH1FociPeakIntensity,"omitnan");
        dataStruct(i).AverageColocCH2PeakIntensity = mean(ColoctempCH2FociPeakIntensity,"omitnan");
        dataStruct(i).AverageNonColocCH1PeakIntensity = mean(nonColoctempCH1FociPeakIntensity,"omitnan");
        
%         clear AverageLengthColoc AverageLengthNonColoc AverageColocCH1PeakIntensity AverageColocCH2PeakIntensity AverageNonColocCH1PeakIntensity NonColoctemplength
    end

        


title = strcat(directory{iter}, '_Colocalized.mat'); %title to save final structure
save(title, 'dataStruct','-mat');
clear dataStruct
end

toc
```
