## Supplementary material for "AMPK Regulates Phagophore-to-Autophagosome Maturation": K-FOCUS Code: README.txt

# K-FOCUS Quantitative Imaging Pipeline
Developed by Barnaba C, Broadbent D, & Schmidt J in July 2023

## Overview
The K-FOCUS quantitative imaging pipeline is a powerful tool designed for analyzing foci co-diffusion and kinetics in 2-color live-cell imaging at a single-cell resolution. The pipeline consists of three essential steps:

- **Cellular Segmentation using CellPose** (Stringer et al., Nature Methods 2021; Pachitariu & Stringer, Nature Methods 2022)
- **Single-cell Foci Tracking using TrackIt** (Kuhn et al., Scientific Reports 2021)
- **K-FOCUS Co-diffusion Analysis**

## Raw Image Naming
For K-FOCUS to operate effectively, it requires that each movie's two channels be saved as separate TIFF files with matching names, except for the channel attributes 'C0' and 'C1'. Additionally, include information about the nature of the experiment (e.g., control or treatment) in the filename.

## Cellular Segmentation
CellPose, a Python-based deep-learning algorithm, facilitates precise single-cell segmentation. Detailed installation procedures can be found in the corresponding papers. We've included a Python code called 'main' in the Cell Segmentation folder, which executes CellPose using a pretrained model. Our customized CellPose code generates a list of region-of-interests (ROIs) for each segmented cell in a MatLab compatible file format. You can then import these ROIs into TrackIt for foci tracking.

## Single Cell Foci Tracking
TrackIt, a MatLab-based GUI integrative tracking and analysis software developed by the Gebhardt group, streamlines single-cell foci tracking. You can import the ROI file containing cell segmentation for each movie into TrackIt. After manually verifying and correcting segmentation, you can execute the tracking algorithm and export the results as a batch file. The output is a MatLab structure that contains single-cell foci tracking data. Important note: Do not change the location of the raw images at this stage, as the batch file references their location.

## K-FOCUS Co-diffusion Analysis
K-FOCUS offers two MatLab scripts:

- **K-Focus_GUI**: This user-friendly GUI application simplifies batch processing. When executed, a GUI interface prompts you to specify the number of experimental conditions (e.g., 2 for control and treatment) and the minimum frame length (e.g., 5). You can manually add conditions and save settings in a file named 'KFocus_Settings'. K-FOCUS_GUI automates the following tasks:
   - Background intensity correction.
   - Calculation of foci intensity and other metrics.
   - Storage of single-cell analysis results in a folder named 'Kfocus_analysis,' which contains subfolders corresponding to each experiment (e.g., control and treatment).

- **TrackColocalization_2CH**: This script performs colocalization analysis. Upon execution, a GUI interface allows you to select the conditions to be analyzed. Simply choose (or drag & drop) the folder corresponding to the experiments (e.g., control and treatment). In 'TrackColocalization_2CH,' you can specify the following colocalization criteria:
   - ColocalizationThreshold
   - TimeColocalized 
   'TrackColocalization_2CH' outputs a MatLab structure containing the number of foci for each channel, colocalized foci, and the lifetime of C1-positive and C1-negative foci.

## Contact
For questions or inquiries, please feel free to contact Jens Schmidt.
